# Epigenetically distinct synaptic architecture in clonal compartments in the teleostean dorsal pallium

**DOI:** 10.1101/2022.10.01.510385

**Authors:** Yasuko Isoe, Ryohei Nakamura, Shigenori Nonaka, Yasuhiro Kamei, Teruhiro Okuyama, Naoyuki Yamamoto, Hideaki Takeuchi, Hiroyuki Takeda

## Abstract

The dorsal telencephalon (i.e. the pallium) exhibits high anatomical diversity across vertebrate classes. The mammalian dorsal pallium accommodates a six layered-structure, the neocortex, whereas the teleostean dorsal pallium possesses various compartmentalized structures among species. The development, function and evolution of the fish dorsal pallium remain unillustrated. Here, we analyzed the structure and epigenetic landscapes of cell lineages in the telencephalon of medaka fish (*Oryzias latipes*) which possesses a clearly delineated dorsal pallium (the Dd2 region). We found that different pallial regions, including Dd2, are formed by mutually exclusive clonal units, and that each pallium compartment exhibits a distinct epigenetic landscape. In particular, Dd2 possesses a unique open chromatin pattern that preferentially targets synapse-related genes. Indeed, Dd2 shows a high density of synapses, which might reflect strong plasticity. Finally, we identified several transcription factors as candidate regulators for the Dd2, which are partially shared with the human neocortex and hippocampus.

## Introduction

The telencephalon is an essential brain part for an animal’s cognitive functions and is highly diverse in structure among vertebrates. The dorsal telencephalon, or the pallium, is divided into several distinct regions in mammals; *e.g*. the cerebral neocortex in the mammalian dorsal pallium (MDP),the hippocampus in the mammalian medial pallium (MMP) and the basolateral amygdala in the mammalian lateral pallium (MLP) ^1^. For decades, the anatomical homologies of the pallial regions in vertebrates has been controversial, especially so for the MDP ^2, 3, 4^. The MDP is characterized by a six-layered structure and by stereotypical projections from all sensory modalities.

The non-mammalian dorsal pallium generally lacks a multilayered structure, with the exception of non-avian reptiles ^5, 6^, but rather exhibits a compartmentalized architecture which is highly diverse across species. Nonetheless, the neocortex in mammals, as well as the dorsal pallium in non-mammalian clades, receives structured input from all sensory modalities and these structures therefore are hypothesized to fulfill similar function. Somatosensory input into the dorsal pallium in marbled rockfish (^2,7^ and visual projections from the optic tectum in the yellowfin goby ^8,9,10^) are just a few specific examples of such projections in teleost species. However, because of the lack of genetic tools in these species, the molecular characterization of the dorsal pallium in teleosts is still largely underexplored. In zebrafish (*Danio rerio*), the most popular fish model organism, a clear dorsal pallium could not be delineated ^11,12^, which has made a detailed characterization difficult in this species.

To study how the specific architecture of the dorsal pallium in teleosts emerges and is maintained throughout life, we combined two orthogonal approaches.First,cell-lineage analysis was used to visualize the cellular subpopulations,or *clonal units*,that are derived from the same neural stem cells. Since post-hatch neurogenesis occurs throughout life in all teleosts, such cell lineage analysis of neural progenitors allows us to collectively label individual cell populations that may serve similar functions ^13–15^. Second, for characterizing gene expression within these clonal units, we focused on epigenetic regulation of transcription, where ATAC-sequencing ^16,17^ allows for the identification of specifically regulated genes within each population,whose expression levels are further quantified by subsequent RNA-sequencing.

Here,we selected medaka fish (*Oryzias latipes*) as a model organism for three reasons. First, medaka possesses a dorsal pallium (Dd2) that is demarcated by a cellular boundary which makes it anatomically distinct ^18^ and facilitates the isolation of this particular region. Second, transgenic medaka lines are available to visualize neural progenitors in the brain ^19^. Third, the genome size of medaka is smaller than that of zebrafish ^20^, the genome sequence is of high quality ^21^ and it is readily available for epigenetic analysis ^22,23^.

We first investigated the clonal architecture of the entire telencephalon in medaka, and found that all anatomical regions within the pallium, including Dd2, are formed by mutually exclusive clonal units. We next observed that clonal units in Dd2 in particular possess a unique open chromatin landscape, where synaptic-related genes were actively regulated. Subsequently, we examined the transcriptional regulatory mechanism in Dd2 to identify some candidate transcription factors (TFs) that contribute to the unique epigenetic landscape in the dorsal pallium. Finally, we performed a TF-binding motif-targeted comparison between medaka and humans, and we found that the regulatory elements between medaka’s Dd2 and the human neocortex and hippocampus exhibit higher similarity. Our data suggest that synaptic architecture in the dorsal pallium in medaka is constructed according to different rules from those of other pallial regions, and that it is regulated actively and carries a potential of a universal functionality within the telencephalic network across vertebrates.

## Results

### Anatomical regions in the adult medaka pallium consist of cell bodies of clonal units

Other than in mammals, in fish the brain grows continuously throughout life via post-hatch neurogenesis ^24^ (Figure 1A left), such that the anatomical classification framework needs to be adapted to this expansion. For example, the nomenclature of the adult medaka telencephalon has been complicated by the difficulties of identifying the small brain structures that emerge in adult brain sections throughout development ^18, 25^. Here, we first re-defined the anatomical regions in the medaka telencephalon based on the three dimensional distribution of cell nuclei by DAPI staining (Figure 1A right, Figure S1, Video S1). To acquire optical sections of the whole adult telencephalon, we applied a tissue clearing solution, Sca/eA2 ^26^, followed by light-sheet microscopic imaging. We also defined the anatomical regions by immunostaining of cell-type specific neural markers, such as CaMK2α, parvalbumin, and GAD65/67 (Table 1, Figure S2). Here we found clear anatomical boundaries in the dorsal pallium where different marker genes expressed along the anterior-posterior axis (Dd2a, 2p) (Figure 1A right, Figure S2. A detailed description of nomenclature is in Supplemental material).

**Figure 1.**
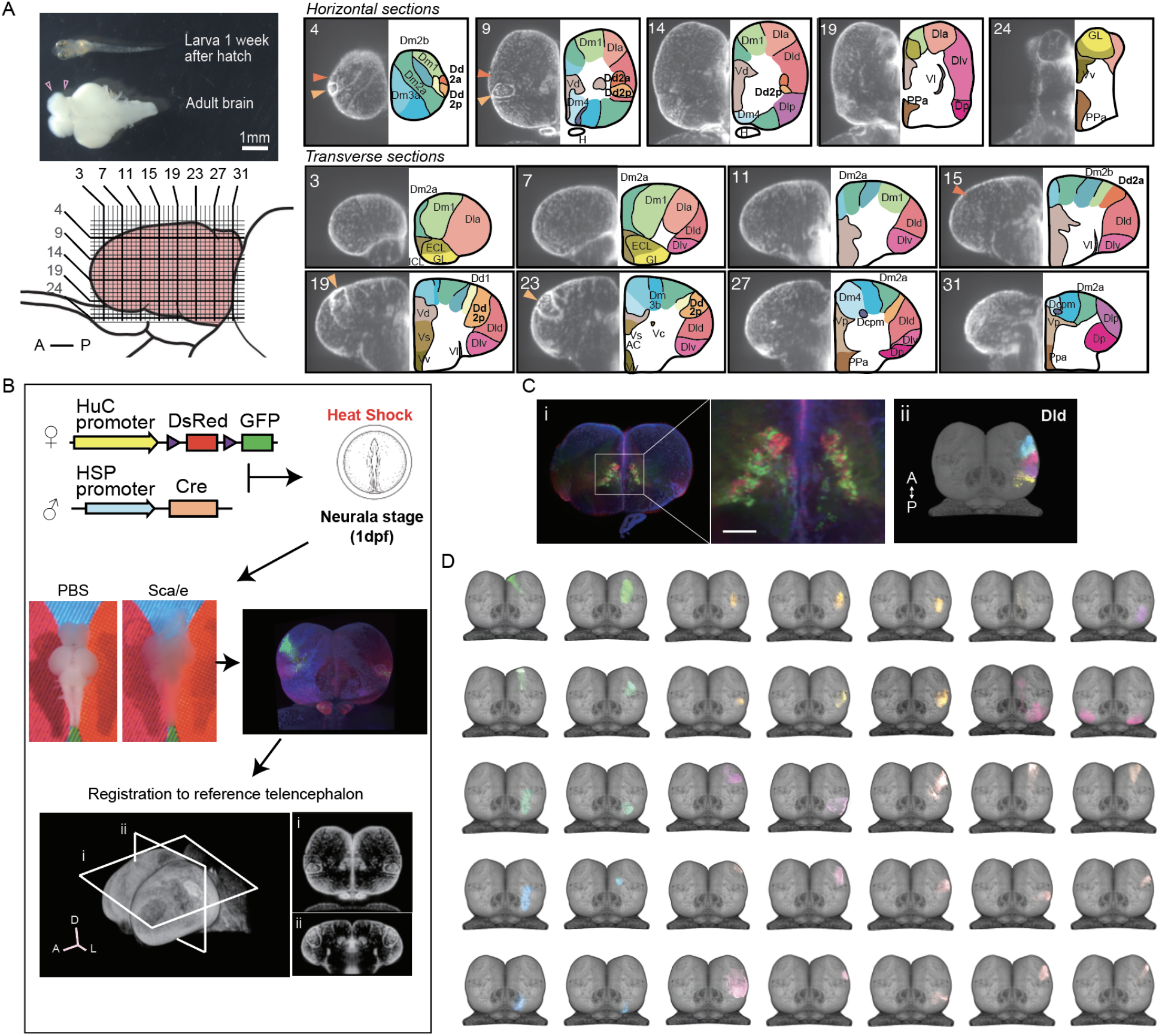
Clonal architecture in the adult medaka telencephalon. (A) A larval medaka fish (one week day post fertilization) and a dissecte dadult brain (top left). Pink triangles indicate the position of the telencephalon in the adult brain. Scale bar:1mm. Schematic drawing of the lateral view of the adult telencephalon (Pink, bottom left). Horizontal and vertical lines indicate the position of optical sections in the brain atlas on the right panel. Re-defined anatomical regions of adult medaka telencephalon are shown (right). Optical horizontal sections (top right) and transverse sections (bottom right). Orange triangles indicate the position of the dorsal pallial regions we focus on (dark orange: Dd2a, light orange: Dd2p). For each section, the left picture shows DAPI signals and the right shows the brain atlas. (B) Experimental procedure to label clonal units. Cre-loxp recombination was induced by a short heat shock at the neurula stage of transgenic embryos (Tg (HuC:loxp-DsRed-loxp-GFP) x Tg(HSP:Cre)). Dissected brains were stained with DAPI, cleared in Sca/e solution, and light-sheet microscopy images were taken.Fluorescent signals were registered to a reference telencephalon. Optical horizontal and transverse sections are shown (i, ii). A: anterior direction, D: dorsal direction, L: lateral direction. (C) Examples of anatomical regions consist of clonal units. Anatomical regions in the pallium were constituted in an exclusive mosaic way in the dorsal part of the lateral pallium (Dld) (i), while the subpallial regions were constituted with clonal units in a mixed way in Vd (ii). A: anterior, P: posterior direction. (D) Examples of the structure of clonal units identified in the telencephalon. Dorsal views are shown. Colors of clonal units indicate the position of cell somas in the anatomical region in (A). Detailed structure of clonal units are shown in Figure S3.

**Table 1.** Summary of marker gene expression in the telencephalic regions in medaka. -: signals not detected, +: signals detected, ++: signals strongly detected, //: not observed.

|  | CaMK2 | Parvalbumin | GAD65/67 | GAD67 |
| --- | --- | --- | --- | --- |
| GL | + | // | ++ | + |
| ICL | - | ++ | // | // |
| Dm1 | - | - | + | // |
| Dm2a | + | - | + | + |
| Dm2b | ++ | - | + | + |
| Dm3a | ++ | - | + | + |
| Dm3b | ++ | - | + | + |
| Dm4 | ++ | ++ | - | // |
| Dd1 | + | ++ | - | + |
| Dd2a | + | ++ | ++ | + |
| Dd2p | ++(ventral) | + | ++(dorsal) | + |
| Dla | + | // | // | // |
| Dld | +/- | + | + | ++ |
| Dlv | + | + | + | ++ |
| Dlp | +/- | - | - | // |
| Dc | + | - | - | // |
| Dcpm | - | - | // | // |
| Dcp | + |  | // | // |
| Dp | ++ | + | - | // |
| Vd | - | ++ | - | - |
| Vs | - | + | - | - |
| VI | - | ++ | - | - |
| Vv | - | ++ | - | - |
| Vp | ++ | + | - | - |

Next, to uncover the clonal architecture in the telencephalon, we genetically visualized the spatial distribution of cell lineages as previously established^19^ (Figure 1B). We used a transgenic line (Tg(HuC:loxp-DsRed-loxp-GFP)) which labels neural progenitors by the HuC promoter, and crossed it into a second line (Tg(HSP-Cre)) which expresses Cre recombinase under the heat shock protein promoter. The Cre-loxp recombination was induced stochastically by heat shock at the neurula stage (stage 16-17 ^27^) to visualize the progenitors derived from the same neural stem cells. We cleared the adult brains with Sca/e solution, stained them with DAPI and performed light-sheet microscope imaging. We first found that GFP-signal distributed differently between the pallium and subpallium; GFP-positive cell somas mixed with DsRed-positive cell somas in the subpallial regions (Figure 1C i), while GFP-positive cells in the pallium formed exclusive compartments (Figure 1B).

For further investigation of the clonal architecture in each anatomical region, we analyzed the structure of the clonal units by applying a normalization method ^28^ (Figure 1B). To that end, we registered the telencephalon of 81 medaka where 561 GFP-positive subpopulations were detected in total to a reference telencephalon. GFP-positive subpopulations that appeared multiple times in the same area were defined as clonal units (Figure 1D, Table 2, Figure S3, A detailed description of the clonal units is in Supplementary materials). For example, in the lateral pallium (Dl), the dorsal and ventral part of Dl were clearly different in terms of how the GFP-positive cell bodies are distributed (Figure S3). Specifically, in the dorsal pallium, the anterior part (Dd2a) and the posterior part (Dd2p) consists of two clonal units each which cluster laterally and medially respectively. Taken together, we found that the anatomical regions in the pallium are formed by mutually exclusive compartments of clonally-related cell bodies.

**Table 2.**
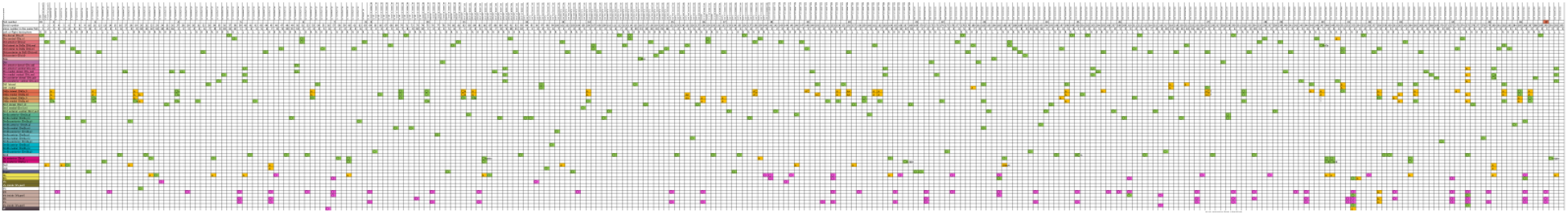
Identified clonal units in 81 medaka. 81 fish brain samples were analyzed. GFP-positive cell subpopulations in each sample were recorded. Green box shows the location of GFP-positive cell bodies. Yellow box shows the location of GFP-positive areas without cell bodies. Pink box shows the region with both GFP- and RFP- positive cells. Circle shows the position of the cell body and the triangle shows the targets of projections from the cell lineage. First few rows indicate the fish ID, serial number of clonal units in each fish, and which hemisphere each clonal unit was detected (L: left, R: right).

### Clonal units tend to project to the same target regions

We next asked whether individual members of a clonal unit exhibit similar projection patterns or whether they target a diversity of different regions. In *Drosophila*, clonal units project to the same target and work as connecting modules ^29^, whereas clonally-related cells in the mammalian visual cortex tend to connect to each other but do not project to the same targets ^30^. To assess this question in teleosts, we traced the axonal projections of the clonal units (Figure 2A, Figure S3). We find that first, axons and dendrites from the clonal units in the subpallium intermingled with each other, and they send projections randomly to both Dc and Dl regions. On the other hand, the pallial clonal units sent axon bundles, for the most part, to the same target region (Figure S3, *Methods*). Further we found many connections that project from and into the dorsal pallium (Dd2), which allow us to hypothesize a local pallial network around Dd2 as shown in Figure 2B.

**Figure 2.**
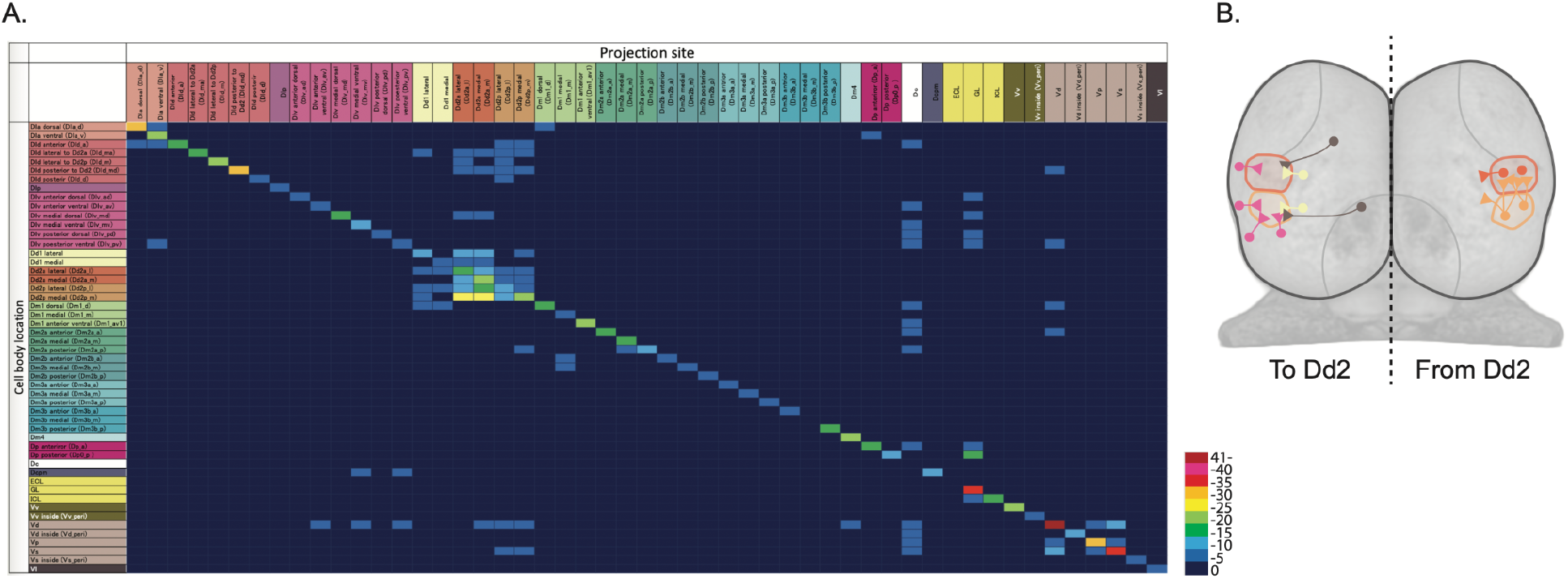
Neuron projections of clonal units in the telencephalon. (A) Connectional matrix between clonal units in the telencephalon. The rows indicate the cell body locations and the columns indicate where the projection ends. The color code indicate the number of the projection among 81 fish we analyzed. (B) Schematic drawing of the axonal projections from and into Dd2 regions across clonal units in the telencephalon.

### Distinct open chromatin structure in the medaka dorsal pallium (Dd2)

The results enumerated above prompted us to hypothesize that gene expression is uniquely regulated in each clonal unit (Figure 3). In order to test this theory we performed ATAC-seq on clonal units dissected from sliced brains to examine open chromatin regions (OCRs) that are either specific to, or shared across, the individually labeled clonal units (Figure 3AB, Figure S4). We generated ATAC-seq data from 100 sliced brain samples, and 65 samples met the criteria for screening high quality data (See Methods) (Figure 3C). To quantitatively test the relationship between the samples, we compared the ATAC-seq peak pattern (See Methods). Hierarchical clustering revealed that clonal units from the same brain regions tend to cluster together (Figure 3D). We then classified OCRs into 15 clusters (OCR Cs) by k-means clustering. We first found that OCR patterns between the subpallium and pallium were dissimilar, which is reflected by their distinct gene expression from the early developmental stage ^31^.

**Figure 3.**
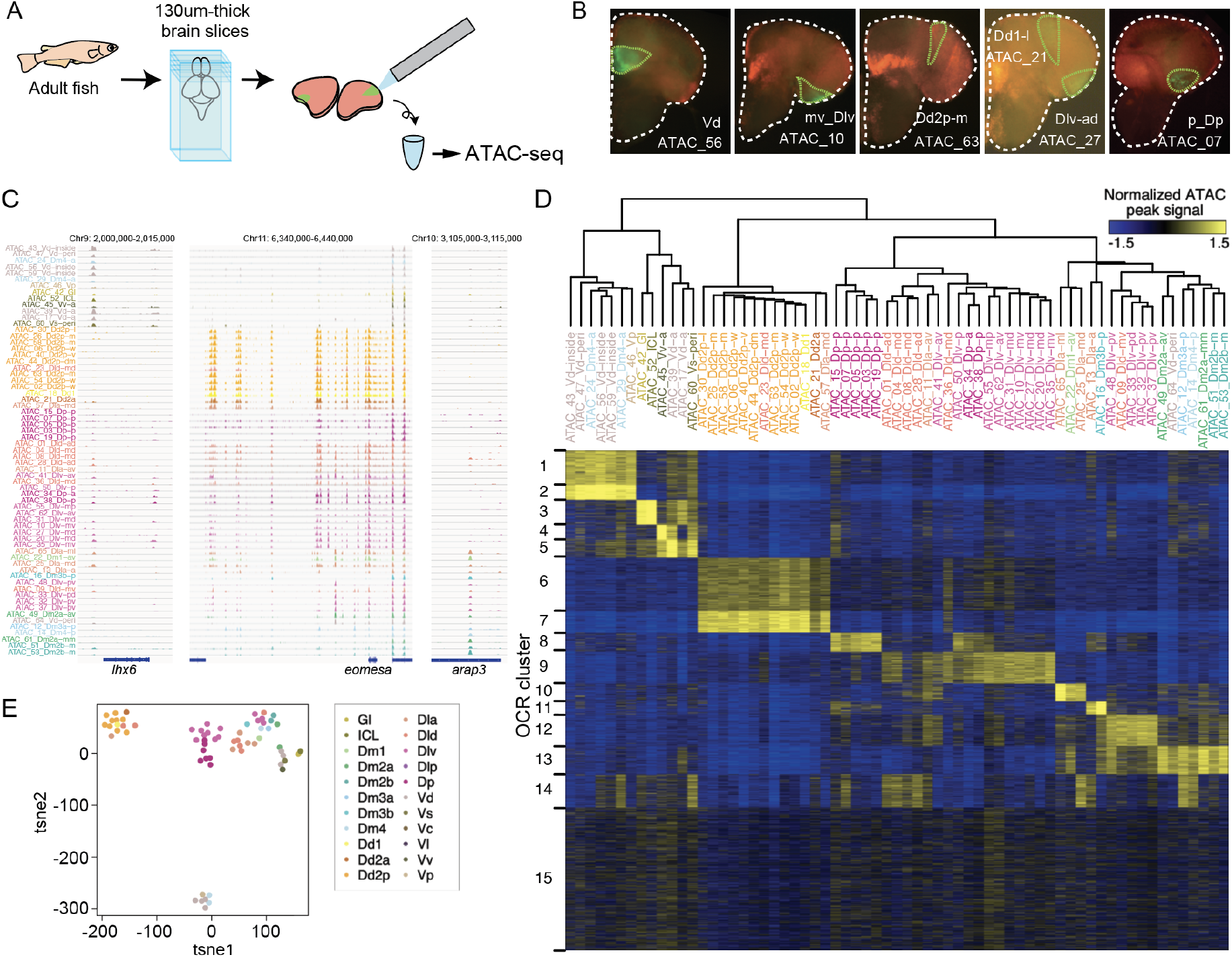
ATAC-seq of clonal units in the adult medaka telencephalon. (A) Procedure of ATAC-sequencing with extracted clonal units from the adult telencephalon. Cre-loxP recombination was induced at the neurula stage in the transgenic embryos (Tg (HSP-Cre) x Tg (HuC:loxP-DsRed-loxP-GFP)) as previously described. After making 130 um-thick brain slices at the adult stage, GFP-positive cell subpopulations were dissected and extracted manually. (B) Examples of GFP-positive clonal units in the brain slices. White dotted lines indicate the outline of the brain slices. Green lines indicate the dissected GFP-positive clonal units. (C) Representative track view of ATAC-seq. Colors indicate the location in the anatomical regions (Figure 1A) of the extracted clonal units. (D) Hierarchical clustering and k-means clustering of ATAC-seq peaks in clonal units. In the heatmap, blues indicate closed chromatin regions and yellow indicates open chromatin regions (OCR). Colors of clonal unit’s names indicate the anatomical regions. (E) Dimensionality reduction analysis (tSNE) of ATAC-seq peak patterns of clonal units.

Within the pallium, clonal units in the different sub-regions show unique patterns, where the medial and posterior pallium contained OCR C13 and OCR C8 respectively, and the ventral-lateral pallium (Dlv), the medial (Dlv-m) and posterior (Dlv-p) showed a variety of different OCRs (Figure 3D).

Remarkably, the clonal units in the dorsal pallium (Dd2) were found to have distinct chromatin structure compared to all other clonal units in the telencephalon (Figure 3D); OCR C6 and 7 were specifically open in the genome of Dd2 samples, and other OCR clusters which are open in other pallial regions (e.g. C8-14) tended to be closed in Dd2. Dimensionality reduction methods, such as principal component analysis (PCA), tSNE, and UMAP analysis, confirmed that, apart from subpallial clonal units, clonal units in Dd2 were distant from other pallial cell lineages (Figure 3E, Figure S4C). We examined the distribution of Dd2-specific ATAC-seq peaks in the genome, and they were located mainly in the intron and intergenic regions (Figure S4D), suggesting that the genomic regions that exhibit these peaks function as Dd2-specific enhancers. In summary, this ATAC-seq analysis suggests that the chromatin structures differ, depending on brain regions, and that it singles out the Dd2 region as an area of particularly distinct regulation.

### Enhanced transcriptional regulation on synapse-related genes in Dd2

Next, to uncover which genes are differentially regulated in Dd2, we examined Gene ontology (GO) term enrichment for genes targeted by each OCR cluster (Figure 4A, Figure S5). First we found that the terms related to axon guidance were enriched in various OCR clusters (Figure 4A top. OCR C1, 3-7, 9, 12, & 14), which suggests that different combinations of genes in the axon guidance pathways express in different clonal units.

**Figure 4.**
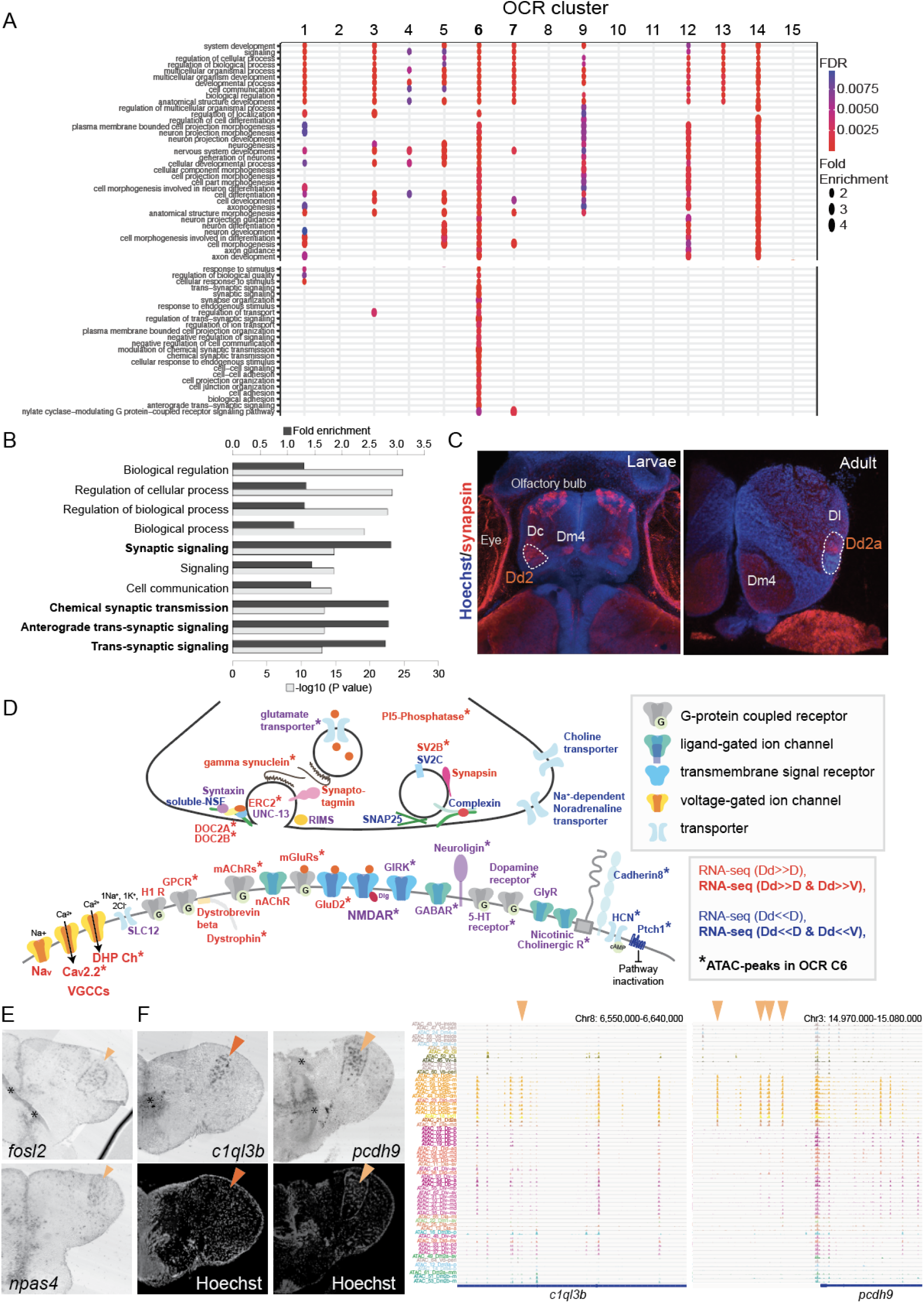
Analyses of Dd2-specific OCR clusters reveal specialized synaptic architecture. (A) Gene ontology (GO) analysis on OCR clusters. GO terms related to axon guidance (top) and synaptic genes (bottom) were shown (See also Figure S6 for other enriched GO terms). Size and color of circles indicate fold enrichment and FDR, respectively. Circle was plotted only if FDR is lower than 0.01. Axon guidance pathways are enriched with pallial clusters (OCR C9 & 12), Dd2 cluster (OCR C6), subpallial cluster (OCR C1, 3, 4, 5) and common cluster (OCR C14). On the other hand, synaptic genes are enriched in OCR C6. (B) GO term enrichment analysis on the genes highly expressed in Dd2. Top10 significantly enriched terms are shown. (C) Anti-synapsin immunohistochemistry on the medaka larvae and adult telencephalon. Blue: Hoechst, Red: anti-synapsin signals. In larvae, strong signals were detected in the olfactory bulbs, Dc, Dm4 and Dd2. In the adult telencephalon, the signals were broadly detected and strongly detected in Dd2a. (D) Schematic summary of the genes actively regulated or preferentially expressing genes in Dd2 regions. Gene names in red indicate all subunits in the gene expressed significantly higher in Dd2 than in other pallial regions, while gene names in blue indicate the expression of all subunits in Dd2 was significantly lower in Dd2. Gene names in purple indicate that the gene included both higher and lower expressing subunits. * indicates the genes that are actively regulated by the Dd2-specific OCR cluster (C6). More detailed information about the differentially regulated subunits of genes is in Figure S7. GABAR: GABAergic receptor, GIRK: Glutamate receptor ionotropic kainate, GluD2: Delta glutamate receptor2, GlyR: Glycine receptor, H1R: histamine receptor, mAChRs: muscarinic Acetylcholine receptors, nAChR: nicotine Acetylcholine receptor, SLC12: kidney-specific sodium-potassium-chloride cotransporter, CaV2.2: N-type voltage-gated calcium channels, DHP Ch: L-type voltage-gated calcium channels, VGCCs: voltage-gated Calcium channels, Nav: voltage-gated sodium ion channels, PI5-phosphatase: Phosphoinositide5-Phosphatase. (E) Expression of immediate early genes (*fosl2* and *npas4*) in the Dd2p region (orange triangles). (F) Expression of synaptic genes specifically expressed in Dd2 visualized by *in situ* hybridization (left). *c1ql3b* was specifically expressed in Dd2a (top left), and *protocadherin9* was detected in Dd2p (bottom left). Orange triangle indicates the gene expression in Dd2. Track view of ATAC-seq peaks in clonal units (right). Orange triangles indicate the Dd2-specific ATAC-seq peaks.

Intriguingly, one of the Dd2-specific OCR clusters, OCR C6, was enriched with GO terms related to synapse-related genes (Figure 4A bottom). As open chromatin regions reflect the transcriptional regulation in both enhancing and suppressing directions, we analyzed the actual expression of synapse-related genes in Dd2 by RNA-seq on Dd2, the pallium regions excluding Dd2 (D), and the subpallium (V) (Figure S6 AB). First, GO term enrichment analysis shows that synaptic signaling-related terms are significantly enriched in genes preferentially expressed in Dd2 (i.e. expressed significantly higher in Dd2 than D) (Figure 4B). Among 82 synaptic genes preferentially expressed in Dd2, 68 % of them were targeted by OCR C6 or C7, which we call “actively up-regulated synaptic genes”. On the other hand, 70 synaptic genes were significantly lowly expressed in Dd2 compared to D, and 40% of them were the targets of OCR C6 which we call “actively down-regulated synaptic genes” (Figure S6 C).

Since the transcription of synaptic genes are actively regulated in Dd2, we checked if synaptic density is higher in the dorsal pallium (Figure 4C, Figure S6D). We observed the high expression of synaptic markers in both larval and adult telencephalon in medaka; postsynaptic marker PSD95 in both Dd2a and Dd2p, and presynaptic marker synapsin in Dd2a. Also, to see if this synapse-enriched area in the dorsal pallium is unique in medaka, we also examined the expression of synaptic marker genes in zebrafish larvae, and PSD95- or synapsin-signals were detected in Dc but not in the dorsal pallium in zebrafish (Figure S6D).

In the end, to characterize the synaptic architecture in the dorsal pallium, we assessed the “actively-regulated synaptic genes” in Dd2 (Figure 4D, Figure S7). Subunits of the glutamate receptors, transporters and modulatory neurotransmitter receptors (5-HT receptors, cholinergic receptors and dopamine receptors) were differentially regulated in both positive and negative directions. On the other hand, a variety of voltage-dependent calcium channels expressed significantly higher in Dd2 than other pallial regions, whereas the inhibitory synaptic genes were significantly lowly expressed. These results implied that neurons in Dd2 maintain persistent activity, which is consistent with the expression of a couple of immediate early genes in Dd2 (Figure 4E). Lastly, among the up-regulated synaptic genes in Dd2, we found a few genes specifically expressed in Dd2 by *in situ* hybridization (Figure 4F); synaptic regulator, *c1ql3b*, in Dd2a; a protocadherin gene, *pcdh9*, in Dd2p.

### Transcriptional regulators of Dd2-specific OCR clusters

Finally, we focused our attention on the mechanism that regulates Dd2-specific gene transcriptions (Figure 5). First, we used ATAC-seq data of clonal units to identify candidates of regulators by searching the known TF binding motifs enriched in the OCR clusters (Figure 5A). In the Dd2-specific OCR clusters (C6 & 7), we found that binding motifs of beta helix-loop-helix (bHLH), zinc finger (Zf) and T-box families’ were significantly enriched, suggesting that multiple TFs regulate the Dd2 gene expressions. We also performed *de novo* motif searching in Dd2-specific OCR clusters and found that the result was consistent with the known-motif search (Figure 5B).

**Figure 5.**
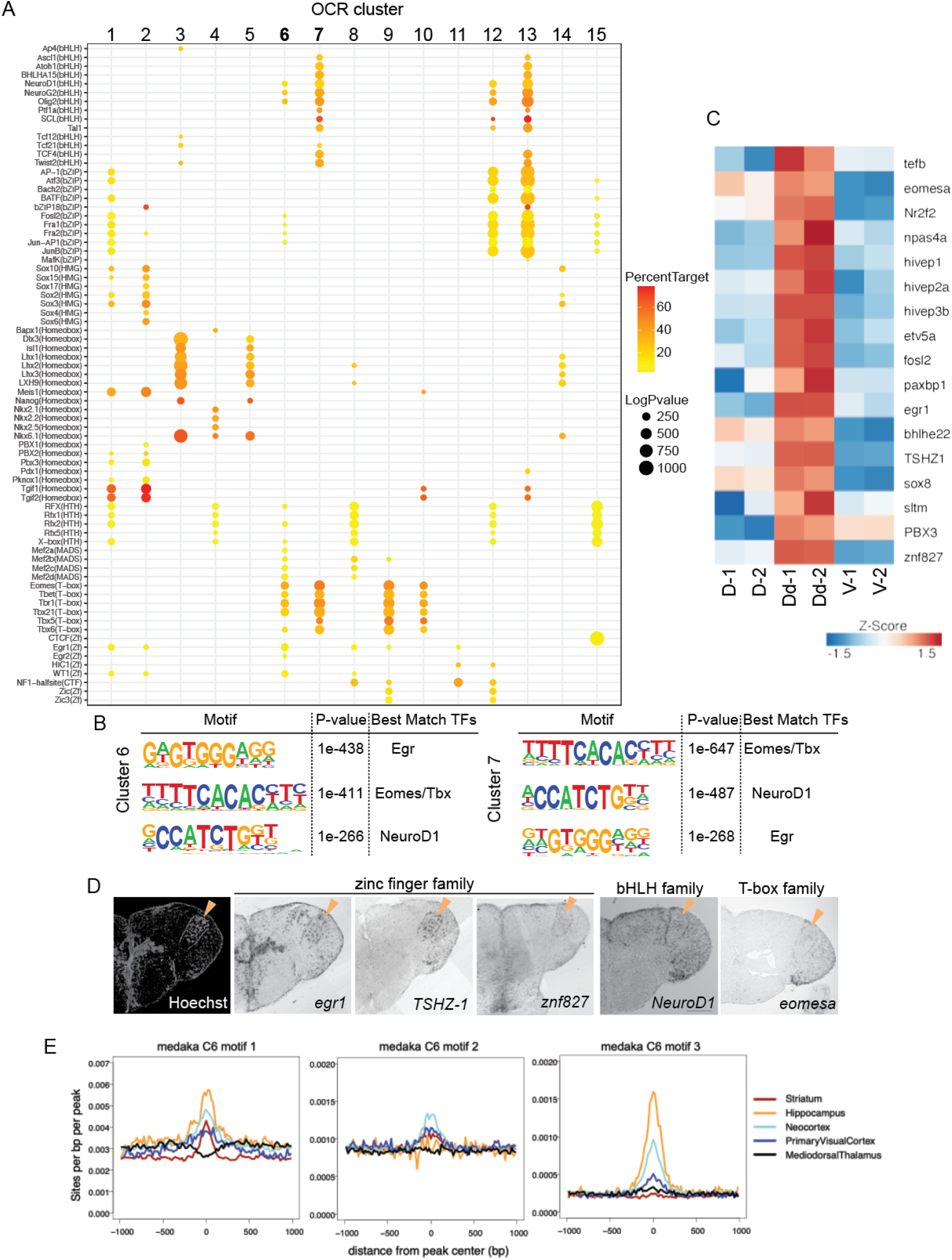
Transcriptional regulation in Dd2. (A) Enrichment of known transcription factor binding motifs in OCR clusters are shown as dot plots. Size and color of circles indicate −log_10_(*p*-value) and percent target, respectively. Circle was plotted only if −log_10_(*p*-value) is higher than 100. OCR C 6 & 7 which are specifically open in Dd2 are enriched with transcription factors of bHLH, bZIP, T-box and Zf family. (B) Result of *de novo* motif finding analysis. Top 3 of the enriched motifs in OCR C6 & 7 and candidate transcription factors that have similar binding motifs are shown. (C) Expression of candidate transcription factors in Dd, the pallium except Dd (D), and the subpallium (V) analyzed by RNA-sequencing (n=2 replicate). (D) Expression pattern of the candidate transcription factors in the telencephalon visualized by *in situ* hybridization. Orange triangles indicate the position of Dd2p. (E) Medaka OCR C6 motif enrichment in human brain region-specific OCRs. Existence of each medaka C6 motif was examined around human brain ATAC-seq peaks, and the motif density was calculated.HC: hippocampus, MDT: medial dorsal thalamus, NCX: neocortex, PVC: primary visual cortex, ST: striatum. Darker blue shades indicate higher correlation.

Next, we assessed the actual expression of the candidate regulators of Dd2. In the RNA-seq data, we found TFs whose binding motifs that were enriched in Dd2-specific OCR clusters were preferentially detected in the Dd2 sample (Figure 5C). Then, we examined the expression of candidate TFs by *in situ* hybridization (Figure 5D, Figure S8). Though we didn’t find a transcription factor that expressed only in Dd2 regions, we found several candidate TFs expressed strongly in Dd2; *tshz-1* (Zf family) expressed strongly in Dd2 and weakly in Dl; several other Zf family, such as *egr-1, npas4*, and *znf827*, expressed in Dd2 and other part of the telencephalon; *eomesa* (T-box family) expressed in Dd, Dl and Dp. *TSHZ* is a member of the C2H2-type zinc-finger protein family. Though its binding motif is unknown, *TSHZ* might possess a binding motif similar to that of other C2H2 zinc-finger proteins. Intriguingly, the motif of *Egr*, a C2H2 zinc-finger protein, was found to be top enriched ones in Dd2-specific OCR clusters (C6 & 7; Figure 5B), which implies that *TSHZ-1* plays a critical role in the differentiation of the Dd2 region.

In the end, we questioned whether medaka Dd2 is homologous to any region in the mammalian pallium (Figure 5E). Since teleost fish and humans are phylogenetically very distant, it is difficult to find corresponding regulatory sequences between the two genomes. Thus, we compared putative transcriptional regulators between medaka clonal units and human brain regions from Fullard et al. ^32^, by investigating the TF motif enrichment in the open chromatin regions.

First, we compared the significantly enriched TF motifs between medaka OCR clusters and human differential OCRs ^32^, by counting the number of overlaps. We found that our medaka OCR clusters shared only few motifs with the human medial dorsal thalamus (MDT), which is consistent with the fact that the thalamus does not belong to the telencephalon (Figure S8B). Also, we confirmed that the subpallium specific medaka OCR cluster (C1) shared a relatively larger number of motifs with the human striatum (ST), a part of subpallium in mammals (Figure S8B). The medaka Dd2-specific OCR clusters (C6 and C7) share motifs most with the human hippocampus (HC), neocortex (NCX), and ST (Figure S8B). However, this analysis might suffer from bias when there are many similar motifs from the same TF family. Thus, we next examined the enrichment of Dd2-specific OCR motifs in human brain open chromatin regions. As shown in Figure 5B, medaka OCR C6 and C7 were both enriched with three TF motifs that best match to Egr, Eomes/Tbx, and NeuroD1 motifs. We therefore examined the distribution of the motif 1, 2, and 3 of OCR C6 in human ATAC-seq peaks. We found that motif 1 and 3 were enriched in HC and NCX open chromatin regions, and motif 2 was enriched in NCX the most (Figure 5E). These results suggest that medaka Dd2 partially share the similar transcriptional regulators with the human hippocampus & neocortex.

## Discussion

Here we report that the dorsal pallium (Dd2) in medaka is formed by clonal units and is distinct from other pallial regions in terms of epigenetic regulation, especially of synaptic architecture (Figure 6).

**Figure 6.**
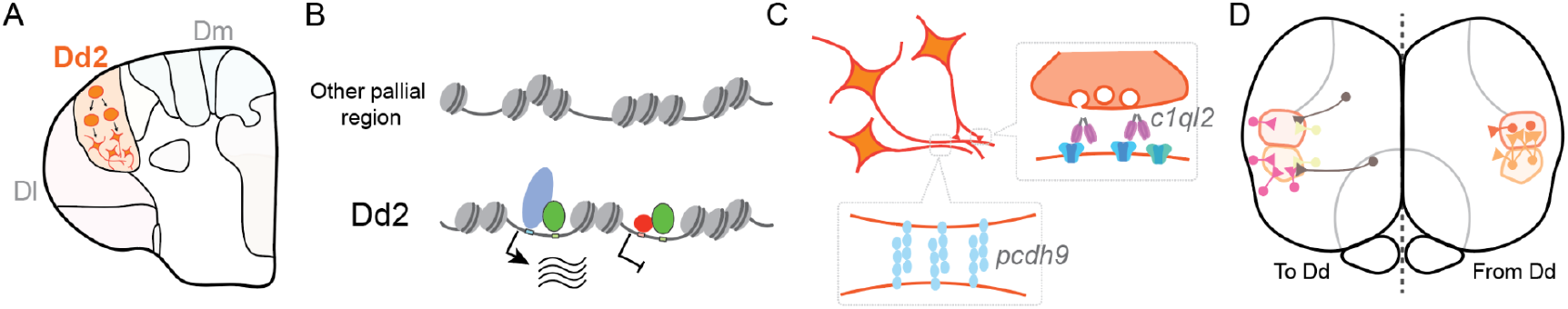
Hypothesized molecular model of specialized synaptic architecture in the dorsal pallium in medaka. Dd2 region is formed by a few clonal units (A). In Dd2, gene transcription was distinctly regulated by the combination of multiple TFs which construct a distinct open chromatin structure (B). This specific OCR contributes to the regulation of specialized synaptic-related gene expression in Dd2, which are suggested to generate specific axonal projections and synaptic architecture (C). We assume that Dd2 is important in the neural networks in the telencephalon of medaka (D). Circles indicate the cell bodies and triangles indicate the projectional terminals.

### Pallial diversity in teleost fish

The pallium in teleosts has been mysterious, since different species exhibit, within its boundaries, different numbers or cellular densities of compartmentalized anatomical sub-regions. On the other hand, the subpallial regions are relatively conserved among teleost species. Our structural analysis in the adult medaka telencephalon revealed that the clonal architecture between the pallium and subpallium differs in the distribution of cells in clonal cells, which possibly explains how the difference in diversity between the pallium and subpallium has emerged: the subpallium is conserved because cells belong to various clonal units intertwined with each other, which has constrained their modification during evolution; whereas the pallium is diverse because of the modular nature of the clonal units which allows for the emergence of diversity.

How did the dorsal pallium evolve in teleosts? One hypothesis is that the teleostean dorsal pallium has been specialized from the teleostean lateral pallium (Dl) ^2^, whereas an alternative hypothesis proposes that the teleostean dorsal pallium evolved independently ^33^. Our results favor the first, based on the molecular mechanisms we uncovered through analysis of expression patterns of transcription factors (TFs) in medaka and zebrafish. Our RNA-seq data identified a couple of TFs strongly expressed in Dd2 in medaka fish. In zebrafish, similar TFs, such as *TSZ-1,eomesa, egr-1*,are expressed in Dl ^34^, suggesting that medaka Dd2 shares a major part of regulatory mechanism with Dl, so Dd2 perhaps is derived from Dl.

Since medaka belong to the Beloniformes family which diverged later than the Cypriniformes family which includes zebrafish, it might be possible that the last common ancestors of bony fish possess the dorsal pallium when they diverged, and the dorsal pallium disappeared later in Cypriniformes family or it is not yet identified in Cypriniformes family. Further analysis in other fish species will be required to explain the evolutionary trajectory of the dorsal pallium.

### Possible neural computations and functions of medaka Dd2

Numerous computations happen in synapses, and synaptic architecture is defined by the expression of a subset of specific molecules. Some neurons in mammals and songbirds exhibit synapses which are characterized by particularly strong plasticity and modulation, where complicated information processing takes place or memories are stored ^35 36^. In our paper, we found that many synaptic genes are differentially regulated in Dd2 and they span the range from excitatory, inhibitory and modulatory synaptic genes. We also identified a synaptic organizer, *C1ql3*,which expressed specifically in the anterior Dd2 (Dd2a), and also expresses in the mammalian hippocampus where it is known to organize extracellular postsynaptic space and bind to GPCRs in a calcium dependent way ^37,38^. Taken together, our results suggest that the transcription of synaptic genes in the dorsal pallium is selectively regulated and might assist information processing and activity dependent plasticity in a manner different from other pallial regions.

What role does the dorsal pallium (Dd2) in medaka play within the telencephalic network? Our findings suggest that Dd2 might function as either a network hub where the flow of information is gated through synaptic modulation ^39^, or, alternatively, it might serve as a working memory module that can transiently store information by persistent activity. Several lines of evidence support this notion. First, a couple of immediate early genes are expressed in Dd2. Second, from structural analysis, we observed axonal projections from many surrounding regions into Dd2 as well as dense and, usually unilateral, connectivity between Dd2 subregions (from Dd2p to Dd2a). Third, cadherin molecules are known to play an important role in regionally targeted axonal projection ^40^, and we found specific expression of *pcdh9* in Dd2p ^41^. Here, it should be noted that our axonal projection analysis might have missed the projections from mature neurons, because the HuC promoter labels preferentially neural progenitors and disappears at later stages ^19^. Thus, brain-wide exhaustive connectome analysis by electron microscopy ^42^ or neurophysiological analysis will be necessary for further study ^43^.

At present, few studies have been done to uncover the behavioral function of the dorsal pallium in teleosts: in electric fish, projection analysis suggests that the dorsal pallium possess hippocampal-like circuitry ^44^. In several cichlid fish species, the dorsal pallium gets activated specifically in the context of social memory tasks ^45^. Further, some teleostean species that show social behaviors requiring memory, exhibit distinct cytoarchitecture in their dorsal pallium. These include cichlid fish that show social dominance ^46^ and, interestingly, medaka fish show mating behavior involving social memory ^47,48,49^. It should be the work of the future to study whether Dd2 in medaka functions in the context of social behavior using memory.

### Mammalian brain regions homologous to the teleost dorsal pallium

The homology of the anatomical region in the telencephalon across vertebrates has been a subject of intense debate. Especially the correspondence of the pallial regions between teleosts and amniotes has been controversial, since the growth of neural tubes in early development proceeds in the opposite direction, such as evagination in amniotes and eversion in teleosts ^50,51^. Traditionally, connectional and immunohistochemical analyses were performed to study this homology ^1^.

In the current study, ATAC-seq analysis provides an orthogonal approach that can directly perform cross-species comparison of the regulatory elements in a genome-wide fashion. Also, we applied ATAC-seq and RNA-seq to clonal units from the brain slices rather than single cells so that we could trace the spatial information of the chromatin and transcripts in the samples. Our analysis provides insight that medaka Dd2-specific open chromatin regions shared the binding motifs of transcriptional regulators with the human neocortex and hippocampus. Though not so many studies are done in the human brain, some studies in mammals suggest that synaptic architecture is regulated heavily in the hippocampus and neocortex, where density and size ^52^ of synapses is controlled, presumably in the context of synaptic plasticity ^53^.

Altogether, we propose that, at least in medaka, the dorsal part of the pallium accommodates a region whose synaptic architecture is distinctly regulated by unique chromatin structure, and might partially share the same transcriptional regulators with hippocampus and neocortex in mammals. Further studies of the dorsal pallium with an epigenetic approach will shed light on the evolutionary mechanisms of brain diversity.

## Acknowledgements

We are grateful to many thoughtful comments and heartfelt support from Takeo Kubo in the University of Tokyo. We extremely thank Florian Engert in Harvard University for reading and editing the manuscript. We acknowledge comments from Kei Ito in Koln University. We appreciate the financial and experimental support from Florian Engert lab, Nicholas Bellono lab, Sophia Liang in Catherine Dulac lab, and Burcu Erdogan in Jessica Whited lab in Harvard University. This work was supported by NIBB Collaborative Research Program (16-522, 17-513 and 18-513) to HTakeuchi; Japan Society for the Promotion of Science (JSPS) KAKENHI Grant Numbers JP 16H06987 (to YI), 16KT0072 (to HTakeuchi), 20K20303 (to YK); Japan Society for the Promotion of Science (JSPS) grant number JP21K06013 and Grant-in-Aid for Scientific Research on Innovative Areas grant number JP21H00245 to R.N.

## Author contributions

Conceptualization, YI, and RN; Methodology, YI, RN, YK, SN; Data analysis, RN, YI, NY; Writing - Original Draft, YI; Writing - Review & Editing, RN, HTakeda, NY, TO, HTakeuchi, YK, SN; Visualization, YI, RN; Funding Acquisition, YI, RN, HTakeuchi.

## Methods

### Ethics statement

All experiments were conducted using protocols specifically approved by the Animal Care and Use Committee of the University of Tokyo (permit number: 12–07). All surgeries were performed under anesthesia using MS-222, and all efforts were made to minimize suffering, following the NIH Guide for the Care and Use of Laboratory Animals.

### Fish husbandry

Medaka fish are kept at 14h/10h on light at 28 °C. The transgenic line (Tg) (HuC:loxp-DsRed-loxp-GFP) and Tg (HSP:Cre/Crystallin-CFP) were generated as previously reported ^19^. Both males and females were used in the experiments.

### Clonal units visualization

To induce Cre/loxP recombination, embryos of the double Tg line, being crossed from female Tg (HuC:loxp-DsRed-loxp-GFP) and male Tg (HSP:Cre) line, were mildly heated at 38 °C for 15 min. Heat shock was applied using a thermal cycler to polymerase chain reaction tubes containing two eggs each. After the mild heat shock treatment, the embryos were maintained at 26°C until they hatched. Then the fish were kept in the fish tank and raised till the adult stage.

### Whole-brain clearing and imaging

Adult fish brains were dissected by the usual procedure ^54^. After the dissection, the brains were fixed in 4% paraformaldehyde/phosphate-buffered saline (PBS) overnight. Then the brains were washed twice with PBST (0.5% Triton-X100 in PBS) and immersed into Sca/eA2 solution for about three hours on ice ^26^. When the brains were confirmed to be transparent, they were transferred into a new Sca/eA2 solution with DAPI (0.5 ul/2ml) overnight at 4°C. The next day, the brains were transferred into the new Sca/eA2 solution and washed with PBST. To prepare samples for signal detection, the brains were embedded in 0.75% agarose gel with Sca/A2-Triton solution and fixed on glass coverslips. Fluorescent signal detection was performed using a hand-made light sheet microscopy Digital Light-sheet Microscope (DSLM)^55^ and a commercial one (Zeiss, Lightsheet Z.1).

In order to define the anatomical regions, we used DAPI signals as landmarks. For whole-brain imaging, we dissected adult medaka brains, fixed with 4% paraformaldehyde/PBS, stained with DAPI in Sca/eA4 (0.5/2000) for two days and soaked with Sca/eA4 for several days.

### Brain registration

To compare fluorescent signals among multiple brain samples, we performed image registration with CMTK registration GUI in Fiji ^28^ for details, please see: https://github.com/jefferislab/BridgingRegistrations) as described in a previous report ^29^. To make a reference brain, we used the DAPI signals of the best images obtained by light-sheet microscopy and processed with Gaussian blur. Three-dimensional reconstruction images were made with FluoRender ^56^.

### Clonal units identification

To identify the structure of clonal units, first we compared the structure of GFP-positive neurons among the registered brains in FluoRender ^56^. We extracted GFP-positive clusters as clonal units when we found GFP-positive regions overlapped in multiple registered brains. We tracked the neuron projection by observing both registered brains and raw images.

### Immunohistochemistry & In situ hybridization

In order to define the anatomical regions, we examined the expression of marker genes by performing immunostaining as previously reported ^19^. Briefly, after cutting 14-um thick cryosections of adult brains, sections were incubated overnight with the primary antibody (anti-CaMK2α (abcam, ab22609), parvalbumin (Millipore, MAB1572), GAD65/67 (Sigma, G5163) and GAD 67 (Sigma G5038), 1:2000 each), synapsin (Synaptic Systems, 106 011), PSD (abcam, ab18258) in 1% BSA/DMSO/Triton-X1000/PBS. Though we also tried anti-GAD 65 so far, we couldn’t find the difference of intensity inside among anatomical regions. After washing several times with DMSO/Triton-X1000/PBS, the sections were incubated with fluorescent Alexa 488 (ThermoFisher, 1:1000) and DAPI (1:4000) in 1% BSA/DMSO/Triton-X1000/PBS.

To analyze the expression of genes in the pallium, we performed *in situ* hybridization as previously reported ^57^. Primers designed for making RNA probes were listed below.

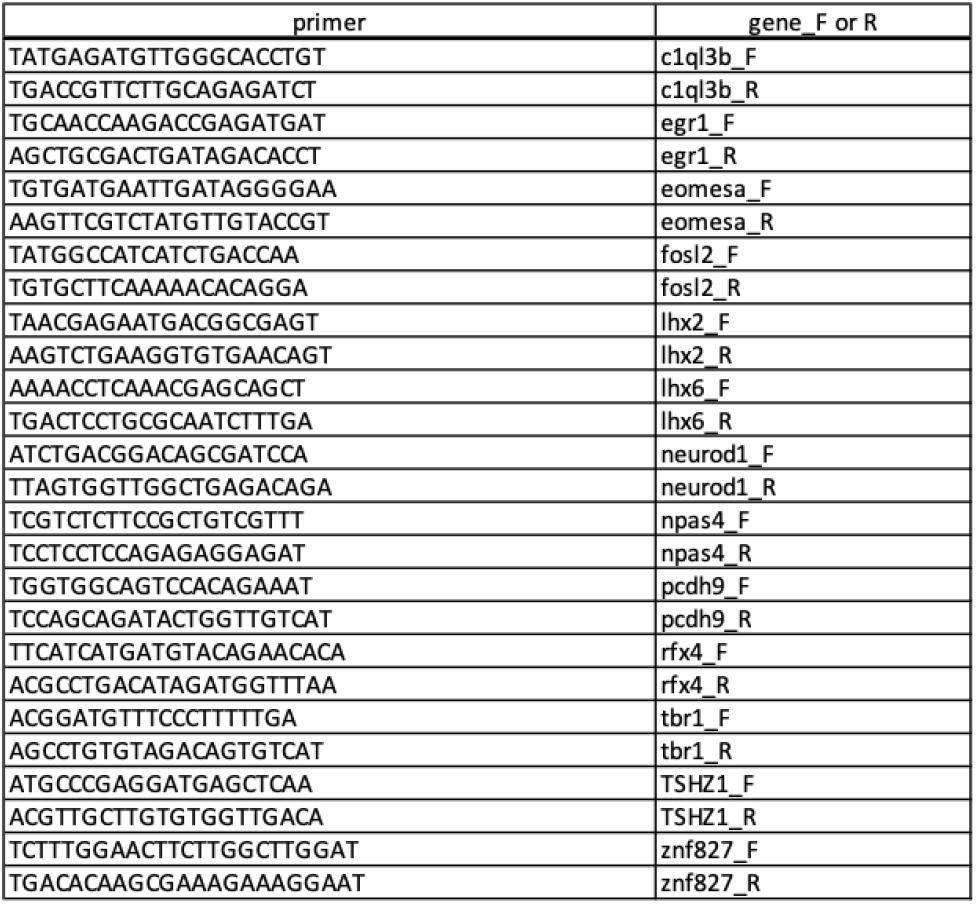

### Brain slicing for ATAC-seq and RNA-seq

Adult medaka fish of Tg (HuC:DsRed) were anesthetized using tricaine methanesulfonate (MS222) solution (0.4%). After brain dissection, brains were immersed into artificial cerebrospinal fluid (ACSF) on ice. Then, brains were embedded in 2.5% low-melting agarose in ACSF and frozen at −20°C for 8 min. After removing the excess agarose gel, 130-μm thick brain slices were cut using a vibrating blade microtome (Leica VT1000S). The slices were collected, placed onto the slide glass, and dissected using a razor blade under a fluorescence microscope based on the DsRed or GFP signal (Leica MZ16F).

### ATAC-seq

ATAC-seq was performed as previously described ^58^ with some modifications. After collecting the GFP-positive compartments into a 1.5-ml tube with PBS, the nuclei were extracted in 500 μl of cold lysis buffer(10 mM Tris-HCl pH 7.4, 10 mM NaCl, 3 mM MgCl_2_, 0.1% Igepal CA-630), centrifuged for 10 minutes at 500 x g, and supernatant was removed. Tagmentation reaction was performed as described previously ^58^ with Nextera Sample Preparation Kit (Illumina). After tagmented DNA was purified using MinElute kit (Qiagen), two sequential PCR were performed to enrich small DNA fragments. First, 9-cycle PCR were performed using indexed primers from Nextera Index Kit (Illumina) and KAPA HiFi HotStart ReadyMix (KAPA Biosystems), and amplified DNA was size selected to a size of less than 500 bp using AMPure XP beads (Bechman Coulter). Then a second 7-cycle PCR was performed using the same primer as the first PCR, and purified by AMPure XP beads. Libraries were sequenced using the Illumina HiSeq 1500 platform.

### RNA-seq

After slicing the Tg (HuC:DsRed) brain using a vibratome, the pallium, sub-pallium and Dd region were dissected based on the DsRed signals and collected in Trizol (Thermo Fisher Scientific). 30 fish were used for one replicate and two replicates were made. Brain slices were homogenized in 1 ml of Trizol, and 200 μl of chloroform was added, and total RNA was isolated using RNeasy MinElute Cleanup Kit (Qiagen). mRNA was enriched by poly-A capturing and RNA-seq libraries were generated using KAPA mRNA HyperPrep Kit (KAPA Biosystems).Libraries were sequenced using the Illumina HiSeq 1500 platform.

### ATAC-seq data processing

The sequenced reads were preprocessed to remove low-quality bases and adapter derived sequences using Trimmomatic v0.32 ^59^, and then aligned to the medaka reference genome version 2.2.4 (ASM223467v1) by BWA ^60^. Reads with mapping quality (MAPQ) larger than or equal to 20 were used for the further analyses. MACS2 (version 2.1.1.20160309) ^61^ was used to call peaks and generate signals per million reads tracks using following options; ATAC-seq: macs2 callpeak --nomodel --extsize 200 --shift -100 -g 600000000 -q 0.05 -B --SPMR.

We evaluated the quality of our sequenced samples by the following methods and used only high quality ATAC-seq data. First, we counted the number of mapped reads after removing redundant reads, and selected samples with 2 million reads or more. Then, we calculated the fraction of reads in peaks (FRiP) values ^62^, and selected the data that have a FRiP value of 0.2 or higher.

We collected all peaks from 65 ATAC-seq data that met the above criteria and merged the peaks if there were overlap between them. For each peak region, the number of mapped reads per total reads were calculated for each sample, and then normalized by the average of all samples.

Hierarchical clustering was performed by calculating a Euclidean distance matrix and applying Ward clustering ^63^ using the *hclust* and *dendrogram* functions in R. ATAC-seq peaks were clustered using k-means clustering using *kmeans* function in R. Principal component analysis (PCA)was performed using the *prcomp* function in R. t-distributed stochastic neighbor embedding (tSNE)^64^ plot was generated using *Rtsne* package in R, with perplexity=5 option. Uniform manifold approximation and projection (UMAP)^65^ was performed using *umap* package in R, with n_neighbors=7 and n_components=2 options.

To identify the target gene of each ATAC-seq peak, the closest transcription start site (TSS) was searched, and determined as the target gene if the distance to the TSS was less than 10 kb.

### RNA-seq data processing

The sequenced reads were preprocessed to remove low-quality bases and adapter derived sequences using Trimmomatic v0.32 ^59^, and were aligned to the medaka reference genome version 2.2.4 (ASM223467v1) by STAR ^66^ and reads with mapping quality (MAPQ) larger than or equal 20 were used for the further analyses.

Genes expressing significantly higher or lower in Dd2 were identified using DESeq2 (padj <0.05) ^67^.

### Gene Ontology (GO) analysis

Gene Ontology (GO) analyses were performed using the Gene Ontology resource ^68^.

### TF Motif identification

MotifsGenome.pl script from HOMER program ^69^ was used to identify the enriched motifs at ATAC-seq peaks.

### Comparison between medaka and human ATAC-seq data

We used region-specific (neocortex, primary visual cortex, hippocampus, mediodorsal thalamus, and striatum) OCRs from human brain ATAC-seq data ^32^. Enrichments of known motifs were compared between medaka OCR clusters and human region-specific OCRs, and the number of shared motifs was counted. The annotatePeaks.pl script from HOMER program ^69^ was used to examine the enrichment of medaka OCR C6 motif 1, 2, and 3 in human region-specific OCRs with -size 2000 -hist 20 options.

### Data access

Sequencing data generated in this study have been submitted to the DDBJ BioProject database under accession number PRJDB14398.

## Supplementary figures

**Figure S1.**
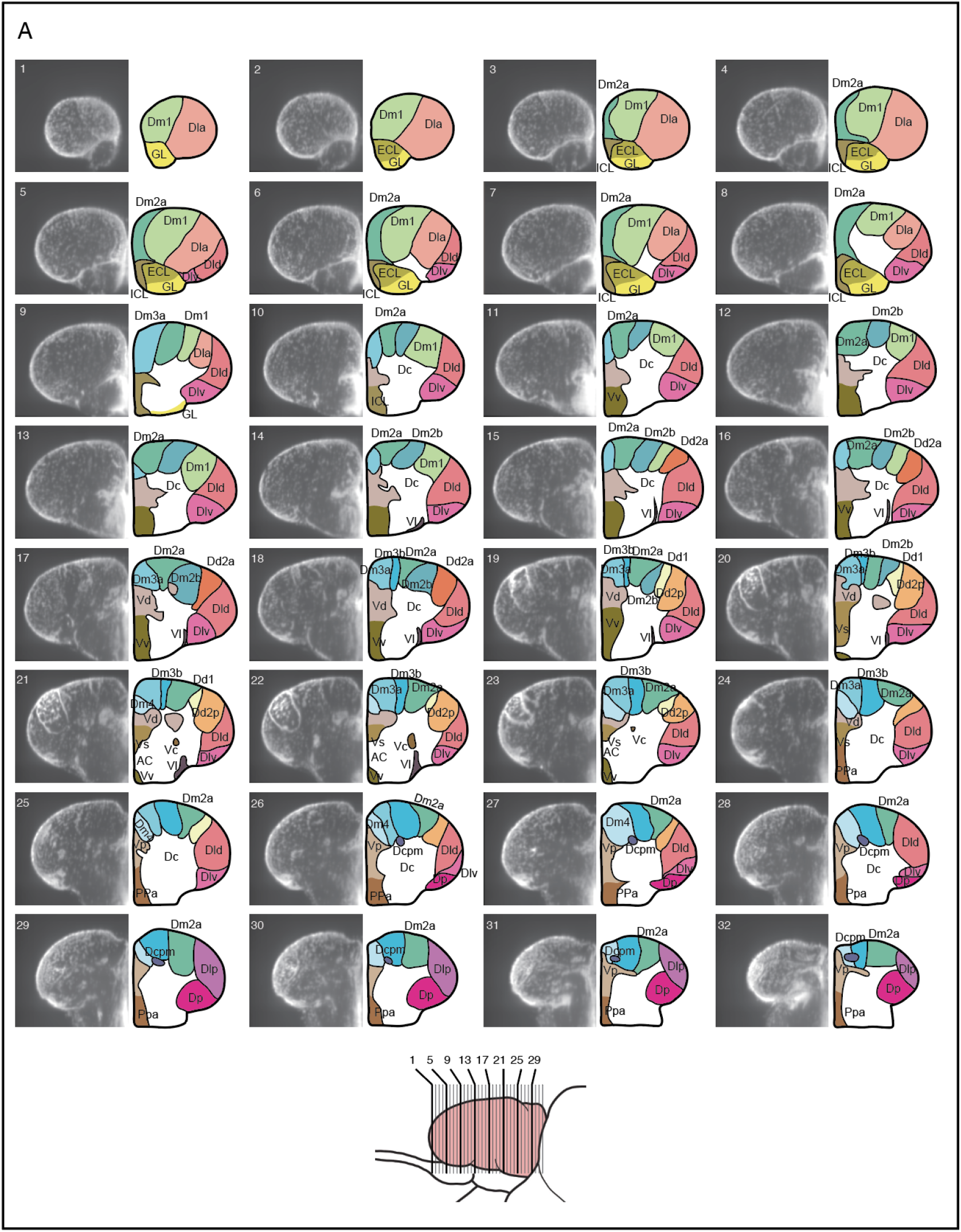
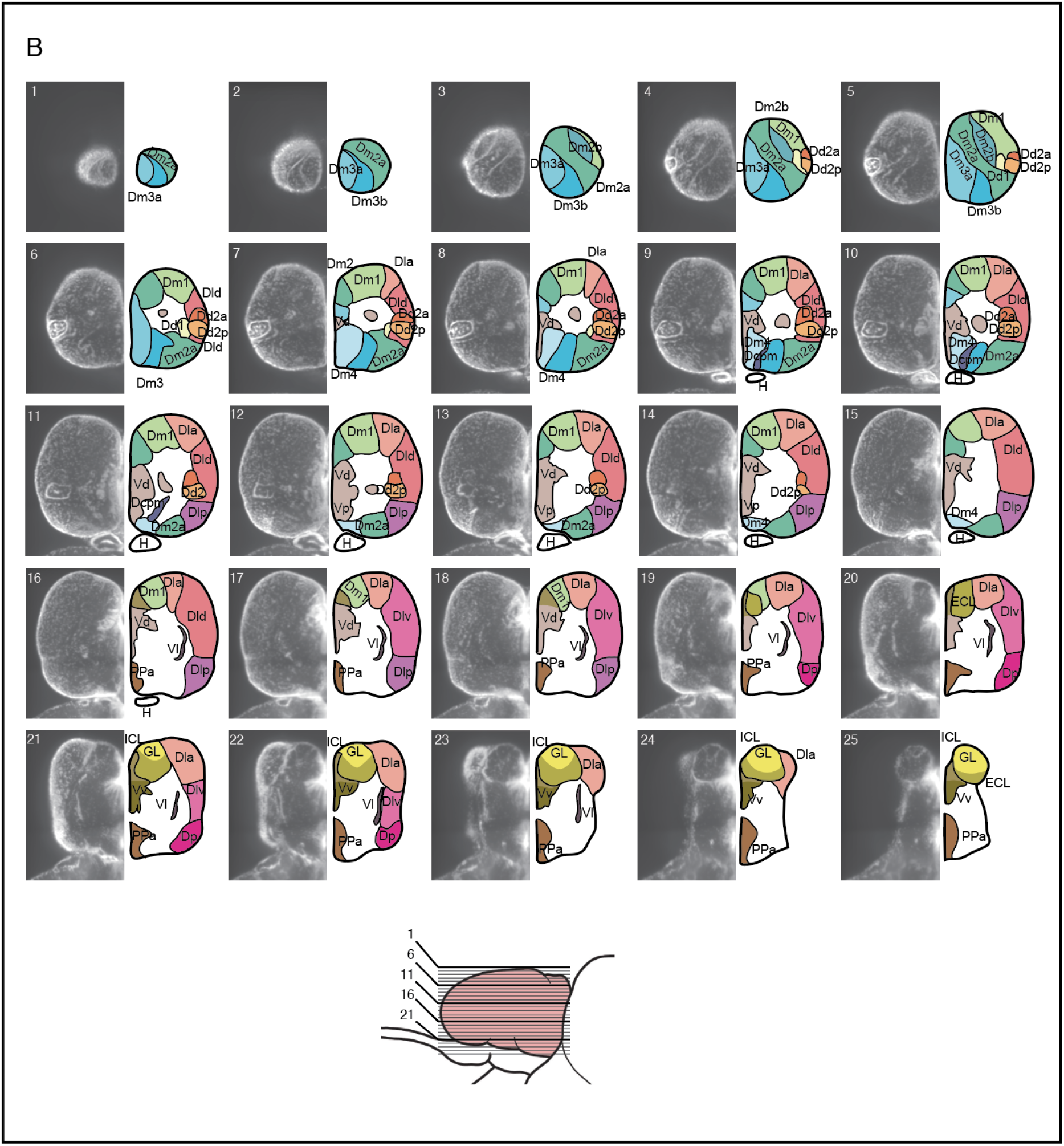
Details of re-defined atlas of the telencephalon. Transverse (A) and horizontal (B) optical sections of the reference telencephalon (left) and re-defined brain atlas (right). Lines in the schematic drawing of the lateral view of the telencephalon indicate the locations of the optical sections (bottom).

**Figure S2.**
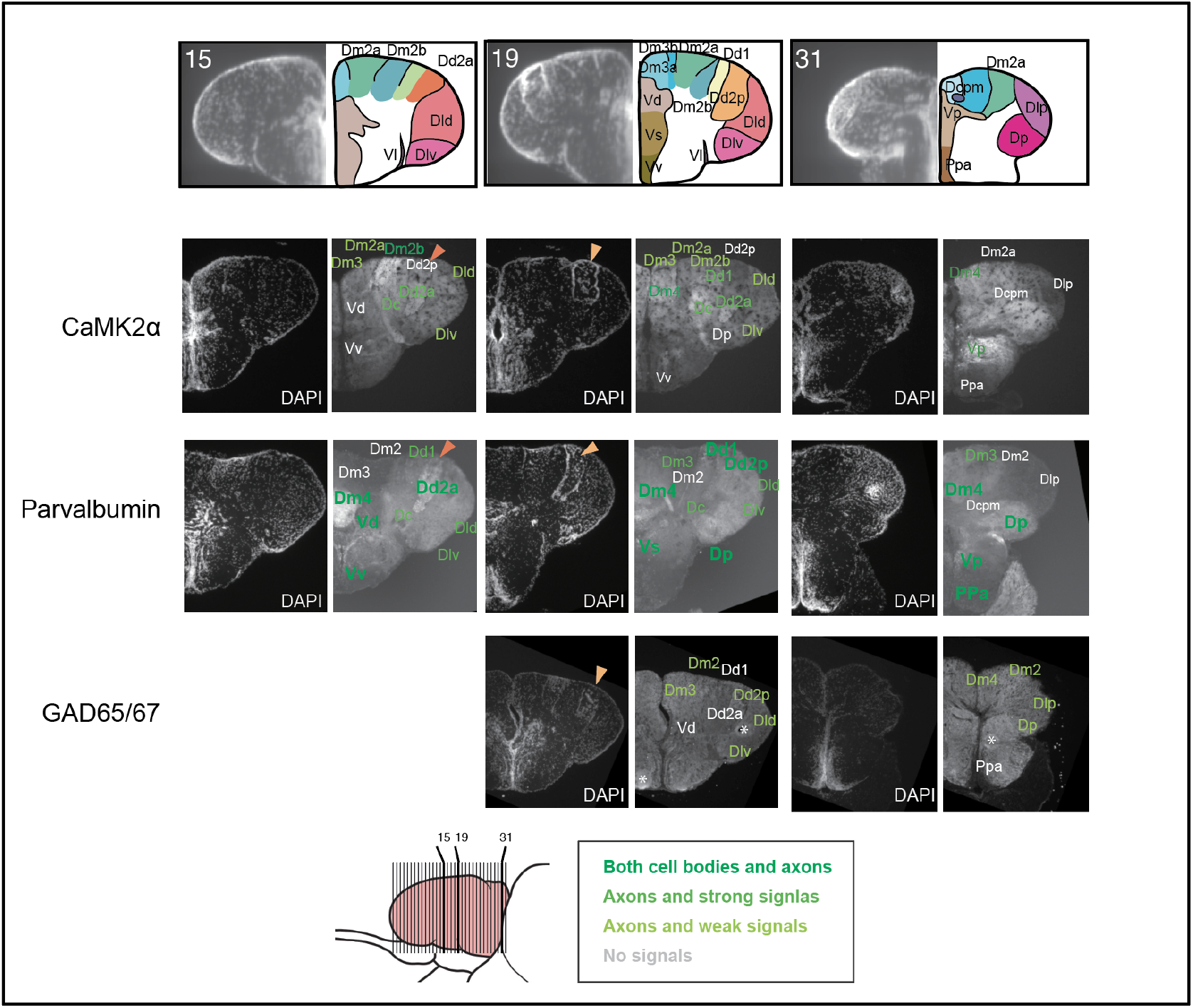
Expression of cell-type marker genes in the adult medaka telencephalon. Immunostaining of CaMK2α, Parvalbumin, and GAD65/67 allowed us to define anatomical regions. Numbers in the first row indicate the location of the transverse sections (bottom). Orange triangles indicate the position of Dd2 in medaka (dark orange:Dd2a, light orange:Dd2p). Shades of colors indicate the expression levels of the genes. Asterisk symbols indicate the noise from staining.

**Figure S3.**
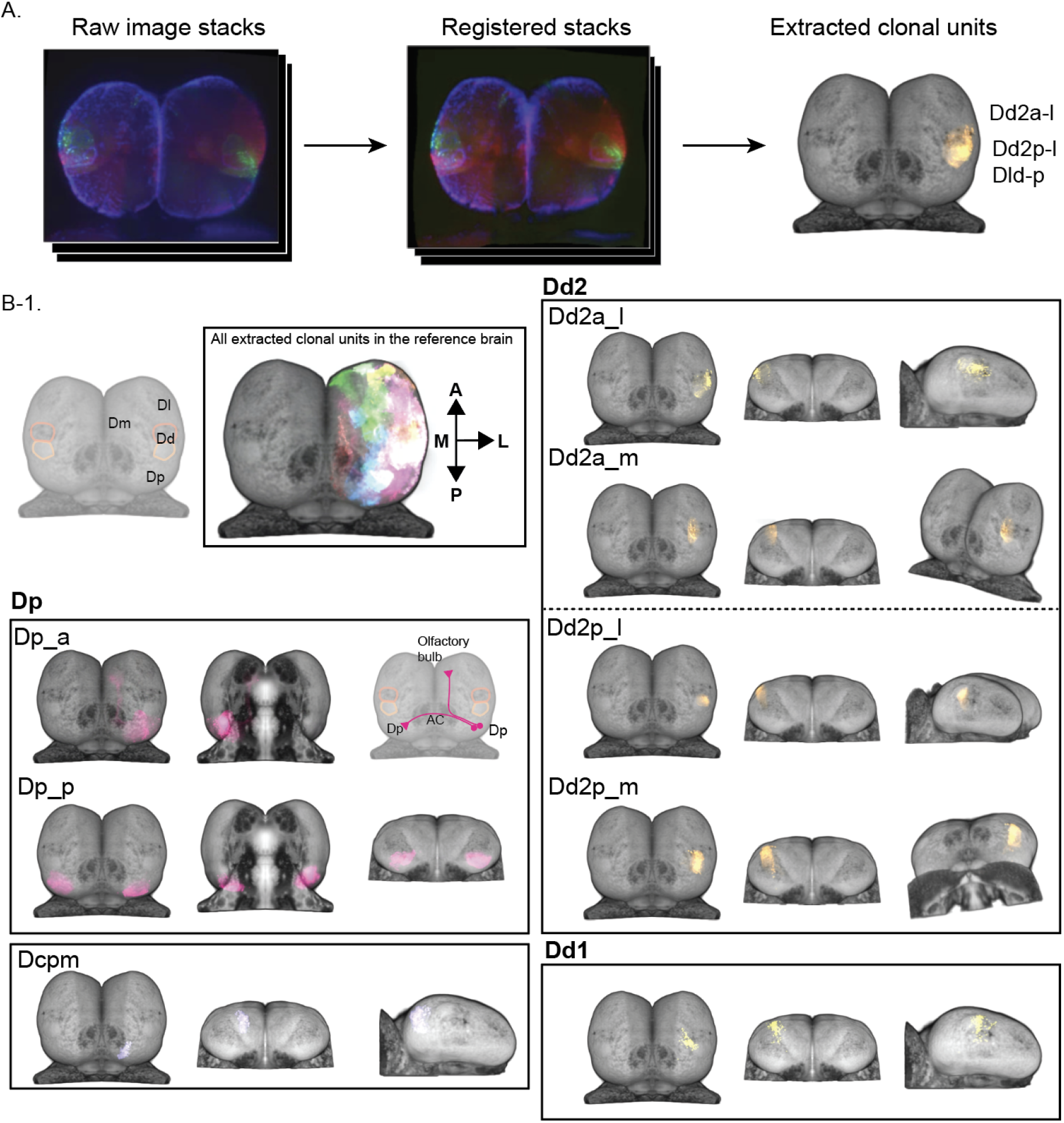
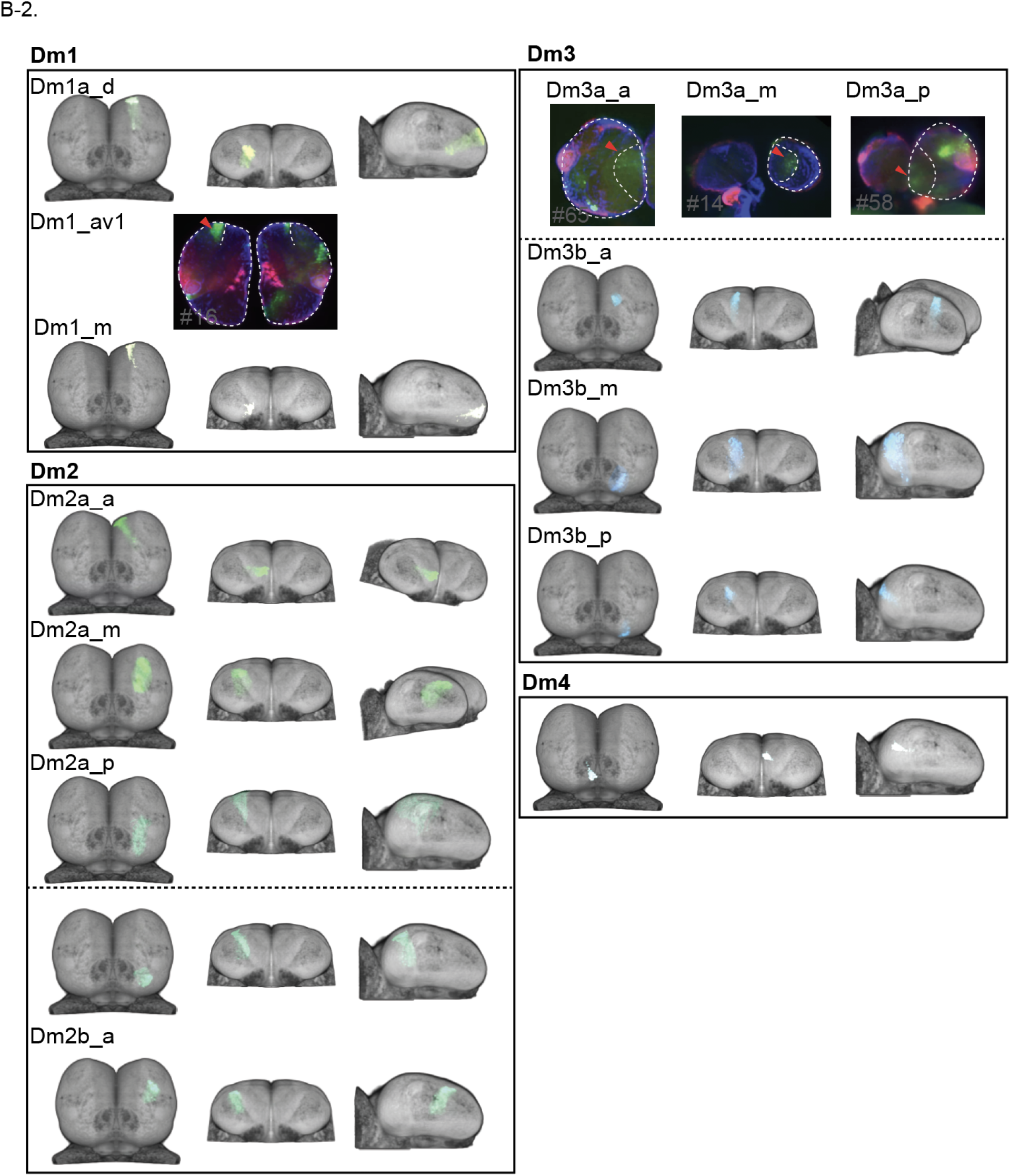
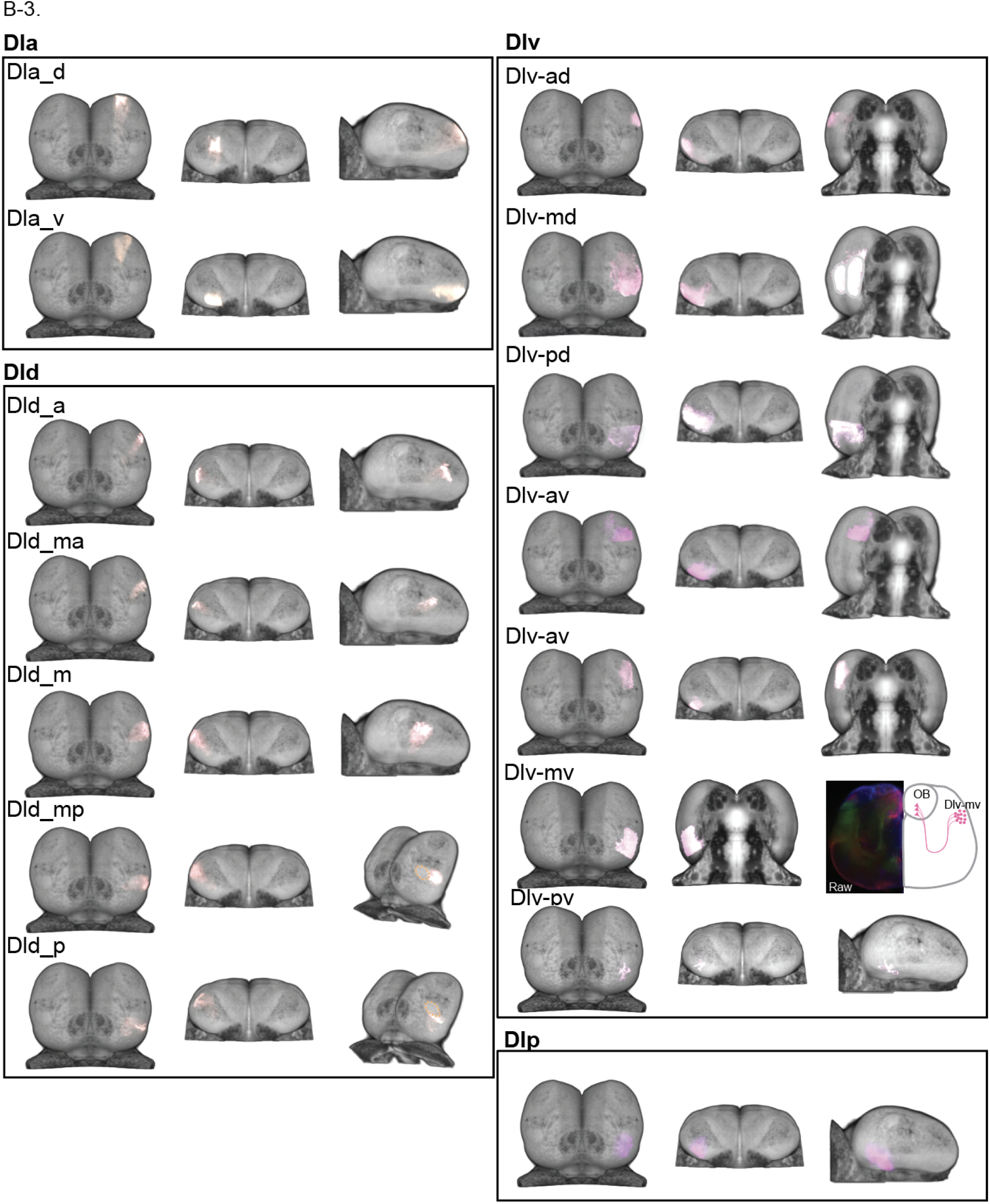
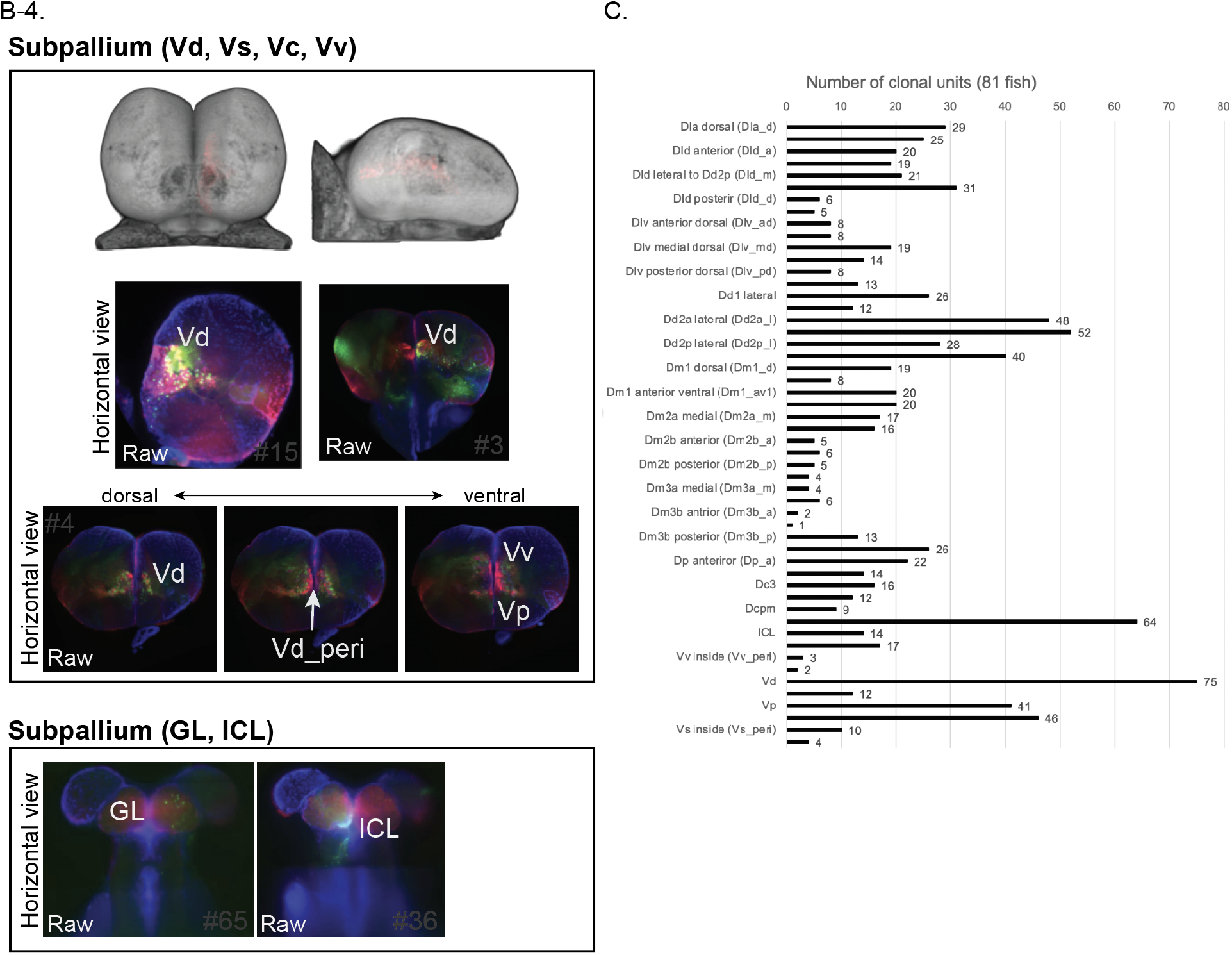
Clonal architecture in the adult medaka telencephalon. (A) Procedure of identifying the structure of clonal units. Raw image stacks were registered to the reference telencephalon with DAPI signals. Then GFP-positive cells were extracted. When GFP-positive cells were observed repeatedly, we identified the population as a clonal unit. (B) Detailed structural description of clonal units. All extracted clonal units are displayed (top left). Clonal units in Dd, Dp (B-1), Dm (B-2), Dl (B-3) and subpallium (B-4) are shown. Locations of the soma and neural axon terminal are indicated by circles and triangles (bottom). For the pallial clonal units, each panel shows a dorsal view (left column), front view (middle column) and different point of view (right column) of clonal units registered in the reference brain. Some of the clonal units were not able to be registered, so we show raw images. In the subpallium panel, registered clonal units (top) and raw data (bottom) are shown. GFP-positive cells and DsRed-positive cells were mixed in the subpallium. (C) Number of clonal units found in each brain anatomical region. We detected 561 GFP-positive neural populations from 81 fish. We found fewer GFP-positive populations in Dm2b and Dm3a. Since we gave mild heat shock to induce Cre-lox recombination by thermal cycler, it could be possible that the embryos tend to stay in a certain posture in the PCR tubes so that a few neural stem cells are not likely to get heat shock, which led to this bias.

**Figure S4.**
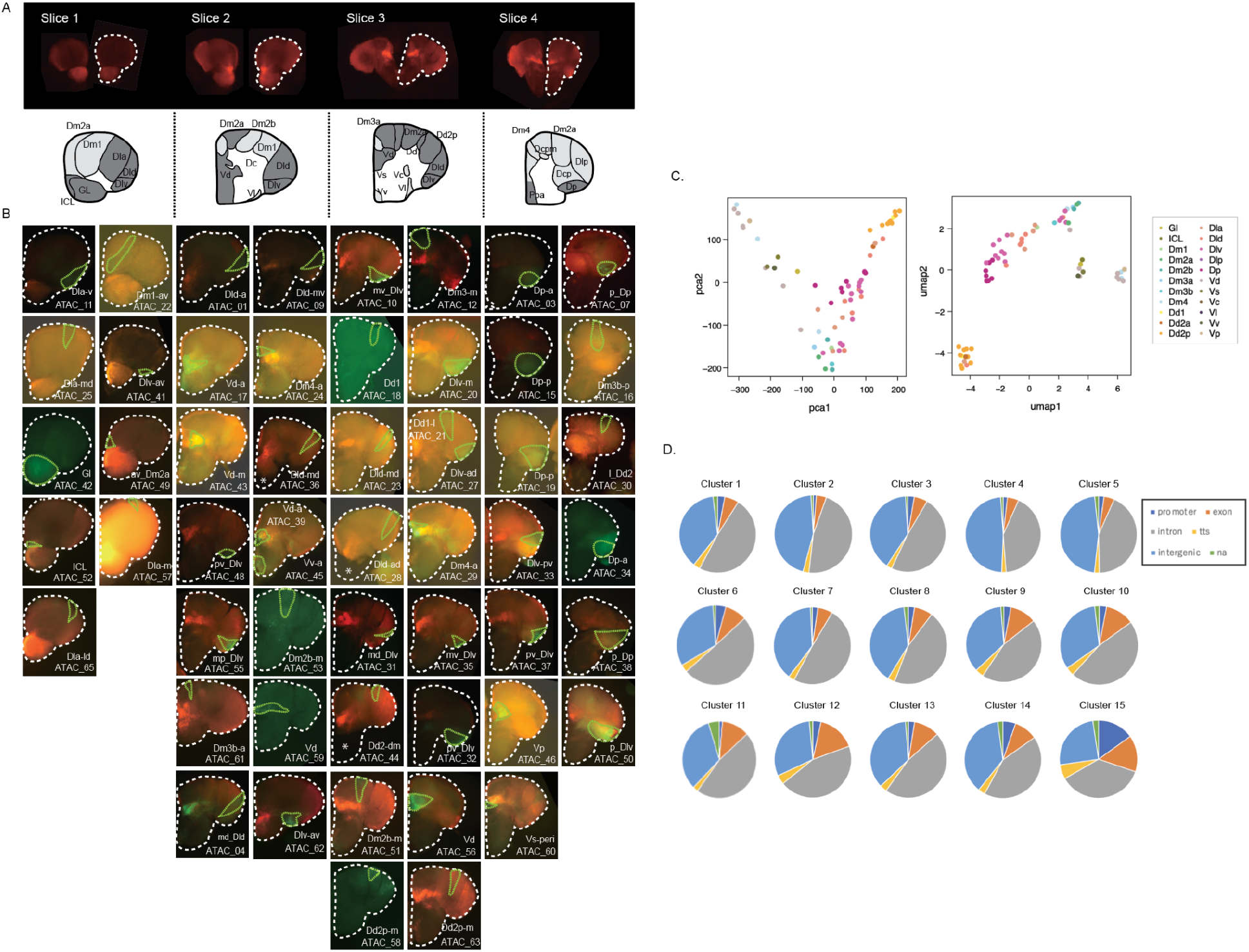
Analysis of ATAC-seq in clonal units. (A) 130 um-thick brain transverse sections from one adult brain (top). Clonal units used in ATAC-seq are shaded in dark gray in the schematic drawing (bottom). (B) Pictures of extracted GFP-positive clonal units (region surrounded by green dotted line). White dotted line shows the outline of brain slices. (C) Dimensionality reduction analysis of ATAC-seq peak patterns of clonal units using principal component analysis (PCA) (left) and Uniform Manifold Approximation and Projection (UMAP) analysis (right). Color indicates the location of clonal units in the anatomical regions. All dimensionality reduction analyses show that the subpallial clonal units are distinct from the pallial clonal units and that clonal units in Dd2 contain a unique open chromatin structure from other pallial clonal units. (D) Pie charts show the genomic distribution of the OCR clusters. OCR C15, which was identified as common open peaks among the telencephalon, includes more peaks in promoter regions.

**Figure S5.**
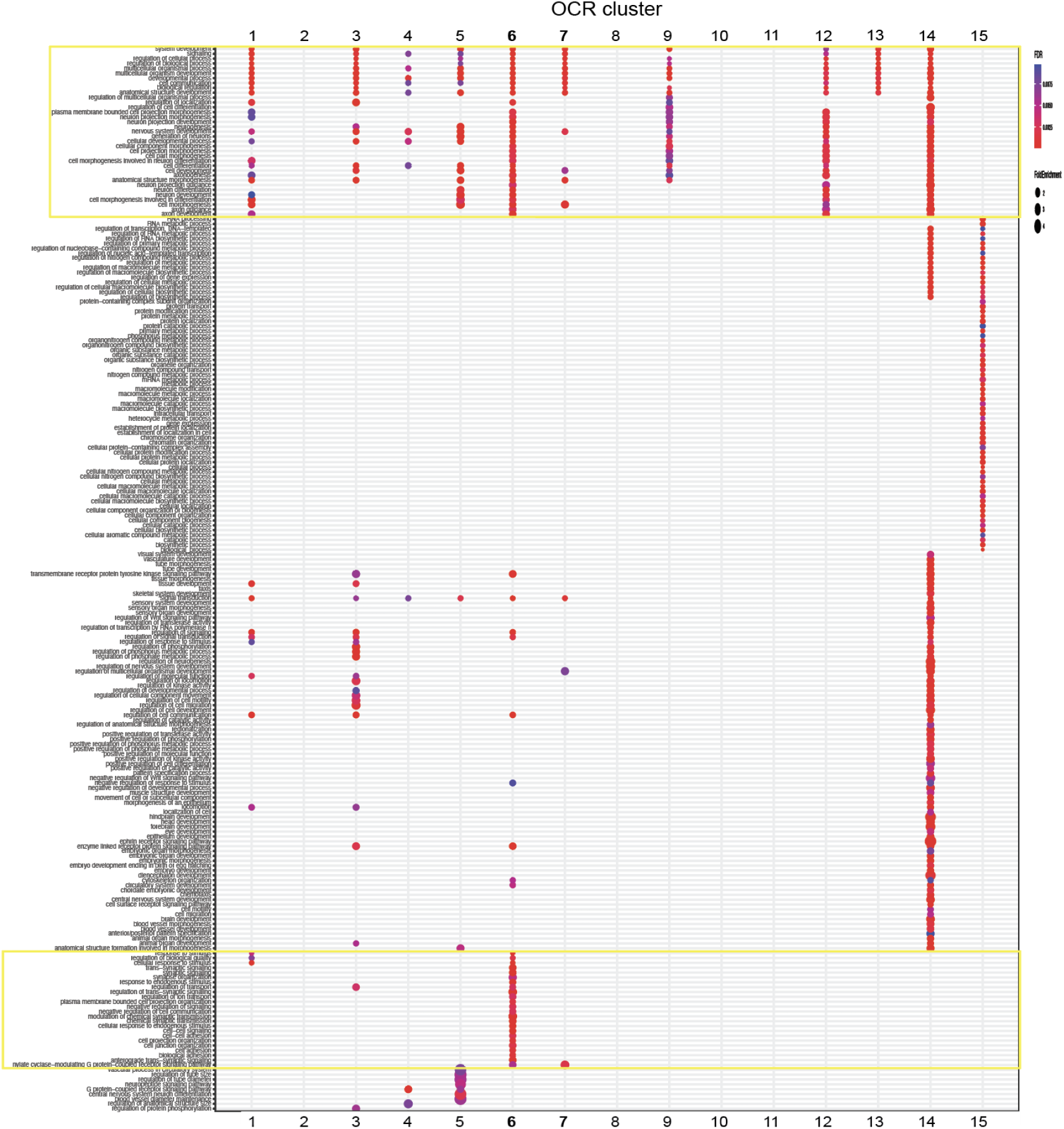
Gene ontology term enrichment analysis on OCR Clusters. GO term enrichment analysis was performed on OCR clusters in Figure 3D. In OCR C8, 10, 11, no GO term was enriched. Size and color of circles indicate fold enrichment and FDR, respectively. Circle was plotted only if FDR is lower than 0.01. Yellow rectangle indicates the GO terms related to axon projection pathways (top) and synapse organization (bottom), being focused in Figure 4A. OCR C14, 15 were identified as common open peaks among the telencephalon, which included many metabolic and biosynthesis processes (OCR C 14), and developmental signaling pathways that might reflect the adult neurogenesis process (OCR C 15). Also, one of the subpallial OCR clusters (OCR C5) was enriched with neuropeptide signaling, neural differentiation and vascular processes, which is consistent with previous avian studies that show neuropeptide expression in the subpallium ^70^.

**Figure S6.**
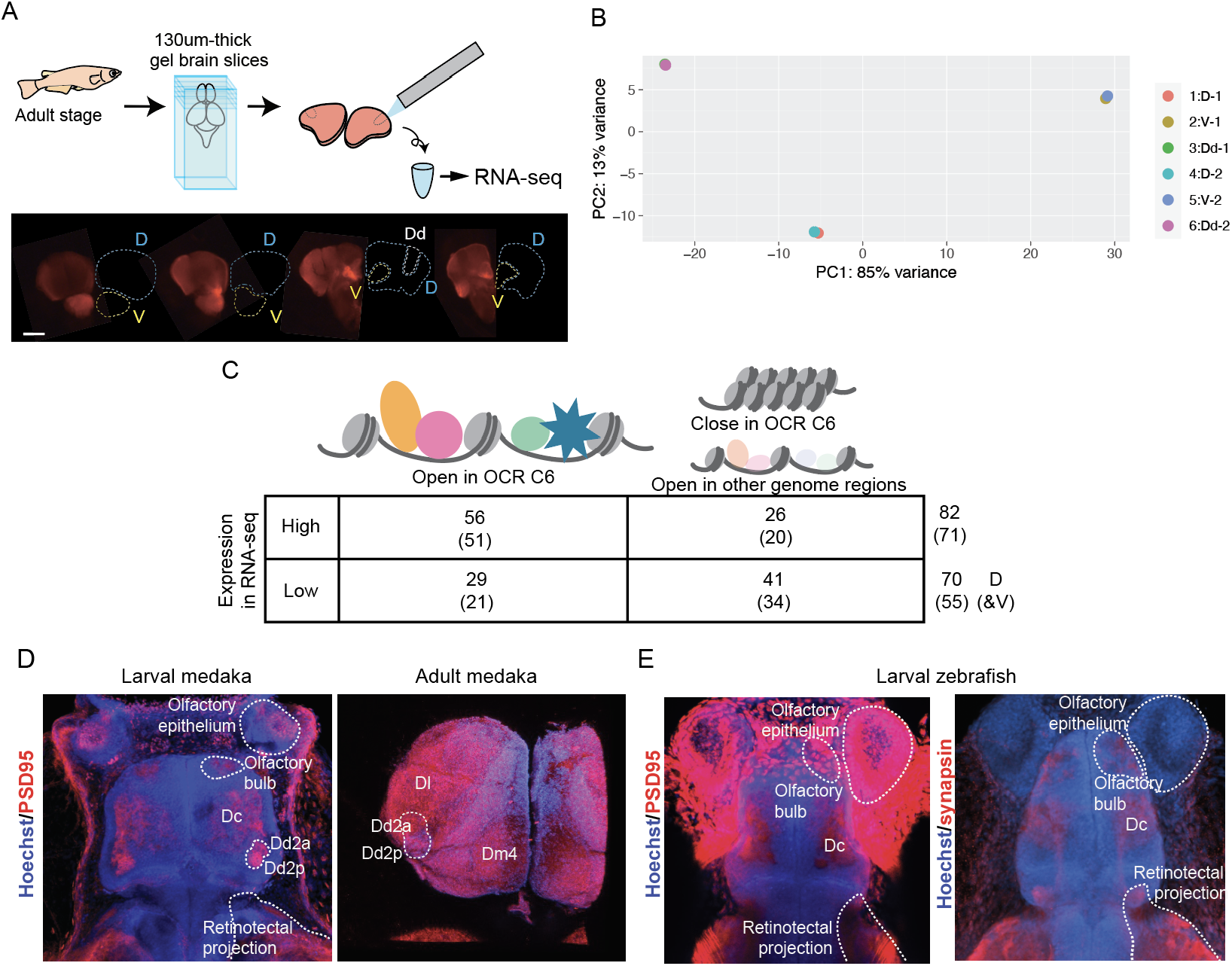
RNA-seq and ATAC-seq analysis on Dd2. (A) Procedure for RNA-seq analysis. After making 130 um-thick slices, Dd2 (Dd, white lines), the pallium except Dd (D, Cyan lines) and subpallium (V, yellow lines) were dissected based on the DsRed signals in the Tg (HuC-DsRed) line. Scale bar: 100um. (B) PCA analysis of the result of RNA-sequencing (two replicates for each sample). (C) Synaptic related genes differentially expressed and actively regulated in Dd2 from other regions. The number of synaptic genes with significantly high or low expression in the sample Dd2 detected by RNA-seq, and the number of synaptic-related genes targeted by OCR C6 are shown. By ATAC-seq, we identified genes actively down-regulated. (D) Maximum projection image of anti-PSD95 immunostaining on larval medaka (left) and dissected adult medaka (right) telencephalon. Blue: Hoechst signal, red: PSD95 signal. In larvae, PSD95 signals were detected in olfactory epithelium, olfactory bulb, Dc, retinotectal projection, and both Dd2a and Dd2p. Noisy signals were detected in the skin. In the adult telencephalon, PSD95 signals were broadly detected but we observed strong signals in Dl, Dd2a and Dm4. (E) Maximum projection image of anti-synapsin immunostaining on larval zebrafish. Both PSD95 and synapsin were detected in the olfactory bulb, Dc and retinotectal projection. Noisy signals were detected in the skin in anti-PSD95 immunostaining. No strong signal in the dorsal telencephalon was observed.

**Figure S7.**
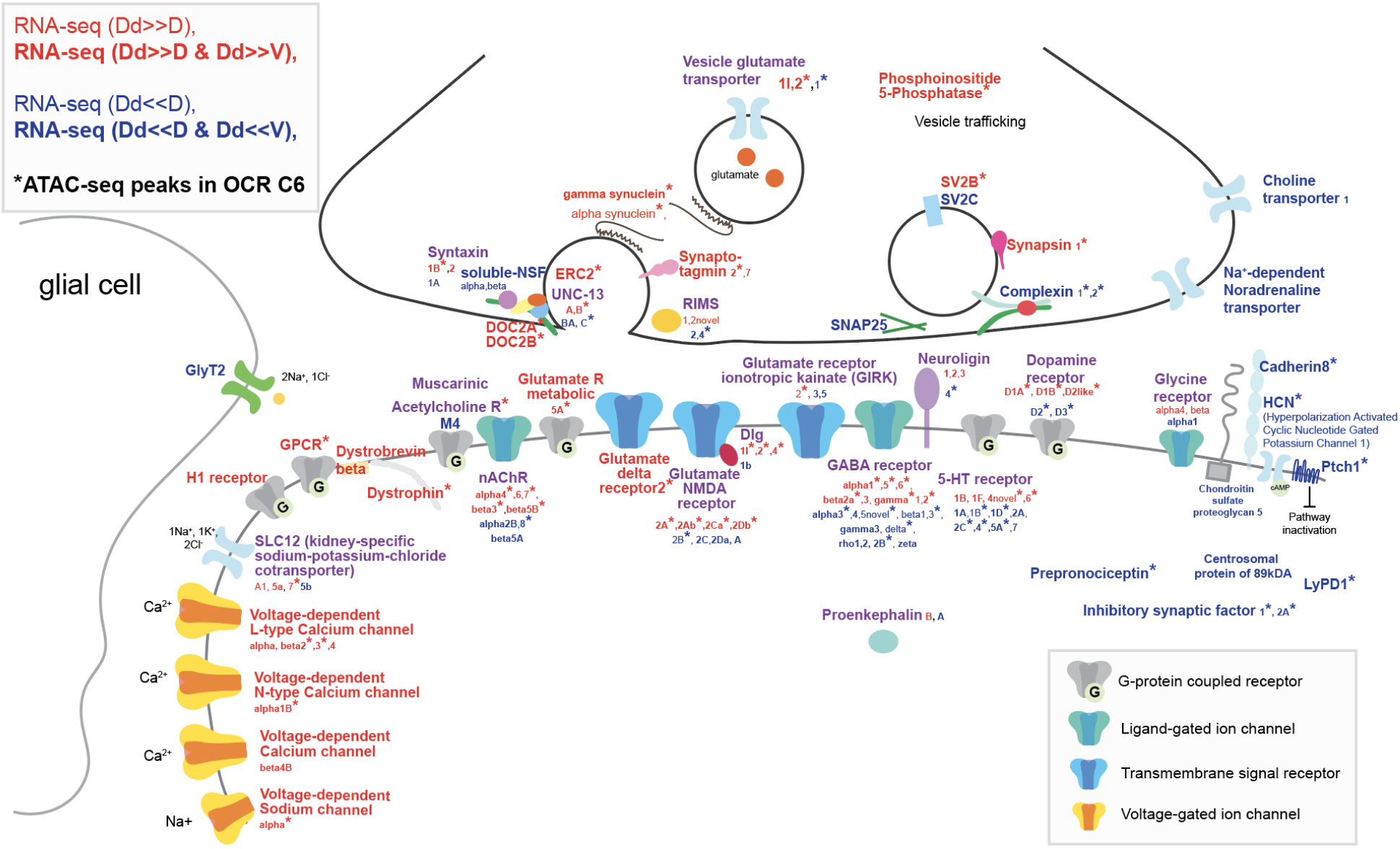
Characterization of synaptic genes expressed and regulated in Dd2. Schematic representation of synaptic genes detected as specifically expressed and regulated in Dd2 by ATAC-seq and RNA-seq. We found that different subunits of neurotransmitter receptors and ion channels are both up- and down-regulated in Dd2 compared to other pallial regions; excitatory glutamate receptors and glutamate transporters; modulatory neuronal genes, such as 5-HT receptors,cholinergic receptors and dopamine receptors; inhibitory receptors, such as GABAergic receptors. On the other hand, the voltage-dependent calcium channels expressed significantly higher in Dd2 than other regions, and the expression of inhibitory synaptic genes was suppressed. Red gene names indicate significantly highly expressing genes, blue gene names indicate actively significantly lowly expressing genes, and purple gene names indicate that subunits of the genes were differently expressed in Dd2. * indicates the genes targeted by Dd2-specific OCR C 6.

**Figure S8.**
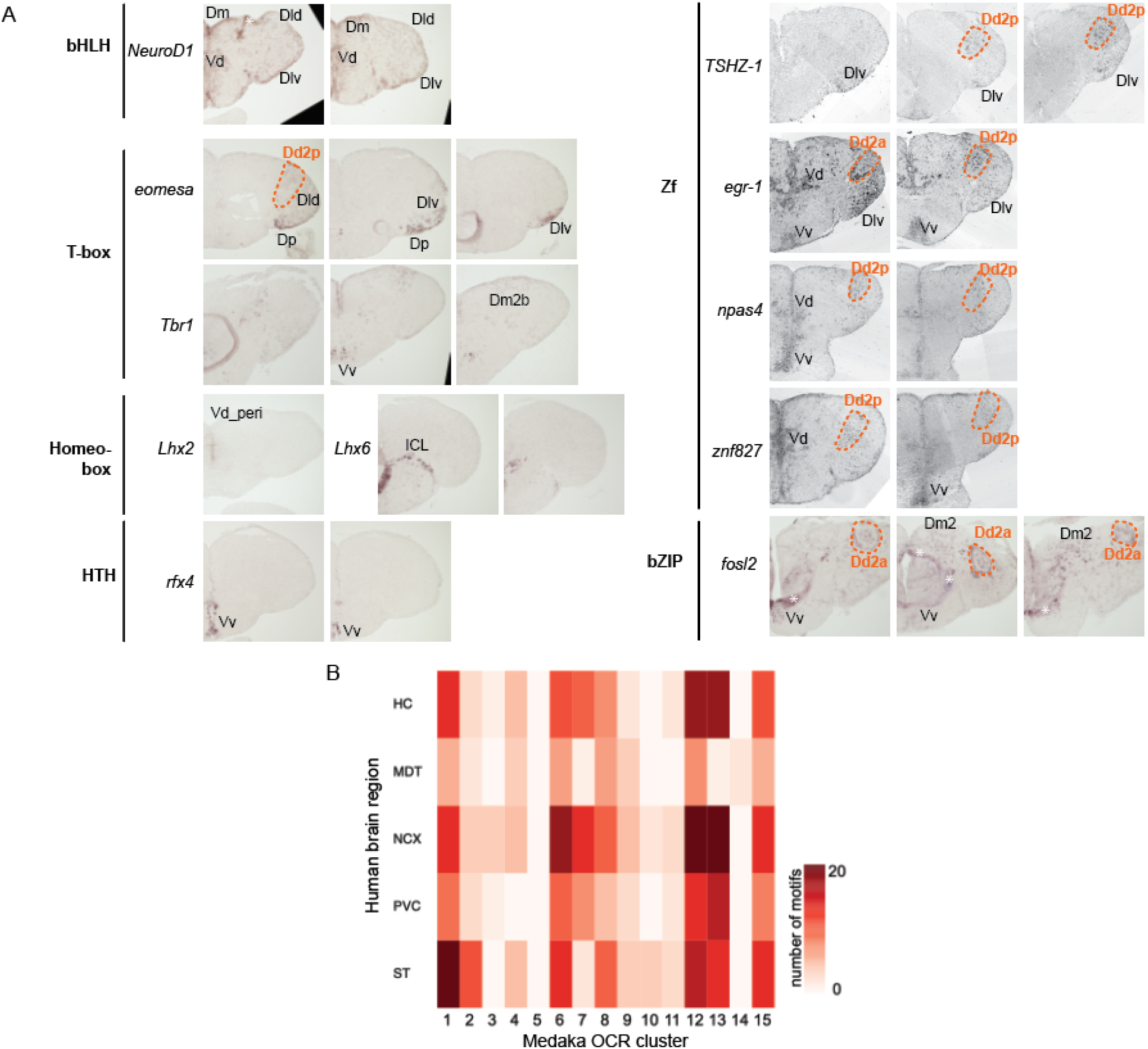
Expression of candidate transcription factors in the adult medaka telencephalon. (A) Brain regions where RNA signals were detected by *in situ* hybridization are indicated by names. Expression in Dd2 is shown with the orange line and name. bHLH: beta-helix-loop-helix family, HTH: helix-turn-helix family, Zf: zinc-finger family, bZIP: basic leucine zipper family. (B) A heatmap showing the number of shared motifs between human brain regional ATAC-seq peaks and medaka OCR clusters. HC: hippocampus, MDT: medial dorsal thalamus, NCX: neocortex, PVC: primary visual cortex, ST: striatum. Darker shade indicates larger overlap between medaka and human.

## Supplementary material

### 1. Nomenclature of the adult medaka telencephalon

For the medaka brain research, two brain atlases of adult medaka have been used ^18, 25^. Because performing three-dimensional reconstruction from brain slices is difficult, some inconsistencies in nomenclature were observed among references. To overcome this difficulty, we performed a three dimensional imaging of the nuclei-stained adult medaka telencephalon, which allowed us to analyze the anatomical boundaries in more accuracy. The number in the name of anatomical regions (such as Dm1, Dm2, Dm3) was defined by the order of the emergence in the anterior to posterior axis. Also, we performed immunostaining of marker genes (CaMK2α, parvalbumin, and GAD67) and examined the expression of a Tg line (HuC: DsRed) to confirm the homology of anatomical regions to other teleost species.

#### Pallium

##### - Dc

The dorso-central telencephalon (Dc) is defined as the center of the dorsal pallium in teleost ^2^. In most teleostean species, cells in Dc have larger size of cell body (Cichlid fish:^71^, sea bass: ^72^) and are sparsely distributed. According to the medaka brain atlas of Ishikawa et al ^25^., Dc is defined in the center of the pallium as well. However, in the other brain atlas of medaka fish^18^, multiple subregions are separately defined as Dcs. In our atlas, we defined the center region of the pallium that has less dense cell populations as Dc, and defined the aggregates of cells in the posterior medial center of dorsal pallium as Dcpm.

In zebrafish, *Danio rerio*, parvalbumin-positive axons project to Dc ^12^. In gymnotiform fish pallium, CaMK2α is widely expressed in Dc ^73^. In medaka pallium, both CaMK2α and parvalbumin are expressed in Dc, but we observe weak GAD-positive signals in the same area, which is consistent with other species.

Since the cell bodies in Dc of other teleosts are bigger than those of other anatomical regions, mature neurons are supposed to locate in Dc. In a big-sized brain, the matured neurons locate sparsely, which exhibit a clear anatomical region, Dc. However, as the size of medaka fish brain is not so big, it is difficult to find a clear region for Dc. According to the marker-gene expression patterns, we may name the region ventral to Dd1 and Dm2, the Dc.

##### Dcpm

The posterior medial nucleus of the dorso-central telencephalon (Dcpm) is surrounded by Dm4 and Dm3b. Although this region is not clearly defined by other previous papers, the cell density is higher than Dm3b and you can find it easily in the horizontal sections. In our immunostaining experiments, we didn’t observe CaMK2α, parvalbumin, nor GAD signals.

##### - Dm

The medial part of the dorsal pallium (Dm) contains small cell-body neurons with high density. In the horizontal optic sections, we found several linearly aligned cells on the dorsal surface of the telencephalic hemispheres which correspond to the boundaries of Dm subregions. These boundaries can be good landmarks to define the regions. From anterior to posterior, we named Dm1, Dm2, Dm3 and Dm4. We also found a clear boundary inside Dm2 and Dm3 so that we named them with a and b. We didn’t name them as different sub-regions because the gene-marker expressions were consistent in Dm2 and Dm3.

Dm1 region is located at the most anterior part of the telencephalon. It is densely packed with small cells. Dm2 locates posterior to the Dm1 region, Dm2 is packed with cells more densely than Dm1. In the horizontal sections, it is clear that Dm2 is located from anterior-medial to posterior-lateral part of the telencephalon. Dm3 located in the medial and posterior part of the hemisphere. Dm4 located dorsal to Vd and ventral to Dm3.

In zebrafish, parvalbumin is not expressed in Dm. Also CaMK2α is not expressed in gymnotiform fish. In medaka, Dm can be subdivided depending on the expression of CaMK2α and parvalbumin; in Dm1, weak CaMK2α expression but no parvalbumin expression were observed; in Dm2, weak CaMK2α expression in Dm2a, strong CaMK2α expression in Dm2b in cell bodies, but rare parvalbumin expression were observed; in Dm3, strong CaMK2α signals were observed in cell bodies, and parvalbumin positive fibers were observed; in Dm4, weak CaMK2α and strong parvalbumin expression were observed in cell bodies.

##### - Dl

In the previous medaka brain atlas, the anterior region of dorsal lateral pallium is simply named as Dl ^18^. But here we named this anterior part of dorso-lateral telencephalon the Dla, because the nuclear density is less than Dld, Dlv and Dlp. We also divided Dl into the dorsal and ventral part, since the nuclear density is different between the dorsal and ventral parts. We observed higher DAPI signal density in Dlp than Dld and Dlv.

In gymnotiform fish, both CaMK2α and GAD are expressed in Dl. In zebrafish, GAD67 was sparsely detected, and strong parvalbumin signals was detected in Dl ^12^. In medaka, genes were expressed uniformly in Dla, Dld and Dlv. Mild expression of CaMK2α, parvalbumin, and GAD were detected in these three subdivisions. On the other hand, in Dlp, neither CaMK2α and parvalbumin was expressed. GAD was weakly expressed in Dlp.

It is not easy to draw a line between the anterior part of the dorso-lateral telencephalon (Dla) and the posterior part of the dorso-lateral telencephalon (Dld) from sagittal sections. But in the horizontal sections, we found that the density of nucleuses are more sparse in Dla than Dld and Dlv. Dla region located lateral to the olfactory bulb and anterior to Dm1, Dld and Dlv.

The dorsal part of the middle and posterior part of the dorso-lateral telencephalon (Dld)^18^ is next to Dla and the cell-density is more than that of Dla. The boundary between Dld and Dlv is not clear. But the cells are more densely distributed in Dlv than Dld.

The ventral part of the middle and posterior part of the dorso-lateral telencephalon (Dlv)^18^ is highly packed with cells.

There is no clear boundary among the posterior regions of the dorso-lateral telencephalon (Dlp), the posterior of dorsal telencephalon (Dp), Dld, and Dlv. But we observed that Dlp was highly dense with a small nucleus and the cell distribution pattern was different from neighboring regions. Dlp is located posterior to Dld and Dlv. Also it is located dorsal to Dp.

##### - Dd

The dorso-dorsal telencephalon (Dd) is the region which is located in the dorsal part of the pallium. In the previous report^18^, Dd is subdivided into two regions, Dd1 and Dd2. In the other report ^25^, only Dd is defined which corresponds to Dd2 of Ralf H Anken, 1998^18^. Since the boundary between Dd1 and Dd2 was visible in DAPI staining, we followed the definition of Ralf H Anken, 1998^18^.

Dd2 is clearly demarcated in the telencephalon. As observed with the horizontal sections, Dd2 can be subdivided into 2 regions, Dd2a (anterior part of Dd2) and Dd2p (posterior part of Dd2). Dd2p is surrounded by cells, and the cell density is relatively higher than Dd2a. The boundary between Dd1 and Dm2a is clear. Dd1 is located medial to Dd2a and Dd2p.

In gymnotiform fish, DDi (intermediate subdivision of the dorsodorsal telencephalon) and DDmg (magnocellular subdivision of the dorsodorsal telencephalon) express GAD, but don’t express CaMK2. In zebrafish, the existence of Dd region is still in debate. But previous papers suggest that central pallium (Dc) which is the parvalbumin positive and GAD negative region is suggested to be homologous to the mammalian dorsal pallium ^11, 12^.

In medaka, parvalbumin was strongly expressed in the cell bodies of the anterior part of Dd2 (Dd2a), and we detected weak signals in the posterior Dd2 (Dd2p). CaMK2α are expressed in Dd2a and Dd1. While GAD was expressed in Dd2p, it was not expressed in Dd2a. In Dd1, parvalbumin was expressed. CaMK2α was also expressed weakly, but GAD was not expressed.

##### - DP

The posterior part of dorsal telencephalon (Dp) ^18^ is remarkably denser with the cell nucleus than other posterior anatomical regions. Dp is located posterior to Dlv. In zebrafish, parvalbumin and GAD67 are not expressed in Dp. In medaka, the expression of CaMK2α was not strong, while the parvalbumin and GAD were strongly expressed.

#### Subpallium

In many references, there are some anatomical regions defined in the ventral telencephalic areas. However, the DAPI staining and whole-telencephalon imaging suggest that the ventral regions can only be separated by AC, and there was no structurally clear boundary between Vd, Vs and Vp.

##### Vd

Cells in the dorsal medial part of the subpallium are called Vd. We also observed some cell clusters that are located laterally and inside the pallium. Those cell clusters correspond to the Vc region in some references. But according to the cell density and some gene expression, we define those regions also as Vd. It is reported that Vd corresponds to the striatum in the mammalian brain.

In zebrafish, GAD67 and parvalbumin are expressed strongly in Vd. In medaka, we found some parvalbumin-positive cell bodies. But CaMK2α-positive cell bodies were not found, though weak signals (maybe axons) were detected. Also, GAD was weakly detected in Vd.

##### Vs

The supracommissural part of ventral telencephalon (Vs) is located ventral to Vd. But there is no clear boundary between Vs and Vd. In medaka, we found some parvalbumin-positive cell bodies. But CaMK2α-positive signals were weak.

##### Vv

In zebrafish, GAD67 is expressed strongly in the ventral part of the subpallium (Vv), but not parvalbumin signals. In medaka, parvalbumin-positive cells were sparsely detected. CaMK2α-positive signals were not detected.

##### Vl

The lateral part of the subpallium (Vl) located near Dlv. In zebrafish, GAD67 is expressed strongly in Vl but not parvalbumin signals. In medaka, strong signals of anti-parvalbumin were detected but not anti-CaMK2α signals.

#### Olfactory bulb

In the anterior part of the telencephalon, the external (ECL) and internal (ICL) cellular layer of the olfactory bulb and the glomerular layer of the olfactory bulb (GL) are clearly found. In zebrafish, parvalbumin is strongly expressed in the ventral layer of the olfactory bulb. In zebrafish, parvalbumin is not detected in ICL and ECL

### 2. Structure of clonal units

Here we describe the structure of clonal units visualized by the stochastic Cre-loxp recombination induced in the transgenic line (HSP:Cre x HuC:loxp-DsRed-loxp-GFP). We observed the GFP and DsRed signals in cell somas, dendrites and axons. DAPI staining allowed us to visualize the anatomical boundaries in the telencephalon (See also Figure S3).

#### Pallium

When we observed GFP-positive neurons in the pallium, those somas were mostly clustered and located inside the same anatomical boundaries.

##### - Dc

We found many axon bundles projecting through the Dc region, but we didn’t observe any GFP-positive cell soma just located inside the Dc region. In the teleost pallium, neural stem cells exist on the surface of the hemisphere. So, it is possible that cells in the Dc region are mature cells and that’s why they were not visualized with HuC promoter.

##### - Dm

In general, GFP-positive cells visualized in the Dm project axons from the dorsal to ventral direction toward either Dc, Dd or subpallial regions.

##### Dm1

We found that cells in Dm1 project their axons inside Dm1 from anterior to posterior direction. We identified the structure of three clonal units in Dm1 (Dm1-ad, Dm1-av1, Dm1-m), but these didn’t cover the whole Dm1 regions.

##### Dm2

3 clonal units (anterior, middle, posterior) clonal units were identified in both Dm2a and Dm2b. We found that the axons of cells in Dm2a projected into the Dc region and Vd, while the axons in Dm2b projected into the Dc region and Vd and Vp..

##### Dm3

3 clonal units (anterior, middle, posterior) clonal units were identified in both Dm3a and Dm3b. We found that the axons of cells in Dm3a projected into the Dc region and Vd, while the axons in Dm3b projected into the Dc region and Vd and Vv.

##### - Dl

In general, we observed GFP positive cell soma near the surface of the hemisphere and GFP-positive axon bundles project toward Dc, Dd or the olfactory bulb.

##### Dld

We found there are 5 clonal units in Dld; one in the anterior part (Dld_a), one in the middle anterior part (Dld_ma), one in the middle part (Dld_m), one in the middle posterior part (Dld_mp), and the one in the posterior part (Dld_p). Dld_mp was also located lateral to the Dd2p and projected its axons into the Dd2p. Dld_p was located posterior to the Dd2p and projected its axons into the Dd2p. Clonal units in Dld projected to Dc and surrounding regions

##### Dlv

We found that Dlv has two layers (dorsal and ventral). In general, the clonal units in the dorsal part of Dlv (Dlv_d) project their axons to Dc from lateral to medial direction, while the clonal unit in the ventral part of Dlv (Dlv_v) project their axons from lateral to medial direction and project into the olfactory bulb.

##### Dlp

We found cells in Dlp clustered near the surface of the brain. Long axonal projection was not observed.

##### - Dd

###### Dd1

We found that cells project to Dc and Dd2.

###### Dd2

We found that clonal units in the Dd2a and Dd2p project differently: cells in Dd2a project not only inside Dd2a but also to Dm and Dc, while cells in Dd2p project to Dd2a.

##### - Dp

Two distant axonal projections were observed. Cells whose soma located the anterior part of Dp projected axons into the ipsilateral olfactory bulb. On the other hand, cells whose soma located the posterior part of Dp projected axons into the contralateral Dp regions through anterior commissure (AC). We also observed an axonal bundle projected to the midbrain from Dp regions.

#### Subpallium

In general, we found that GFP- and DsRed-positive cells mixed inside the subpallial region. Also we observed that the axons of suballium are shorter than those of pallium and we didn’t observe axonal bundles. That’s why we couldn’t identify the independent structure of clonal units in the subpallium.

##### - Vd, Vs, Vp and Vv

We observed the mixture of GFP- and RFP-positive cells in the subpallium throughout the anterior to posterior direction. Cells in the Vd project their axons into Dm, Dc, and Dd. Also we observed GFP positive cells in the peripheral region of Vd and Dm4 (Vd_peri).

##### - Olfactory bultb (GL, ICL)

We observed GFP-positive cells distributed sparsely in GL and ICL.

