## Supplementary material for "Epigenetically distinct synaptic architecture in clonal compartments in the teleostean dorsal pallium": Table 2

| axonal terminal | #_141102_06_write_merge_2 | #_141102_06_write_merge_3 | #1_composite_8 | #1_composite_8 | #1_composite_8 | #1_composite_8 | #1_composite_8 | #1_composite_8 | #1_composite_8 | #1_composite_8 | #2_composite_2-1 | #2_composite_2-1 | #2_composite_2-1 | #2_composite_2-1 | #2_composite_2-1 | #2_composite_2-1 | #2_composite_2-1 | #2_composite_2-1 | #2_composite_2-1 | #2_composite_2-1 | #2_composite_2-1 | #2_composite_2-1 | #2_composite_2-1 | #2_composite_2-1 | #2_composite_2-1 | #2_composite_2-1 | #3_composite_4-1 | #3_composite_4-1 | #3_composite_4-1 | #3_composite_4-1 | #3_composite_4-1 | #3_composite_4-1 | #3_composite_4-1 | #3_composite_4-1 | #3_composite_4-1 | #3_composite_4-1 | #3_composite_4-1 | #3_composite_4-1 | #4_composite_5-1 | #4_composite_5-1 | #4_composite_5-1 | #4_composite_5-1 | #4_composite_5-1 |
|---|---|---|---|---|---|---|---|---|---|---|---|---|---|---|---|---|---|---|---|---|---|---|---|---|---|---|---|---|---|---|---|---|---|---|---|---|---|---|---|---|---|---|---|
| fish number | #1 | | 2 | | | | | | | | 3 | | | | | | | | | | | | | | | | 4 | | | | | | | | | | | | 5 | | | | |
| Serial number | 1 | 2 | 3 | 4 | 5 | 6 | 7 | 8 | 9 | 10 | 11 | 12 | 13 | 14 | 15 | 16 | 17 | 18 | 19 | 20 | 21 | 22 | 23 | 24 | 25 | 26 | 27 | 28 | 29 | 30 | 31 | 32 | 33 | 34 | 35 | 36 | 37 | 38 | 39 | 40 | 41 | 42 | 43 |
| clone number in the same fish | 1 | 2 | 1 | 2 | 3 | 4 | 5 | 6 | 7 | 8 | 1 | 2 | 3 | 4 | 5 | 6 | 7 | 8 | 9 | 10 | 11 | 12 | 13 | 14 | 15 | 16 | 1 | 2 | 3 | 4 | 5 | 6 | 7 | 8 | 9 | 10 | 11 | 12 | 1 | 2 | 3 | 4 | 5 |
| Left or Right hemisphere | R | R | L | L | L | R | R | R | R | R | L | L | L | L | L | L | L | L | R | R | R | R | R | R | R | R | L | L | L | L | R | R | R | R | R | R | R | R | L | L | L | L | L |
| Dla dorsal (Dla_d) | ○ | | | | | | | | | | | | | | | | | | | | | | | | | | | | | | | | | | | | ○ | | | | | | |
| Dla ventral (Dla_v) | | | | | | | | | | | | | | | ○ | | | | | | | | | | | | | | | | | | | | | | | ○ | | | | | |
| Dld anterior (Dld_a) | | ○ | | | ○ | | | | | | | | | | | | | | | | | | | | | | | | | | | | | | | | | | | | | | |
| Dld lateral to Dd2a (Dld_ma) | | | | | | | | | | | | | | | | | | | | | | | | | | | | | | | | | | | | | | | | | | | |
| Dld leteral to Dd2p (Dld_m) | | | | | | | ○ | | | | | | | | | | | | | | | | | | | | | | | | | | | | | | | | | | | | |
| Dld posterior to Dd2 (Dld_md) | | | | | | | | ○ | | | | ○ | | | | | | | | | | | | | | | | | | | | ○ | | | | | | | | | | | |
| Dld posterir (Dld_d) | | | | | | | | | | | | | | | | | | | | | | | | | | | | | | | | | | | | | | | | | | | |
| Dlp | | | | | | | | | | | | | | | | | | | | | | | | | | | | | | | | | | | | | | | | | | | |
| Dlv anterior dorsal (Dlv_ad) | | | | | | | | | | | | | | | | | | | | | | | | | | | | | | | | | | | | | | | | | | | |
| Dlv anterior ventral (Dlv_av) | | | | | | | | | | | | | | | | | | | | | | | | | | | | | | | | | | | | | | | | ○ | | | |
| Dlv medial dorsal (Dlv_md) | | | | | | | | | | | | | | | | | ○△ | | | | | | | | | ○ | | ○ | | | | | | | | | | | | | | | |
| Dlv medial ventral (Dlv_mv) | | | | | | | | | | | | | | | | | | | | | | | | | | | | | | | | | | | | ○ | | | | ○ | | | |
| Dlv posterior dorsal (Dlv_pd) | | | | | | | | | | | | | | | | | | | | | | | | | | | | | | | | | | | | | | | | | | | |
| Dlv poesterior ventral (Dlv_pv) | | | | | | | | | | | | | | | | | | | | | | | | | | | | | | | | | | | ○ | | | | | ○ | | | |
| Dd1 lateral | | | | | | | | | | | | | | | | | | | | | | | | | | | | | | | | | ○ | | | | | | | | | | |
| Dd1 medial | | | | | | | | | | | | | | | | | | | | | | | | | | | | | | | | | | | | | | | | | | | |
| Dd2a lateral (Dd2a_l) | | | △ | | | | | | | | △ | | | | | | | | △ | | | | | | | | ○△ | | | | | | | | | | | | | | | | |
| Dd2a medial (Dd2a_m) | | | △ | | | | | | | | △ | | | | | | | | △ | △ | | | | | | | ○△ | | | | | | | | | | | | | | | | |
| Dd2p lateral (Dd2p_l) | | | △ | | | | | | | | ○△ | | | | | | | | △ | | | | | | | | | | | | | | | | | | | | | | | | |
| Dd2p medial (Dd2p_m) | | | ○△ | | | | | | | | ○△ | | | | | | | | ○△ | △ | | | | | | | ○△ | | | | ○ | | | | | | | | | | | ○ | |
| Dm1 dorsal (Dm1_d) | | | | | | | | | | | | | | | | | | | | | | | | | ○ | | | | | | | | | | | | | | | | | | |
| Dm1 medial (Dm1_m) | | | | | | | | | | | | | | | | | | | | | | | | | | | | | | | | | | | | | | | | | | | |
| Dm1 anterior ventral (Dm1_av1) | | | | | | | | | | | | | | | | | | | | | | | | | | | | | | | | | | | | | | | | | | | |
| Dm2a anterior (Dm2a_a) | | | | | | | | | | | | | | | | | | | | | | | | | | | | | | | | | | | | | | | | | | | |
| Dm2a medial (Dm2a_m) | | | | | | ○ | | | | | | | | | | | | | | | | | | | | | | | | | | | | | | | | | | | | | |
| Dm2a posterior (Dm2a_p) | | | | | | | | | ○ | | | | | | | | | ○ | | | | | | | | | | | | | | | | | | | | | | | | | |
| Dm2b anterior (Dm2b_a) | | | | | | | | | | | | | | | | | | | | | | | | | | | | | | | | | | | | | | | | | | | |
| Dm2b medial (Dm2b_m) | | | | | | | | | | | | | | | | | | | | | | | | | | | | | | | | | | | | | | | | | | | |
| Dm2b posterior (Dm2b_p) | | | | | | | | | | | | | | | | | | | | | | | | | | | | | | | | | | | | | | | | | | | |
| Dm3a antrior (Dm3a_a) | | | | | | | | | | | | | | | | | | | | | | | | | | | | | | | | | | | | | | | | | | | |
| Dm3a medial (Dm3a_m) | | | | | | | | | | | | | | | | | | | | | | | | | | | | | | | | | | | | | | | | | | | |
| Dm3a posterior (Dm3a_p) | | | | | | | | | | | | | | | | | | | | | | | | | | | | | | | | | | | | | | | | | | | |
| Dm3b antrior (Dm3b_a) | | | | | | | | | | | | | | | | | | | | | | | | | | | | | | | | | | | | | | | | | | | |
| Dm3b medial (Dm3b_m) | | | | | | | | | | | | | | | | | | | | | | | | | | | | | | | | | | | | | | | | | | | |
| Dm3b posterior (Dm3b_p) | | | | | | | | | | | | | | | | | | | | | | | | | | | | | | | | | | | | | | | | | | | |
| Dm4 | | | | | | | | | | | | | | | | ○ | | | | | ○ | | | | | | | | | | | | | | | | | | | | ○ | | |
| Dp anteriror (Dp_a) | | | | | | | | | | | | | | | | | | | | | | ○ | | | | | | | | | | | | ○ | | | | | | | | | |
| Dp posterior (Dp0_p ) | | | | | | | | | | | | | ○ | | | | | | | | | | | | | | | | | | | | | | | | | | | | | | |
| Dc3 | | △ | | | △ | ○ | | | | | | | | | | | | △ | | | | | | | | | | | | | | | | | | | | | | | | | |
| Dc4 | | | | | | | | | | | | | | | | | | | | | | | | | | | | | | | | | | | | | | | | | | | |
| Dcpm | | | | | | | | | | ○ | | | | | | | | | | | | | | | | | | | | | | | | | | | | | | | | | |
| GL | | | | | | | | | | | | | | | | | | | | | | △ | ○ | | | | | | | | | | | △ | | | | | | △ | | | |
| ICL | | | | | | | | | | | | | | | | | | | | | | | | | | | | | | | | | | | | | | | | | | | |
| Vv | | | | | | | | | | | | | | | | | | | | | | | | ○ | | | | | | | | | | | | | | | | | | | |
| Vv inside (Vv_peri) | | | | | | | | | | | | | | | | | | | | | | | | | | | | | | | | | | | | | | | | | | | |
| Vc | | | | | | | | | | | | | | | | | | | | ○ | | | | | | | | | | | | | | | | | | | | | | | |
| Vd | | | | ○ | | | | | | | | | | ○ | | | | | | | | | | | | | | | ○ | | | | | | | | | | ○ | | | | |
| Vd inside (Vd_peri) | | | | | | | | | | | | | | | | | | | | | | | | | | | | | | | | | | | | | | | | | | | |
| Vp | | | | | | | | | | | | | | | | | | | | | | | | | | | | | | | | | | | | | | | ○ | | | | |
| Vs | | | | | | | | | | | | | | | | | | | | | | | | | | | | | | | | | | | | | | | ○ | | | | |
| Vs inside (Vs_peri) | | | | | | | | | | | | | | | | | | | | | | | | | | | | | | | | | | | | | | | | | | | |
| Vl | | | | | | | | | | | | | | | | | | | | | | | | | | | | | | | | | | | | | | | | | | | |

| #4_composite_5-1 | #4_composite_5-1 | #4_composite_5-1 | #4_composite_5-1 | #4_composite_5-1 | #4_composite_5-1 | #4_composite_5-1 | #5_composite_6-2 | #5_composite_6-2 | #5_composite_6-2 | #5_composite_6-2 | #5_composite_6-2 | #5_composite_6-2 | #6_composite_7-1 | #6_composite_7-1 | #6_composite_7-1 | #6_composite_7-1 | #6_composite_7-1 | #6_composite_7-1 | #7_140204_31 | #8_140204_32_stit ch_rgb_1-1 | #8_140204_32_stit ch_rgb_1-1 | #8_140204_32_stit ch_rgb_1-1 | #8_140204_32_stit ch_rgb_1-1 | #8_140204_32_stit ch_rgb_1-1 |
|---|---|---|---|---|---|---|---|---|---|---|---|---|---|---|---|---|---|---|---|---|---|---|---|---|
| | | | | | | | 6 | | | | | | 7 | | | | | | 8 | 9 | | | | |
| 44 | 45 | 46 | 47 | 48 | 49 | 50 | 51 | 52 | 53 | 54 | 55 | 56 | 57 | 58 | 59 | 60 | 61 | 62 | 63 | 64 | 65 | 66 | 67 | 68 |
| 6 | 7 | 8 | 9 | 10 | 11 | 12 | 1 | 2 | 3 | 4 | 5 | 6 | 1 | 2 | 3 | 4 | 5 | 6 | 1 | 1 | 2 | 3 | 4 | 5 |
| L | R | R | R | R | R | R | L | R | R | R | R | R | L | R | R | R | R | R | R | L | L | L | L | L |
| | | | | | | | | | | | | | | | | | | | | | | | ○ | |
| | | | | | | | ○ | | | | | | | | | | | | | | | | | |
| | | | | | | | ○ | | | | | | | | | | | | ○ | | | | | |
| | | | | | | | | | | | | | | | | | | | | | | | | ○ |
| | | | ○ | | | | | | | | ○ | | | | | | ○ | | | | | | | |
| ○ | | | | | | | | | | | | | | | | | | | | | | | | |
| | | | | | | ○ | | | | | | | | | | | | | | | | | | |
| | | | | | | ○ | | | | | | | | | | | | | | | | | | |
| | | | | | | | | | | ○ | | | | | | | | | | | | | | |
| | | | | | | | | | △ | | | | | | | | | | | | | | | |
| | | | | | | | | | △ | | | | | | | | | | | | | ○ | | |
| | | | | | | | | | ○ | | | | | | | | | | | | | | | |
| | | | | | ○ | | | | | | | | | | | | | | | | | | | |
| | | | | | | | | | | | | | | | | | | ○ | | | | | | |
| | | | | | | | | | | | | | | | | | | | | | ○ | | | |
| | | | | ○ | | | | ○ | | | | | | | | | | | | | | | | |
| | | | | | | | | | | | | | | ○ | | ○ | | | | | | | | |
| | | | | | | | | | | | | | | | | ○ | | | | | | | | |
| | △ | | | | | | | | | | | | | | | | | | | | | | | |
| | △ | | | | | | | | | | | | | | | | | | | | | | | |
| | | ○ | | | | | | | | | | | | | | △ | | | | | | | | |
| | | | | | | | | | | | | | ○ | | | | | | | | | | | |
| | ○ | | | | | | | ○ | | | | | ○ | | | | | | | ○ | | | | |
| | | | | | | | | | | | | | ○ | | | | | | | | | | | |
| | ○ | | | | | | | | | | | | | | | | | | | | | | | |
| | ○ | | | | | | | | | | | | | | | | | | | ○ | | | | |
| | | | | | | | | | | | ○ | | | | | | | | | | | | | |

| #8_140204_32_stit ch_rgb_1-1 | #8_140204_32_stit ch_rgb_1-1 | #8_140204_32_stit ch_rgb_1-1 | #8_140204_32_stit ch_rgb_1-1 | #8_140204_32_stit ch_rgb_1-1 | #8_140204_32_stit ch_rgb_1-1 | #9_140204_36_stit ch_rgb_big-2 | #9_140204_36_stit ch_rgb_big-2 | #9_140204_36_stit ch_rgb_big-2 | #9_140204_36_stit ch_rgb_big-2 | #9_140204_36_stit ch_rgb_big-2 | #9_140204_36_stit ch_rgb_big-2 | #9_140204_36_stit ch_rgb_big-2 | #10_140204_37_sti tch_rgb_big-2 | #10_140204_37_sti tch_rgb_big-2 | #10_140204_37_sti tch_rgb_big-2 | #10_140204_37_sti tch_rgb_big-2 | #10_140204_37_sti tch_rgb_big-2 | #10_140204_37_sti tch_rgb_big-2 | #10_140204_37_sti tch_rgb_big-2 | #10_140204_37_sti tch_rgb_big-2 | #10_140204_37_sti tch_rgb_big-2 | #10_140204_37_sti tch_rgb_big-2 | #10_140204_37_sti tch_rgb_big-2 | #10_140204_37_sti tch_rgb_big-2 |
|---|---|---|---|---|---|---|---|---|---|---|---|---|---|---|---|---|---|---|---|---|---|---|---|---|
| | | | | | | 10 | | | | | | | 11 | | | | | | | | | | | |
| 69 | 70 | 71 | 72 | 73 | 74 | 75 | 76 | 77 | 78 | 79 | 80 | 81 | 82 | 83 | 84 | 85 | 86 | 87 | 88 | 89 | 90 | 91 | 92 | 93 |
| 6 | 7 | 8 | 9 | 10 | 11 | 1 | 2 | 3 | 4 | 5 | 6 | 7 | 1 | 2 | 3 | 4 | 5 | 6 | 7 | 8 | 9 | 10 | 11 | 12 |
| L | R | R | R | R | R | L | L | L | L | R | R | R | L | L | R | R | R | R | R | R | R | R | R | R |
| | | ○ | | | | | | | | | | | | | | | | | | | | | | |
| | | | | | | | | | | | | | | | | | | | | | | ○ | | |
| | | | | | | | | | | | | ○ | | | | | | | | | | | | |
| | | | | | | | | | | | ○ | | | | | | | | | | | | | |
| | | | | | | | | | | | | | | | | | | | | | | | ○ | |
| | | | | | | | | | ○ | | | | | | | | | | | | | | | |
| | | | | | | | | | | | | | | | | | | | | | ○ | | | |
| | | | | | | | | | | | | | | | | | | | ○ | | | | | |
| | | | | | | | | | | | | | | | | | | | | | ○ | | | |
| | | | | | | | | | | | | | | | | | | | | | ○ | | | |
| | ○ | | | | | ○ | | | | | | | ○△ | | △ | | | | | | | | | |
| | ○ | | | | | ○ | | | | | | | ○△ | | △ | | | | | | | | | |
| | ○ | | | | | | | | | | | | ○ | | ○a | | | | | | | | | |
| | | | | | | ○a | | | | | | | | | | | | | | | | | | |
| | | | | | | | | | | | | | | ○ | | | | | | | | | | |
| | | | | | | | | | | | | | | | | | | | | ○ | | | | |
| ○ | | | ○ | | | | | | | | | | | | | | | | | | | | | |
| | | | | | | | | ○ | | | | | | | | | | | | | | | | |
| | | | | | | | | | | | | | | | | | | | | | | | | ○ |
| | | | | | | | | | | | | | | | | | ○oppo△ | | | | | | | |
| | | | | | | | | | | | | | | | | | ○ | | | | | | | |
| | | | | | | | | | | ○ | | | | | | | | | | | | | | |
| | | | | | | | | | | | | | | | | | △ | ○ | | | | | | |
| | | | | | | | ○ | | | | | | | | | ○ | | | | | | | | |
| | | | | ○ | | | | | | | | | | | | | | | | | | | | |
| | | | | | | | ○ | | | | | | | | | ○ | | | | | | | | |

| #11_131122_07m_stitch_m-1 | #11_131122_07m_stitch_m-1 | #11_131122_07m_stitch_m-1 | #11_131122_07m_stitch_m-1 | #11_131122_07m_stitch_m-1 | #11_131122_07m_stitch_m-1 | #11_131122_07m_stitch_m-1 | #12_131122_10m_stitch_m_rescale-1 | #12_131122_10m_stitch_m_rescale-1 | #12_131122_10m_stitch_m_rescale-1 | #12_131122_10m_stitch_m_rescale-1 | #12_131122_10m_stitch_m_rescale-1 | #13_131122_11m_stitch_m_rescale-1 | #13_131122_11m_stitch_m_rescale-1 | #13_131122_11m_stitch_m_rescale-1 | #13_131122_11m_stitch_m_rescale-1 | #13_131122_11m_stitch_m_rescale-1 | #13_131122_11m_stitch_m_rescale-1 | #13_131122_11m_stitch_m_rescale-1 | #13_131122_11m_stitch_m_rescale-1 | #13_131122_11m_stitch_m_rescale-1 | #13_131122_11m_stitch_m_rescale-1 | #13_131122_11m_stitch_m_rescale-1 | #13_131122_11m_stitch_m_rescale-1 | #13_131122_11m_stitch_m_rescale-1 | #13_131122_11m_stitch_m_rescale-1 | #13_131122_11m_stitch_m_rescale-1 | #13_131122_11m_stitch_m_rescale-1 | #14_131122_16m_stitch_m-1 | #14_131122_16m_stitch_m-1 | #14_131122_16m_stitch_m-1 | #14_131122_16m_stitch_m-1 | #14_131122_16m_stitch_m-1 | #14_131122_16m_stitch_m-1 | #15_130919_#5_3_1_rescale-1 | #15_130919_#5_3_1_rescale-1 | #15_130919_#5_3_1_rescale-1 | #15_130919_#5_3_1_rescale-1 | #16_130919_9_m_stitch_m-2 | #16_130919_9_m_stitch_m-2 | #16_130919_9_m_stitch_m-2 | #16_130919_9_m_stitch_m-2 | #16_130919_9_m_stitch_m-2 | #16_130919_9_m_stitch_m-2 | #16_130919_9_m_stitch_m-2 | #16_130919_9_m_stitch_m-2 | #16_130919_9_m_stitch_m-2 | #17_140901_02_w ms-1 | #17_140901_02_w ms-1 | #17_140901_02_w ms-1 |
|---|---|---|---|---|---|---|---|---|---|---|---|---|---|---|---|---|---|---|---|---|---|---|---|---|---|---|---|---|---|---|---|---|---|---|---|---|---|---|---|---|---|---|---|---|---|---|---|---|---|
| 12 | | | | | | | 13 | | | | | 14 | | | | | | | | | | | | | | | | 15 | | | | | | 16 | | | | 17 | | | | | | | | | 18 | | |
| 94 | 95 | 96 | 97 | 98 | 99 | 100 | 101 | 102 | 103 | 104 | 105 | 106 | 107 | 108 | 109 | 110 | 111 | 112 | 113 | 114 | 115 | 116 | 117 | 118 | 119 | 120 | 121 | 122 | 123 | 124 | 125 | 126 | 127 | 128 | 129 | 130 | 131 | 132 | 133 | 134 | 135 | 136 | 137 | 138 | 139 | 140 | 141 | 142 | 143 |
| 1 | 2 | 3 | 4 | 5 | 6 | 7 | 1 | 2 | 3 | 4 | 5 | 1 | 2 | 3 | 4 | 5 | 6 | 7 | 8 | 9 | 10 | 11 | 12 | 13 | 14 | 15 | 16 | 1 | 2 | 3 | 4 | 5 | 6 | 1 | 2 | 3 | 4 | 1 | 2 | 3 | 4 | 5 | 6 | 7 | 8 | 9 | 1 | 2 | 3 |
| L | L | L | R | R | R | R | L | R | R | R | R | L | L | L | L | L | L | L | R | R | R | R | R | R | R | R | R | L | L | L | L | R | R | R | R | R | R | L | L | L | R | R | R | R | R | R | L | L | L |
| | | | | | | | | | | | | | | | | | | | | | | | | | | | | | | | | | | | | | | | | | | | | | | | | | | | | | | | | | ○ | | ○ | | | | | | | | | | | | | | | | | | | | | | | | | | | | | | ○ |
| | | | | | | | | | | | | | | | | | | | | ○ | | | | | | | | | | | | | | | ○ | | | | | | | | | | ○ | | | | |
| | | | | | | | ○ | | | | | | | | | | | | | | | | | ○ | | | | | | | | | | | ○ | | | | | | | | | ○ | | | | | ○ |
| | | | | | | | | | | | | | ○ | | | | | | ○ | | | | | | | | | | | | ○ | | | | | | | | | | | | | | | | | | |
| | | | | | | | | | | | | | ○ | | | | | | | | | | | | | | ○ | | | | | | | | | | | | | | | | | | | | | | |
| | | | | | | | | ○ | | | | | | ○ | | | | | | | | | | | | ○ | | | | ○ | | | | | | | | ○ | | | | | | | | | | | |
| | | | | | | | | | | | | | | | | | | | | | | | | | ○ | | | | | | | | | | | | | | | | | | | | | | | | |
| | | | ○ | | | | | | | | | | | | | | | | | | ○ | | | | | | | | | | | | | | | | | | | | | | | | | | | | |
| | | | ○ | | | | | | | | | | | | | | | | | | | | | | | | | | | | | | | | | | | | | | | | | | | | | | |
| | | | | | | | | | | | | △ | | | | | | | | | | | | | | | | | | | | | | | | | | | | | | | | △○ | | | | | |
| | | | | | | | | | | | | △ | | | | | | | | | | | | | | | | | | | △p | | | | | | | | | | | | | △○ | | | | | |
| | | | | | | | | | | | | | | | | | | | | | | | | | | | | | | | △a | | | △ | | | | △ | | | | | | | | | | | |
| | | | | | | | | | | | | ○ | | | | | | | | | | | | | | | | | | | | | | △ | | | | △ | | | | | | | | | | | |
| | | | | | | | | | | | | | | | | | | | | | | | ○ | | | | | | | | | | ○ | | | | | | | | | | | | | | | | |
| | | | | | | | | | | ○ | | | | | | | | | | | | | | | | | | | | | | | | | | | | | | | | | | | | | | | |
| | | | | | | | | | | | | | | | | ○ | | | | | | | | | | | | | | | | | | | | | | | ○ | | | | | | | | | ○ | |
| ○ | ○ | | | | | | | | | | | | | | ○ | | | | | | | | | | | | | | | | | | | | | | | | | | | | ○ | | | | | | |
| | | | | | | ○ | | | | | | | | | | | | | | | | | | | | | | | | | | | | | | | | | | | | | | | | | | | |
| | | | | | | | | | | | | | | | | | | | | | | | | | | | | | | | | ○ | | | | | | | | | | | | | | | | | |
| | | | | | ○ | | | | | | | | | | | | | | | | | | | | | | | | | | | | | | | | | | | | | | | | | | | | |
| | | | | | | | | | | | | | | | | | ○ | | | | | | | | | | | | ○ | | | | | | | ○ | | | | | ○ | | | | | | | | |
| | | | | | | | | | | | | | | | | | | | | | | | | | | | | | | | | | | | | | | | | | | | | | △ | | | | |
| | | | | | | | △ | | | | | | | | | | | | | | | | | | | | | | | | | | | | | | | | | | | | | | | | | | |
| | | | | ○ | | | | | | | | | | | | | | | | | | | | | | | | | | | | | | | | | | | | | | | | | | | | | |
| | | | | | | | | | ○ | | | | | | | | | | | | | | | | | | | | | | | | | | | | | | | ○ | | | | | | | | | |
| | | | | | | | | | | | | | | | | | | | | | | | | | | | | | | | | | | | | | | | | | | | | | ○ | ○ | | |
| | | | | | | | | | | | | | | | | | | | | | | | | | | | | | | | | | | | | | | | | | | | | | | ○ | | |
| | | ○ | | | | | | | | | | | | | | | | | | | | | | | | | | | | | | | | | | | | | | | | | | | | | | | |
| | | | | | | | | | | ○ | | | | | | | | | | | | | | | | | | ○ | | | | | | ○ | | | | | | | | | | | | | | | |
| | | | | | | | | | | | | | | | | | | | | | | | | | | | | ○ | | | | | | ○ | | | | | | | | | | | | | | | |

| #17_140901_02_w ms-1 | #17_140901_02_w ms-1 | #17_140901_02_w ms-1 | #17_140901_02_w ms-1 | #19_140901_04_w ms-1 | #19_140901_04_w ms-1 | #19_140901_04_w ms-1 | #19_140901_04_w ms-1 | #19_140901_04_w ms-1 | #19_140901_04_w ms-1 | #19_140901_04_w ms-1 | #19_140901_04_w ms-1 | #20_140901_05_w ms-1 | #20_140901_05_w ms-1 | #20_140901_05_w ms-1 | #20_140901_05_w ms-1 | #20_140901_05_w ms-1 | #20_140901_05_w ms-1 | #20_140901_05_w ms-1 | #20_140901_05_w ms-1 | #20_140901_05_w ms-1 | #20_140901_05_w ms-1 | #20_140901_05_w ms-1 | #20_140901_05_w ms-1 | #20_140901_05_w ms-1 | #21_141212_08-1 | #21_141212_08-1 | #21_141212_08-1 | #21_141212_08-1 | #21_141212_08-1 | #22_141212_09-1 | #22_141212_09-1 | #22_141212_09-1 | #22_141212_09-1 | #22_141212_09-1 | #22_141212_09-1 | #22_141212_09-1 | #22_141212_09-1 | #23_141212_10-1 | #23_141212_10-1 | #23_141212_10-1 | #23_141212_10-1 | #23_141212_10-1 | #23_141212_10-1 | #23_141212_10-1 | #23_141212_10-1 | #23_141212_10-1 | #23_141212_10-1 | #23_141212_10-1 | #23_141212_10-1 |
|---|---|---|---|---|---|---|---|---|---|---|---|---|---|---|---|---|---|---|---|---|---|---|---|---|---|---|---|---|---|---|---|---|---|---|---|---|---|---|---|---|---|---|---|---|---|---|---|---|---|
| | | | | 19 | | | | | | | | 20 | | | | | | | | | | | | | 21 | | | | | 22 | | | | | | | | 23 | | | | | | | | | | | |
| 144 | 145 | 146 | 147 | 148 | 149 | 150 | 151 | 152 | 153 | 154 | 155 | 156 | 157 | 158 | 159 | 160 | 161 | 162 | 163 | 164 | 165 | 166 | 167 | 168 | 169 | 170 | 171 | 172 | 173 | 174 | 175 | 176 | 177 | 178 | 179 | 180 | 181 | 182 | 183 | 184 | 185 | 186 | 187 | 188 | 189 | 190 | 191 | 192 | 193 |
| 4 | 5 | 6 | 7 | 1 | 2 | 3 | 4 | 5 | 6 | 7 | 8 | 1 | 2 | 3 | 4 | 5 | 6 | 7 | 8 | 9 | 10 | 11 | 12 | 13 | 1 | 2 | 3 | 4 | 5 | 1 | 2 | 3 | 4 | 5 | 6 | 7 | 8 | 1 | 2 | 3 | 4 | 5 | 6 | 7 | 8 | 9 | 10 | 11 | 12 |
| R | R | R | R | L | L | L | L | L | R | R | R | L | L | L | L | L | L | R | R | R | R | R | R | R | L | R | R | R | R | L | L | L | L | R | R | R | R | L | L | L | L | R | R | R | R | R | R | R | R |
| | | | | | | | | | | | | | | | | | | | | | | | | | | | | | | | | | | ○ | | | | | | | | | | | | | | | |
| | | | ○ | | | | | | | | | | | | | | | | | | | | | | | | | | | | | | ○ | | | | | | | | | | ○ | | | | | | |
| | | | | | | | | | | | | | | | | | ○ | | | | | | | | | | | | | | | | | | | ○ | | | | | | | ○ | | | | | | |
| | | | | ○ | | | | | | | | | | | | | | | | | | | | | | | | | | | | | | | | | | | | | | | | ○ | | | | | |
| | | | | | | | | | | | | ○ | | | | | | | | | | | | | | | | | | | | | | | ○ | | | | | | | | | | ○ | | | | |
| | | | | | | | | | | | | | | | | | | | | | | | | | | | | | | | | | | | | | | | | | | | | | | ○ | | | |
| | | | | | | | | | | | | | | | | | | | | | | | | | | | | ○ | | | | | | | | | ○ | | | | | | | | | | | | |
| | | | | | | | | | | | | | | | | | | | ○ | | | | | | | | | | | | | | | | | | | | | | | | | | | | | | |
| | | | | | ○ | | | | | | | | | | | | | | | | | | | | | | | | | | | | | | | | | | | | | | | | | | | | |
| | | | | | | | | | | | | | | | | | | | ○ | | | | | | | | | | ○ | | | | | | | | | | | | | | | | | | | | |
| | | | | | | ○ | | | | | ○ | | | | | | | | | | | | | | | | | | | | | | | | | | | ○ | | | | | | | | | | | |
| | | | | | | | | | | | | | | | | | | | | | | | | | | | | | | | | | | | | △ | | | | | | | | | | | | | |
| | | | | △○ | | | | | | △ | | △p | | | | | △ | △ | | | | | | | | | | | | | | | | | | | | | | | | | | | | | | | |
| | | | | | | △○ | | | | △ | | △p | | | | | △ | △ | | | | | | | | | | | | | | | | | | | | | | | | | | | | | | | |
| | | | | | | | | | | | | | | | ○ | | | | | | | | | | | | | | | | | | | | | | | | | | | | | | | | | | |
| | | | | | | | | ○ | | ○ | | | | | ○ | | | ○ | | | | | | | | | | | | ○ | | | | | | | | ○ | | | | | | | | | | | |
| | | | | | | | | | | | | | ○ | | | | | | | | | | | ○ | | | | | | | | | | | | | | | | | | | | | | | | | |
| | | | | | | | | | | | | | | | | | | | | | | | | | | | | | | | | | | | | | | | ○ | | | | | | | | | | |
| | | ○ | | | | | | | | | | | | | | ○ | | | | | | | | | | | | | | | | | | | | | | | | | | | | | | | | | |
| | | | | | | | | | | | | | | | | | | | | | | | | | | | | | | | | | | | | | | | | | | | | | | | | | ○ |
| | | | | | | | | | | | | | | | | | | | | | | | | | | | | | | | | | | | | | | ○ | | | | | | | | | | | |
| | | | | | | | | | | | | | | | | | | | | | | | | | | | | | | | ○ | | | | | | | | | | | | | | | ○ | | |
| | | | | | | | | | | | | | | | | | | | | ○ | | | | | | | | | | | | | | | | | | | | | | ○ | | | | | | | |
| | | | | | | | | | | | | | | | | | | | | | | | ○△oppo | | | | | | | | | | | | | | | | | | | | | | | | | | |
| | | △ | | | | | | | | | | | △ | | | | | | | | | | | | | | | | | | | | | | | | | | | | | △oppo | | | | | | | |
| | | | | | | | | | | | | | | | | | | | | | | | | | ○ | ○ | | | | | | | | | | | | | | | | | | | | | | | |
| | ○ | | | | | | | | ○ | | | | | | | | | | | △ | | ○ | | | | | | | | | | ○ | | | | | | | | | ○ | | | | | | | | |
| | | | | | | | | | ○ | | | | | | | | | | | | | | | | | | | | | | | | | | | | | | | | ○ | | | | | | | | |
| ○ | | | | | | | | | | | | | | | | | | | | | | | | | | | | | | | | | | | | | | | | | | | | | | | | | |
| | | | | | | | | | ○ | | | | | ○ | | | | | | | ○ | | | | | | ○ | | | | | ○ | | | | | | | | | ○ | | | | | | | ○ | |
| | | | | | | | | | | | | | | | | | | | | | | | | | | | | | | | | | | | | | | | | | ○ | | | | | | | | |
| | | | | | | | | | | | | | | | | | | | | | | | | | | | ○ | | | | | | | | | | | | | | | | | | | | | | |
| | | | | | | | ○ | | ○ | | | | | | | | | | | | ○ | | | | | | ○ | | | | | | | | | | | | | | | | | | | | | ○ | |

| #24_141213_04-1 | #24_141213_04-1 | #24_141213_04-1 | #24_141213_04-1 | #24_141213_04-1 | #24_141213_04-1 | #25_141213_05-1 | #25_141213_05-1 | #25_141213_05-1 | #25_141213_05-1 | #25_141213_05-1 | #25_141213_05-1 | #25_141213_05-1 | #25_141213_05-1 | #26_141213_06-1 | #26_141213_06-1 | #26_141213_06-1 | #26_141213_06-1 | #26_141213_06-1 | #26_141213_06-1 | #26_141213_06-1 | #26_141213_06-1 | #26_141213_06-1 | #26_141213_06-1 | #26_141213_06-1 | #26_141213_06-1 | #26_141213_06-1 | #26_141213_06-1 | #26_141213_06-1 | #26_141213_06-1 | #26_141213_06-1 | #27_141213_09-1 | #27_141213_09-1 | #27_141213_09-1 | #27_141213_09-1 | #27_141213_09-1 | #27_141213_09-1 | #27_141213_09-1 | #27_141213_09-1 | #27_141213_09-1 | #27_141213_09-1 | #27_141213_09-1 | #28_141213_10-1 | #28_141213_10-1 | #28_141213_10-1 | #29_141213_11-1 | #29_141213_11-1 | #29_141213_11-1 | #29_141213_11-1 | #29_141213_11-1 |
|---|---|---|---|---|---|---|---|---|---|---|---|---|---|---|---|---|---|---|---|---|---|---|---|---|---|---|---|---|---|---|---|---|---|---|---|---|---|---|---|---|---|---|---|---|---|---|---|---|---|
| 24 | | | | | | 25 | | | | | | | | 26 | | | | | | | | | | | | | | | | | 27 | | | | | | | | | | | 28 | | | 29 | | | | |
| 194 | 195 | 196 | 197 | 198 | 199 | 200 | 201 | 202 | 203 | 204 | 205 | 206 | 207 | 208 | 209 | 210 | 211 | 212 | 213 | 214 | 215 | 216 | 217 | 218 | 219 | 220 | 221 | 222 | 223 | 224 | 225 | 226 | 227 | 228 | 229 | 230 | 231 | 232 | 233 | 234 | 235 | 236 | 237 | 238 | 239 | 240 | 241 | 242 | 243 |
| 1 | 2 | 3 | 4 | 5 | 6 | 1 | 2 | 3 | 4 | 5 | 6 | 7 | 8 | 1 | 2 | 3 | 4 | 5 | 6 | 7 | 8 | 9 | 10 | 11 | 12 | 13 | 14 | 15 | 16 | 17 | 1 | 2 | 3 | 4 | 5 | 6 | 7 | 8 | 9 | 10 | 11 | 1 | 2 | 3 | 1 | 2 | 3 | 4 | 5 |
| L | L | R | R | R | R | L | L | L | R | R | R | R | R | L | L | L | L | L | L | L | L | R | R | R | R | R | R | R | R | R | L | L | L | L | L | L | R | R | R | R | R | L | L | R | L | L | L | L | L |
| | | | | | ○ | | | ○ | | | | | | | | | | ○ | | | | | | | | | | | | | | | | | | ○ | | | | | | | | | | | | | |
| | | | | | | | | | | | | | | | | | | | | | | | | | | | | | | | | | | | | | | | | | | ○ | | | | | | | |
| | | | | | | | | | | | | | | | | | | | | | | | | | | | | | | | | | | | | | | | | | ○ | | | | ○ | | | | |
| | | | | | | | | | | | | | | | | | | | | | | | | | | | ○ | | | | | | | | | | | | | | | | | | | | | | |
| | | | ○ | | | | | | | | | | | | | | ○ | | | | | | | | | | | | | | | | | | | | | | | | | | | | | | | | |
| | | | | | | | | | | | ○ | | | | | | | | | ○ | | | | | ○ | | | | | | ○ | | | | | | | | ○ | | | | | | | | ○ | | |
| | | | | | | | | | | | | | | | | | | | | | | | | | | | | | | | | | | ○ | | | | | | | | | | | | | | | |
| | | | | | | | | | ○ | | | | | | | | | | | | | | | | | | | | | | | | | | | | | | | | | | | | | | | | |
| | | | | | | | | | | ○ | | | | | | | | | | | | | | | | | | | | | | | | | | | | | | | | | | | | ○ | | | |
| | | | | | | | | | ○ | | | | | | | | | | | | | | | | | | | | | | | | | | | | | | | | | | | | | | | | |
| | | | | | | | | | | | | | | | | | | | | | | | | | | | | | | | | | | | | | | | | | | | | | | | | | ○ |
| | | | | ○ | | | | | | | | | | | | | | | | | | | | △ | | | △ | | | | | | | | | | | | | | | | | | | | | | |
| | | | | | | | | | | | | | | | | | | | | | | | | △ | | | | | | | ○ | | | | | | | | | | | | | | | | | | |
| | | | | △ | | | | | | | △ | | | | | | | | | | | | | | | | | | | | △○? | | | | | | | ○ | | | | | | | | | | | |
| | | | | △ | | | ○ | | | | | | | | | | | | | | | | | | | | | | | | △○? | | | | | | | ○ | | | | | | | | | | | |
| | | | △ | | | | | | | | | | | | | | ○ | | | | | | | ○ | | | | | | | | | | | | | | | | | | | | | | | | | |
| | | | | △ | | | | | | | | | | | | | | | | | | | | | | | | | | | | | | | | | | ○ | | | | | | | | | | | |
| | ○ | | | | | | | | | | | | | | | | | | | | | | | | | | | | | | | | | | | | | | | | | | | | | | | | |
| ○ | | | | | | | | | | | | | ○ | | | | | | | | ○ | | | | | | | | | | | | | | | | | | | | | | | ○ | | | | | |
| | | | | | | | | | | | | | | | | | | | | | | | | | | | | | | | | | | | ○ | | | | | | | | | | | | | | |
| | | | | | | | | | | | | | | | ○ | | | | | | | | | | | ○ | | | | | | | | | ○ | | | | | | | | | | | | | | |
| | | | | | | | | | | | | | | | | | | | | | | | ○ | | | | | | | | | | | | | | | | | | | | | | | | | | |
| | | ○ | | | | m○△ | | | | | | | | | | | | | ○ | | | | | | | | | | | | | ○ | | | | | | | | ○ | | | | | | | | | |
| | | | | | | | | | | | | | | | | | | | | | | | | | | | | | | ○ | | | | | | | | | | | | | ○ | | | | | | |
| | | | | | | | | | | | | | | | | ○ | | | | | | | | | | | | | | | | | | | | | | | | | | | | | | | | | |
| | | | | | | ○ | | | | | | ○ | | ○ | | ○ | | | | | | | | | | | | | ○ | | | | | | | ○ | | | | ○ | | | | | | | | ○ | |
| | | | | | | | | | | | | | | | | ○ | | | | | | ○ | | | | | | | | | | | | | | | | | | | | | | | | | | | |
| | | | | | | | | | | | | | | | | | | | | | | | | | | | | | ○ | | | | | | | ○ | | | | ○ | | | | | | | | | |
| | | | | | | ○ | | | | | | | | | | | | | | | | | | | | | | | ○ | | | | | | | ○ | | | | | | | | | | | | | |
| | | | | | | | | | | | | | | | | | | | | | | ○ | | | | | | | | | | | | | | | | | | | | | | | | | | | |

| #29_141213_11-1 | #29_141213_11-1 | #29_141213_11-1 | #30_141213_12-1 | #30_141213_12-1 | #30_141213_12-1 | #30_141213_12-1 | #30_141213_12-1 | #31_141213_13-1 | #31_141213_13-1 | #31_141213_13-1 | #31_141213_13-1 | #32_141213_14-1 | #32_141213_14-1 | #32_141213_14-1 | #32_141213_14-1 | #32_141213_14-1 | #32_141213_14-1 | #33_141213_15-1 | #33_141213_15-1 | #33_141213_15-1 | #33_141213_15-1 | #33_141213_15-1 | #33_141213_15-1 | #33_141213_15-1 | #33_141213_15-1 | #33_141213_15-1 | #33_141213_15-1 | #34_141213_17-1 | #34_141213_17-1 | #34_141213_17-1 | #34_141213_17-1 | #34_141213_17-1 | #34_141213_17-1 | #34_141213_17-1 | #35_141213_18-1 | #35_141213_18-1 | #35_141213_18-1 | #35_141213_18-1 | #35_141213_18-1 | #35_141213_18-1 | #36_141213_20-1 | #36_141213_20-1 | #36_141213_20-1 | #36_141213_20-1 | #36_141213_20-1 | #37_141213_21-1 | #37_141213_21-1 | #37_141213_21-1 | #37_141213_21-1 |
|---|---|---|---|---|---|---|---|---|---|---|---|---|---|---|---|---|---|---|---|---|---|---|---|---|---|---|---|---|---|---|---|---|---|---|---|---|---|---|---|---|---|---|---|---|---|---|---|---|---|
| | | | 30 | | | | | 31 | | | | 32 | | | | | | 33 | | | | | | | | | | 34 | | | | | | | 35 | | | | | | 36 | | | | | 37 | | | |
| 244 | 245 | 246 | 247 | 248 | 249 | 250 | 251 | 252 | 253 | 254 | 255 | 256 | 257 | 258 | 259 | 260 | 261 | 262 | 263 | 264 | 265 | 266 | 267 | 268 | 269 | 270 | 271 | 272 | 273 | 274 | 275 | 276 | 277 | 278 | 279 | 280 | 281 | 282 | 283 | 284 | 285 | 286 | 287 | 288 | 289 | 290 | 291 | 292 | 293 |
| 6 | 7 | 8 | 1 | 2 | 3 | 4 | 5 | 1 | 2 | 3 | 4 | 1 | 2 | 3 | 4 | 5 | 6 | 1 | 2 | 3 | 4 | 5 | 6 | 7 | 8 | 9 | 10 | 1 | 2 | 3 | 4 | 5 | 6 | 7 | 1 | 2 | 3 | 4 | 5 | 6 | 1 | 2 | 3 | 4 | 5 | 1 | 2 | 3 | 4 |
| R | R | R | L | L | R | R | R | L | R | R | R | L | L | L | L | R | R | L | L | R | R | R | R | R | R | R | R | L | L | L | R | R | R | R | L | L | L | R | R | R | L | L | R | R | R | L | L | L | L |
| | | | | | | | | | | | ○ | | | | | | | | | | | | | | | | | | | | | | | | | | | | | | | | | | ○ | | | | |
| | | ○ | | | | △ | | | | | | | | | | | | | | | | | | | | | | | | ○ | | | | | | | | | | | | | | | | | | | |
| | | | | | | | | | | | | | | | | | | | ○ | | | | | | | | | | | | | | | | | | | | | | | | | | | | | | |
| | | | ○Dd2a | | | | | | | | | | | | | | | | | | | | | | | | | | | | | | | | | | | ○ | | | | | | | | | | | |
| | | | | | | | ○ | | | | | | | ○ | | | | | | | | | | | | | | | ○ | | | | | | | | ○ | | ○ | | | | | | | | | | |
| | | | | | | | ○ | | | | | | ○ | | | | | | | | | ○ | | | | | | | | | | | | | | | | | | | | | | | | | | | |
| | | | | | | | | | | | | | | | | | | | | | | | | | | | | | | | △ | | | | | ○ | | | | | | | | | | | | ○ | |
| | | | | | | | | | | | | | | | | | | | | | | | ○ | | | | | | | | | | | | | | | | | | | | | | | | | | |
| | | | | | | | | | | | | | | | | | | | | | | | | ○ | | | | | | | △ | | | | | ○△ | | | | | | | | | | | | ○ | |
| | | | | | | | | | | | | | | | | | | | | | | | | | ○ | | | | | | | | | | | ○△ | | | | | | | | ○ | | | | | |
| | | | | | | ○ | | | | | | | | | | | | | | | | | | | | | | | | | △ | | | | | | | | | | | | | | | | | ○ | |
| | ○ | | | | | | △ | | | | | | | | | | | ○ | | | ○ | | | | | | | | | | | | | | | | | ○ | | | | | | | | | | | |
| | | | | | | | △ | | | | | | | | | | | | | | | | | | | | | | | | | | | | | | | | | | | | | | | | | | |
| | △ | | △ | | | | ○ | | | | | | | △ | | | | | | | △ | | | | | | | | △ | | | | | | | | | △ | | | ○ | | △ | | | | | | |
| | | | △ | | | | | | | | | | | △ | | | | △ | | | △ | | | | | | | | | | △ | | | | | | | △ | | | ○ | | △ | | | | | | |
| | | | ? | | | | | | | | | | | ○ | | | △ | | | | | | | | | | | | | | | | | | | | | | | | △ | | ○ | | | | | | |
| | | | ? | | | | | | | | | | | | | | | △ | | | ○ | | | | | | | | | | △ | | | | | | | ○ | | | △ | | ○ | | | | | | |
| | | | | | | | | | | | | | | | | | | | | | | | | | | | | | | | | ○ | | | | | | | | | | | | | | | | | |
| | | | | | | | | | | | | | | | | | | | | | | | | | | | | | | | | | | | | | | | | | | ○ | | | | | | | |
| | | | | | | | | | | | | | | | | | | | | | | | | | | | | | | | | | | | | | | | | ○ | | | | | | | | | |
| | | | | | | | | | | | | | | | | | | | | | | | | | | | | | | | | | | ○ | | | | | | | | | | | | | | | |
| | | | | | | | | | | | | | | | | | | | | | | | | | | ○ | | | | | | | | | | | | | | | | | | | | | | | |
| | | | | | | | | | | | | | | ○ | ○ | | | | | | | | | | | | | | | ○ | | | | | | | | | | | | | | | | | | |
| | | | | ○ | ○ | | | | | | | | | | | | | | | ○ | | | | | | | | | | | | | | | | | | | | | | | | | | | ○△oppo | | |
| | | | | ○op | ○opp | | | | | | | | | | | | | | | ○ | | | | | | | | | | | | | | | | | | | | | | | | | | | | | |
| | | | | | | | | | | | | | | | | | | | | | | | | | | | | | | | | | | | | △ | | | | | | | | | | | | | |
| | | | | | | | | | | | | | | | | | | | | | | | | | | | | | | | | | | | | △ | | | | | | | | | | | | | |
| | | | | △ | △ | | | | | | | ○ | | | | | | | | △ | | | | | | | | | | | | | | | | △ | | | | | | ○ | | | | | | △ | |
| | | | | | | | | | ○ | △ | | | | | | | | | | | | | | | | | | ○ | | | | | | | | | | | | | | ○ | | | | | | | |
| ○ | | | | | | | | | ○ | | | | | △ | | | ○ | | | | | | | | | | | ○ | | | ○ | | | | ○ | | | | | | | ○ | | | | ○ | | | |
| | | | | | | | | | ○ | | | | | | | | | | | | | | | | | | | | | | ○ | | | | | | | | | | | | | | | | | | |
| ○ | | | | | | | | ○ | ○ | | | | | △ | | | ○ | | | | | | | | | | | ○ | | | ○ | | | | | | | | | | | ○ | | | | ○ | | | |
| ○ | | | | | | | | ○ | ○ | | | | | △ | | | | | | | | | | | | | | ○ | | | ○ | | | | ○ | | | | | | | ○ | | | | ○ | | | |
| | | | | | | | | | ○ | | | | | | | | | | | | | | | | | | | | | | ○ | | | | | | | | | | | | | | | | | | |
| | | | | | | | | | △ | | | | | | | | | | | | | | | | | | | | | | | | | | | | | | | | | | | | | | | | |

| #39_150220_03-1 | #39_150220_03-1 | #39_150220_03-1 | #39_150220_03-1 | #39_150220_03-1 | #39_150220_03-1 | #40_150220_06-1 | #40_150220_06-1 | #40_150220_06-1 | #40_150220_06-1 | #40_150220_06-1 | #40_150220_06-1 | #40_150220_06-1 | #40_150220_06-1 | #40_150220_06-1 | #41_150220_07-1 | #41_150220_07-1 | #41_150220_07-1 | #41_150220_07-1 | #41_150220_07-1 | #43_150220_12-1 | #43_150220_12-1 | #43_150220_12-1 | #43_150220_12-1 | #43_150220_12-1 | #43_150220_12-1 | #44_150220_13-1 | #44_150220_13-1 | #44_150220_13-1 | #44_150220_13-1 | #44_150220_13-1 | #44_150220_13-1 | #44_150220_13-1 | #45_150220_14-1 | #45_150220_14-1 | #45_150220_14-1 | #45_150220_14-1 | #45_150220_14-1 | #45_150220_14-1 | #45_150220_14-1 | #45_150220_14-1 | #46_150220_15-1 | #46_150220_15-1 | #46_150220_15-1 | #46_150220_15-1 | #46_150220_15-1 | #46_150220_15-1 | #47_150220_16-1 | #47_150220_16-1 | #47_150220_16-1 |
|---|---|---|---|---|---|---|---|---|---|---|---|---|---|---|---|---|---|---|---|---|---|---|---|---|---|---|---|---|---|---|---|---|---|---|---|---|---|---|---|---|---|---|---|---|---|---|---|---|---|
| 38 | | | | | | 39 | | | | | | | | | 40 | | | | | 41 | | | | | | 42 | | | | | | | 43 | | | | | | | | 44 | | | | | | 45 | | |
| 294 | 295 | 296 | 297 | 298 | 299 | 300 | 301 | 302 | 303 | 304 | 305 | 306 | 307 | 308 | 309 | 310 | 311 | 312 | 313 | 314 | 315 | 316 | 317 | 318 | 319 | 320 | 321 | 322 | 323 | 324 | 325 | 326 | 327 | 328 | 329 | 330 | 331 | 332 | 333 | 334 | 335 | 336 | 337 | 338 | 339 | 340 | 341 | 342 | 343 |
| 1 | 2 | 3 | 4 | 5 | 6 | 1 | 2 | 3 | 4 | 5 | 6 | 7 | 8 | 9 | 1 | 2 | 3 | 4 | 5 | 1 | 2 | 3 | 4 | 5 | 6 | 1 | 2 | 3 | 4 | 5 | 6 | 7 | 1 | 2 | 3 | 4 | 5 | 6 | 7 | 8 | 1 | 2 | 3 | 4 | 5 | 6 | 1 | 2 | 3 |
| L | L | L | R | R | R | L | L | L | L | L | R | R | R | R | L | L | R | R | R | L | L | L | R | R | R | L | L | L | R | R | R | R | L | L | L | R | R | R | R | R | L | L | R | R | R | R | L | R | R |
| | | | | | ○ | | | | | | | | | | | | | | | | | ○ | | | | | | | | | | | | | | | | | | | | | | | | | | | |
| | | | | | | | | | | | | | | | | | | | | | | | | | | | | | | | | | | | | | | ○ | | | | | | | | | | | |
| | | | | ○ | | | | | | | | | | | | | | | | | | | | | | | | | | | | | | | | | | | ○ | | | | | | | | | | |
| | | | | | | | | | | | | | | ○ | | | | | | | | | | | | | | | ○ | | | | | | | | | | | | | | | | | | | | |
| | | | | | | | | | | | ○ | | | | | | | | | | | | | | | | | | | | | | | | | | | | | | | | | | | | | | |
| | | | | | | | | | | | | | | | | | | | | | | | | | ○ | | | | | | | | | | | | | | | | | | | | | △ | | | |
| | | | | | | | | | | | | | | | | | | | | | | | | | | | | | | | | | | | | | | | | | | | | | | ○ | | | |
| | | | | | | | | | | | | | | | | | | | | | | | | | | | | | | | | | | | | | ○ | | | | | | | | | ○ | | | |
| | | | | | | | ○ | | | | | | | | | | | | | | | | | | | △ | | | | | | | | | | | | | | | | | | | | | | | |
| | | | | | | | | | | | | | | | | | | | | | | | | ○ | | | | | | | | | | | | | | | | | | | | | | ○ | | | |
| | | | | | | | ○ | | | | | | | | | | | | | | | | | | | △ | | | | | | | | | | | | | | | | | | | | | | | |
| △ | | | ○ | | | | | | | | | | | ○ | | | | | | | | | | | | | | | △ | | | | | | | ○ | | | | | | | | | | | | | |
| △ | | | ○ | | | | | | | | | | | ○ | | | | | | | | | | | | | | | △ | ○ | | | | | | ○ | | | | | | | | | | | | | |
| | | | | | | | | | | | | | | | | | | | | | | | | | △only edge | | | | △ | | | | | | | | | | | | | | | | | | | | |
| ○ | | | | | | | | | | | | | | | | | | | | | | | | | | | | | | | | | | | | ○ | | | | | ○ | | | | ○ | | | | |
| | | ○ | | | | | | | | | | | | | | | | | | | | | | | | | | ○ | | | ○ | | | | | | | | | | | | | | | △ | | | |
| | | | | | | | | | | | | | | | | | | | | | ○ | | | | | | | | | | | | | | | | | | | | | | | | | △ | | | |
| | | | | | | | | | | | | | | | | | ○ | | | | | | | | | | | | | | | | | | | | | | | | | | | | | △ | | | |
| | ○ | | | | | | | | | | | | | | | | | | | | | | | | | | | | | | | | | | | | | | | | | | | | | | | | |
| | | | | | | | | | | | | | | | | | | ○ | | | | | | | | | | | | | | | | | | | | | | | | | | | | | | | |
| | | | | | | | | | | | | | | | | | | | | | | | | | | | | | | | | | | | | | | | | ○ | | ○ | | | | | | | |
| | | | | | | | | | ○ | | | | | | | | | | | | | | | | | | | | | | | | | | | | | | | | | | | | | | | |
| | | | | | | | | | | | | | | | | | | | | | | | | | | | | | | | | | | | | | | | | | | | △ | | | | | | |
| | | | | | | ○oppo | | | | | | ○oppo? | | | | | | | | | | | | | | | | | | | | | | | | | | | | | | | | ○oppo | | | | | |
| | | | | | | ○ | | | | | | | | | | | | | | | | | | | | | | | | | | | | | | | | | | | | | | | | | | | |
| | | | | | | | △ | | | | | | | | | | | | | | | | | | | | | | | | | | | | | | | | | | | | △ | | | | | | |
| | | | | | | | △ | | | | | | | | | | | | | | | | | | | | | | | | | | | | | | | | | | | | △ | | | | | | |
| | | | | | | | | | | | | | | | | | | | | | | | | | | ○ | | | | | | | | ○ | | | | | | | | | | | | | | | |
| | | | | | | △ | | | | | | △ | | | | | | | | | | | | | | | | | | | | ○ | ○ | | | | | | | | | | ○ | △ | | | ○v | ○v | |
| | | | | | | | | | | | | | | | | | | | | | | | | | | | | | | | | | | | | | | | | | | | ○ | | | | | | |
| | | | | | | | | | | | | | ○ | | ○ | | | | | ○ | | | | | | | ○ | | | | | | | | | | | | | | | | ○ | | | | | | |
| | | | | | | | | | | | | | ○ | | | | | | | ○ | | | ○ | | | | ○ | | | | | | | | | | | | | | | | ○ | | | | | | |
| | | | | | | | | | | | | | | | | | | | | | | | | | | | | | | | | | | | | | | | | | | | ○ | | | | | | |
| | | | | | | | | | | | | | | | | | | | | | | | | | | | | | | | | | | | | | | | | | | | ○ | | | | | | |
| | | | | | | | | | | | | | | | | | | | | | | | | | | | | | | △? | | | | | | | | | | | | | | | | | | | |

| #48_150220_17-1 | #48_150220_17-1 | #48_150220_17-1 | #48_150220_17-1 | #48_150220_17-1 | #48_150220_17-1 | #48_150220_17-1 | #48_150220_17-1 | #48_150220_17-1 | #49_150220_18-1 | #49_150220_18-1 | #49_150220_18-1 | #49_150220_18-1 | #49_150220_18-1 | #49_150220_18-1 | #49_150220_18-1 | #49_150220_18-1 | #49_150220_18-1 | #50_150220_19-1 | #50_150220_19-1 | #50_150220_19-1 | #50_150220_19-1 | #52_Composite_15 0404_05-1 | #52_Composite_15 0404_05-1 | #52_Composite_15 0404_05-1 | #52_Composite_15 0404_05-1 | #52_Composite_15 0404_05-1 | #53_Composite_15 0404_06-1 | #53_Composite_15 0404_06-1 | #53_Composite_15 0404_06-1 | #53_Composite_15 0404_06-1 | #53_Composite_15 0404_06-1 | #53_Composite_15 0404_06-1 | #55_Composite_15 0404_08-1 | #55_Composite_15 0404_08-1 | #55_Composite_15 0404_08-1 | #55_Composite_15 0404_08-1 | #55_Composite_15 0404_08-1 | #55_Composite_15 0404_08-1 | #56_Composite_15 0404_09-1 | #57_Composite_15 0404_10-1 | #57_Composite_15 0404_10-1 | #57_Composite_15 0404_10-1 | #57_Composite_15 0404_10-1 | #58_Composite_15 0404_11-1 | #58_Composite_15 0404_11-1 | #58_Composite_15 0404_11-1 | #58_Composite_15 0404_11-1 | #58_Composite_15 0404_11-1 | #58_Composite_15 0404_11-1 |
|---|---|---|---|---|---|---|---|---|---|---|---|---|---|---|---|---|---|---|---|---|---|---|---|---|---|---|---|---|---|---|---|---|---|---|---|---|---|---|---|---|---|---|---|---|---|---|---|---|---|
| 46 | | | | | | | | | 47 | | | | | | | | | 48 | | | | 49 | | | | | 50 | | | | | | 51 | | | | | | 52 | 53 | | | | 54 | | | | | |
| 344 | 345 | 346 | 347 | 348 | 349 | 350 | 351 | 352 | 353 | 354 | 355 | 356 | 357 | 358 | 359 | 360 | 361 | 362 | 363 | 364 | 365 | 366 | 367 | 368 | 369 | 370 | 371 | 372 | 373 | 374 | 375 | 376 | 377 | 378 | 379 | 380 | 381 | 382 | 383 | 384 | 385 | 386 | 387 | 388 | 389 | 390 | 391 | 392 | 393 |
| 1 | 2 | 3 | 4 | 5 | 6 | 7 | 8 | 9 | 1 | 2 | 3 | 4 | 5 | 6 | 7 | 8 | 9 | 1 | 2 | 3 | 4 | 1 | 2 | 3 | 4 | 5 | 1 | 2 | 3 | 4 | 5 | 6 | 1 | 2 | 3 | 4 | 5 | 6 | 1 | 1 | 2 | 3 | 4 | 1 | 2 | 3 | 4 | 5 | 6 |
| L | L | L | R | R | R | R | R | R | L | L | L | L | L | L | L | R | R | L | L | R | R | L | L | R | R | R | L | L | L | L | R | R | L | L | L | R | R | R | L | L | L | R | R | L | L | L | L | L | L |
| | | | | | | | | ○ | | | | | | | | | | | | | | | | | | | | | | | | | ○ | | | | | ○ | | ○ | | | | | | | | | |
| | | | | | | | | ○ | | | | | | | | | | | | | ○ | | | | | | | | | | | | | | | | | | | | | | | | | | | | |
| | ○ | | | | | | | | | | | | | | | | | | | | | | | | ○ | | | | | | | | | | | | | | | | | | | | | | | | |
| | | | | | | | | | | | | | | | | | | | | | | | | | | | | | | | | | | | | | | | | | | | ○ | | | | | | |
| | | | | | | ○ | | | | | | | | ○ | | | | | | | | | | | | | | | | | | | | | | | | | | | | | | | | | | | |
| | | | | | | ○ | | | | | | | | ○ | | | | | | | | | | | | | | | | | | | | | | | | | | | | | | | | | | | |
| | | | | | | | | | | | | | | | | | | | | | | | ○ | | | | | | | | | | | | | | | | | | | | | | | | | | |
| | | | | | | ○ | | | | | | | | ○ | | | | | | | | | | | | | | | | | | | | | | | | | | | | | | | | | | | |
| | | | | | | | | | | | | | | | | | | | | | | ○ | | | | | | | | | | | | | | | | | | | | | | | | | | | |
| | | | | | | | △ | | | | | | △ | | | | | | | | | | | | | | | | | | | | | | | | ○ | | | | ○ | | | | | | | | |
| | | | | | | | △ | | | | | | △ | | | | | | | | | | | | | | | | | | | | | | | | ○ | | | | ○ | | | | | | | | |
| | | | | | | | | | | | | | | | | | | | | | | | | | | | | | | | | | | | | | | | | | △ | | | ○ | | | | | |
| | | | | | | | | | | | | | | | | | | | | | | | | | | | | | | | | | | | | | | | | | △ | | | ○ | | | | | |
| ○ | | | | | | | | | | | | | ○ | | | | | | | | | | | | | | | | | | | | | | | | | | | | | | | △ | | | | | |
| | | | | | | | | | | | | | | | | | | | | | | | | | | | | | | | | | | | | | | | | | | | | △ | | | | | |
| | | | | | | | ○ | | | | | | | | | | | | | | | | | | | | | | | | | | | | | | | | | △ | | | | | | | | | |
| | | | | | | | | | | | | | | | | | | | | | | | | | | | | | | | | | | | | | | | | | | | | | | | | | ○ |
| | | | | | | | | | | | | | | | | | | ○ | | | | | | | | | | | | | | | | ○ | | ○ | | | | | | | | | | | | | |
| | | | | | ○ | △ | | | | | | | | | | | | | | | | | | | | | | | | | | | | | | | | | ○ △oppo | | | | | | | | | | |
| | | | | | ○ | | | | | | | | | | | | | | | | | | | | | | | △ | | | | | | | | | | | | | | | | | | | | | |
| | | | | | ○ | | | | | | | | | | | | | | | | | | | | | | | | ○ | | | | | | | | | | | | | | | | | | | |
| | | | | | | | | | | | | | | | ○ | | | | | | | | | | | | | | | | | | | | | | | | | | | | | | | | | | |
| | | | | | | | | | | | | | | | | | | | | | | | | | | | | | | | | | | | | | | | | | | | | | | | | ○ | |
| | | | | | | | | | | | | | | | | | | | | | | | | | | | | | | | | | | | | | | | | | | ○ | | | | | | | |
| | | | | | | | | | | | | | | | | | | | | | | | | | | ○ | | | | | | | | | | | | | | | | | | | | | | | |
| | | | | ○ | | | | | | | | | | | | | | | | | | | | | | | △ | | | | | | | | | | | | | | | | | | | | | | |
| | | | | | | | | | | | ○ △oppo | | | | | | | | | | | ○ | | | | | | | | | | | | | | | | | | | | | | | | | | | |
| | | | | | | | | | | | ○ | | | | | | | | | | | | | | | | | | | | | | | | | | | | | | | | | | | | | | |
| | | | | | | | | | | | | | | | | | | | | | | | | | | | △ | | | | | | | △ | | | | | | | | | | | | | | | |
| | | | | | | | | | | | | | | | | | | | | | | | | | | | △ | | | | | | | △ | | | | | | | | | | | | | | | |
| | | ○ | ○ | | | | | | | | △ | ○ | | | | | ○ | | ○ | ○ | | △ | | | | | | | | | | | | | | | | | | | | | | | | ○ | | | |
| | | | | | | | | | | | | | | | | | | | | | | | | | | | | | | | | | | | ○ | | | | | | | | | | | | | | |
| | | | | ○ | | | | | ○ | | | | | | | ○ | | | | | | | | | | | ○ | | | | | ○ | | | | | | | | | | | | | ○ | | | | |
| | | | | | | | | | | | | | | | | | | | | | | | | | | | ○ | | | | | | | | | | | | | | | | | | | | | | |
| | | | | ○ | | | | | ○ | | | | | | | ○ | | | | | | | | | | | ○ | | | | | | | | | | | | | | | | | | ○ | | | | |
| | | | | ○ | | | | | ○ | | | | | | | ○ | | | | | | | | | | | ○ | | | | | | | | | | | | | | | | | | ○ | | | | |
| | | | | | | | | | | | | | | | | | | | | | | | | | | | ○ | | | | | | | | | | | | | | | | | | | | | | |

| #58_Composite_15 0404_11-1 | #58_Composite_15 0404_11-1 | #58_Composite_15 0404_11-1 | #58_Composite_15 0404_11-1 | #58_Composite_15 0404_11-1 | #58_Composite_15 0404_11-1 | #59_Composite_15 0404_12-1 | #59_Composite_15 0404_12-1 | #59_Composite_15 0404_12-1 | #59_Composite_15 0404_12-1 | #59_Composite_15 0404_12-1 | #59_Composite_15 0404_12-1 | #59_Composite_15 0404_12-1 | #59_Composite_15 0404_12-1 | #60_Composite_15 0404_13-1 | #60_Composite_15 0404_13-1 | #60_Composite_15 0404_13-1 | #60_Composite_15 0404_13-1 | #60_Composite_15 0404_13-1 | #60_Composite_15 0404_13-1 | #60_Composite_15 0404_13-1 | #60_Composite_15 0404_13-1 | #60_Composite_15 0404_13-1 | #60_Composite_15 0404_13-1 | #60_Composite_15 0404_13-1 | #61_Composite_15 0404_14-1 | #61_Composite_15 0404_14-1 | #62_Composite_15 0404_15-1 | #62_Composite_15 0404_15-1 | #62_Composite_15 0404_15-1 | #62_Composite_15 0404_15-1 | #62_Composite_15 0404_15-1 | #62_Composite_15 0404_15-1 | #62_Composite_15 0404_15-1 | #62_Composite_15 0404_15-1 | #62_Composite_15 0404_15-1 | #62_Composite_15 0404_15-1 | #63_Composite_15 0404_17-1 | #63_Composite_15 0404_17-1 | #63_Composite_15 0404_17-1 | #63_Composite_15 0404_17-1 | #63_Composite_15 0404_17-1 | #63_Composite_15 0404_17-1 | #63_Composite_15 0404_17-1 | #63_Composite_15 0404_17-1 | #63_Composite_15 0404_17-1 | #64_Composite_15 0404_18-1 | #64_Composite_15 0404_18-1 | #64_Composite_15 0404_18-1 | #64_Composite_15 0404_18-1 |
|---|---|---|---|---|---|---|---|---|---|---|---|---|---|---|---|---|---|---|---|---|---|---|---|---|---|---|---|---|---|---|---|---|---|---|---|---|---|---|---|---|---|---|---|---|---|---|---|---|---|
| | | | | | | 55 | | | | | | | | 56 | | | | | | | | | | | 57 | | 58 | | | | | | | | | | 59 | | | | | | | | | 60 | | | |
| 394 | 395 | 396 | 397 | 398 | 399 | 400 | 401 | 402 | 403 | 404 | 405 | 406 | 407 | 408 | 409 | 410 | 411 | 412 | 413 | 414 | 415 | 416 | 417 | 418 | 419 | 420 | 421 | 422 | 423 | 424 | 425 | 426 | 427 | 428 | 429 | 430 | 431 | 432 | 433 | 434 | 435 | 436 | 437 | 438 | 439 | 440 | 441 | 442 | 443 |
| 7 | 8 | 9 | 10 | 11 | 12 | 1 | 2 | 3 | 4 | 5 | 6 | 7 | 8 | 1 | 2 | 3 | 4 | 5 | 6 | 7 | 8 | 9 | 10 | 11 | 1 | 2 | 1 | 2 | 3 | 4 | 5 | 6 | 7 | 8 | 9 | 10 | 1 | 2 | 3 | 4 | 5 | 6 | 7 | 8 | 9 | 1 | 2 | 3 | 4 |
| R | R | R | R | R | R | L | L | L | L | R | R | R | R | L | L | L | L | L | L | R | R | R | R | R | R | R | L | L | L | L | L | L | L | L | R | R | L | L | L | R | R | R | R | R | R | L | L | R | R |
| | | | | | | | | | | | | | | | | | | | | | | | | | | | | | | △ | | | | | | | | | | | | | | | | | | | |
| | | | | | | | | | | ○ | | | | | | | | | | | | | | | | | | | | ○ | | | | | | | | | | | | | | | | | | | |
| | | | | ○ | | | | | | | | | | | | | | | | | | | | | | | | | | | | | | | | ○ | | | | | | | | | | | | | |
| | | | | | | | | | | | | | | | | | | | | ○ | | | | | | | | | | | | | | | | | | | | | | | | | | | | | |
| | | | | | | | | | | | | | ○ | | | | | | | | | | | | | | | | | | | | | | ○ | | | | | | | ○ | | | | ○ | | | |
| | | | | | | | | | | | | | | | | | | | | | | | | | | | ○ | | | | | | | | | | | | | | | | | | | | | | |
| | | | | ○ | | △ | | | | | | | | | | | | | | | | | | | | | | | | | | | | | | | | | | | | ○ | | | | | | | |
| | | | | ○ | | | | | | | | | | | | | | | | | | | | | | | | | | | | | | | | | | | | | | | | | | | | | |
| | | | | ○ | | △ | | | | | | | △ | △ | | | | | | | | | | | | | △ | △ | | | | | | | △ | | | | | | | △ | | | | △ | | | |
| | | | | ○ | | △ | | | | | | | △ | △ | | | | | | | | | | | | | △ | △ | | | | | | | ○ | | | | | | | △ | | | | △ | | | |
| | | | | | | △ | | | | | | | ○ | | | | | | | | | | | | | | | | | | | | | | | | | | | | | ○ | | | | | | | |
| | | | | | | ○ | | | | | | | | ○ | | | | | | | | | | | | | | ○ | | | | | | | ○ | | | | | | | ○ | | | | ○ | | | |
| | ○ | | | | | | | | | | | | | ○ | | | | | | | | | | | | | | | | | | | | | | | | | | | | | | | | | ○ | | |
| | | | | | | | | | | | ○ | | | | | | | | | | | | | | | | | | △ | | | | | | | | | | | | | | | | | | | | |
| ○ | | | | | | | | | | | | | | | | | | | | | | | | | | | | | | | | | | | | | | | | | | | | ○ | | | | | |
| | | ○ | | | | | | | | | | | | | | | | | | | | | | | | | | | | | | | | | | | | | | | | | | | ○ | | | | |
| | | | | | | | | | | | | | | | | | | | | | | ○ | | | | | | | | | | | | | | | | | | | | | | | | | | | |
| | | | | | | | | | | | | | | | | | | | | | | | | | | | | | | | | ○ | | | | | | | | | | | | | | | | | |
| | | | | | | | | | | | | | | | | | | | | | | | | | | | | | ○ | | | | | | | | | | | | | | | | | | | | |
| | | | | | | | | | | | | | | | | | | | | | | | | | | | | | ○ | | | | | | | | | | | | | | | | | | | | |
| | | | | | | | | | | | | ○ | | | | | | | | | | | | | | | | | | | | | | | | | | | | | | | | | | | | | |
| | | | | | ○ | | | | | | | | | | | | | | ○ | | ○ | | | | | | | | | | | | | | | | | | | | | | | | | | | | |
| | | | | | | | | | | | | | | | | | | | | | | | | ○△oppo | | | | | | | | | | | | | | | | | | | | | | | | | |
| | | | | | | | | | | | | | | △ | | | | | | | | | | | △ | | | | △ | | | | | | | | | | | | | | | | | | | | |
| | | | | | | | | | | | | | | △ | | | | | | | | | | | △ | | | | △ | | | | | | | | | | | | | | | | | | | | |
| | | | | | | | | | | | | | | | ○ | | | | | | | | | | | | | | | | | | | | | | | | | | | | | | | | | | |
| | | | ○ | | | | | | | | | | | | | ○ | | △oppo | | | | | | | ○ | | | | | | | | ○ | | | | | | ○ | | | | ○ | | | | | | |
| | | | | | | | | | | | | | | | | | | ○ | | | | | | | ○ | | | | | | | | | | | | | | | | | | | | | | | | |
| | | | | | | | | | | | | | | | | | | | | | | | | | ○ | | | | | | | | | | | | | | | | | | | | | | | ○ | |
| | | | | | | | | | | | | | | | | | | | | | | | | | ○ | | | | | | | | | | | | | | | | | | | | | | | | |
| | | | | | | | ○ | | | | | | | | | | | | | | | | | | ○ | | | | △ | | ○ | | | | | | ○ | | | ○ | | △ | | | | | △ | | ○ |
| | | | | | | | ○ | | | | | | | | | | | | | | | | | | ○ | | | | | | | | | | | | ○ | | | | | | | | | | | | |
| | | | | | | | ○ | | | | | | | | | | | | | | | | | | | | | | | | | | | | | | ○ | | | ○ | | | | | | | | | ○ |
| | | | | | | | ○ | | | | | | | | | | | | | | | | | | ○ | | | | | | ○ | | | | | | ○ | | | ○ | | | | | | | | | ○ |
| | | | | | | | ○ | | | | | | | | | | | | | | | | | | | | | | | | ○ | | | | | | ○ | | | | | | | | | | | | |

| #64_Composite_15 0404_18-1 | #64_Composite_15 0404_18-1 | #64_Composite_15 0404_18-1 | #64_Composite_15 0404_18-1 | #64_Composite_15 0404_18-1 | #64_Composite_15 0404_18-1 | #65_Composite_15 0404_19-1 | #65_Composite_15 0404_19-1 | #65_Composite_15 0404_19-1 | #65_Composite_15 0404_19-1 | #65_Composite_15 0404_19-1 | #65_Composite_15 0404_19-1 | #65_Composite_15 0404_19-1 | #65_Composite_15 0404_19-1 | #65_Composite_15 0404_19-1 | #65_Composite_15 0404_19-1 | #65_Composite_15 0404_19-1 | #65_Composite_15 0404_19-1 | #65_Composite_15 0404_19-1 | #66_Composite_15 0404_21-1 | #66_Composite_15 0404_21-1 | #66_Composite_15 0404_21-1 | #66_Composite_15 0404_21-1 | #66_Composite_15 0404_21-1 | #66_Composite_15 0404_21-1 | #66_Composite_15 0404_21-1 | #66_Composite_15 0404_21-1 | #67_Composite_15 0404_23-1 | #67_Composite_15 0404_23-1 | #69_Composite_15 0404_27-1 | #69_Composite_15 0404_27-1 | #69_Composite_15 0404_27-1 | #69_Composite_15 0404_27-1 | #69_Composite_15 0404_27-1 | #69_Composite_15 0404_27-1 | #70_141210_05_sti tch_m-1 | #70_141210_05_sti tch_m-1 | #70_141210_05_sti tch_m-1 | #71_141102_02_w rite_merge_2 | #71_141102_02_w rite_merge_2 | #71_141102_02_w rite_merge_2 | #71_141102_02_w rite_merge_2 | #71_141102_02_w rite_merge_2 | #71_141102_02_w rite_merge_2 | #71_141102_02_w rite_merge_2 | #71_141102_02_w rite_merge_2 | #71_141102_02_w rite_merge_2 | #72_141213_03-1 | #72_141213_03-1 | #72_141213_03-1 |
|---|---|---|---|---|---|---|---|---|---|---|---|---|---|---|---|---|---|---|---|---|---|---|---|---|---|---|---|---|---|---|---|---|---|---|---|---|---|---|---|---|---|---|---|---|---|---|---|---|---|
| | | | | | | 61 | | | | | | | | | | | | | 62 | | | | | | | | 63 | | 64 | | | | | | 65 | | | 66 | | | | | | | | | 67 | | |
| 444 | 445 | 446 | 447 | 448 | 449 | 450 | 451 | 452 | 453 | 454 | 455 | 456 | 457 | 458 | 459 | 460 | 461 | 462 | 463 | 464 | 465 | 466 | 467 | 468 | 469 | 470 | 471 | 472 | 473 | 474 | 475 | 476 | 477 | 478 | 479 | 480 | 481 | 482 | 483 | 484 | 485 | 486 | 487 | 488 | 489 | 490 | 491 | 492 | 493 |
| 5 | 6 | 7 | 8 | 9 | 10 | 1 | 2 | 3 | 4 | 5 | 6 | 7 | 8 | 9 | 10 | 11 | 12 | 13 | 1 | 2 | 3 | 4 | 5 | 6 | 7 | 8 | 1 | 2 | 1 | 2 | 3 | 4 | 5 | 6 | 1 | 2 | 3 | 1 | 2 | 3 | 4 | 5 | 6 | 7 | 8 | 9 | 1 | 2 | 3 |
| R | R | R | R | R | R | L | L | L | L | L | L | L | L | R | R | R | R | R | L | L | L | L | L | R | R | R | L | R | L | L | R | R | R | R | L | L | R | L | L | L | L | L | L | R | R | R | L | L | L |
| | | | | | | | | | | | | | | | | | | | | | | | | | | | | | | | | ○ | | | | | △ | | | ○ | | | | | | ○ | | | |
| | | | | | | | | | | | | | | | | | | | | ○ | | | | | | ○ | | | | | | | ○ | | | | △ | | | ○ | | | | | | | | | |
| | | | | | | | | | | | | | | | | | | | | | | | | | | | | | | | | | | | | | ○ | | | | | | | | | | | | |
| | | | | ○ | | ○ | | | | | | | | | | | | | | | | | | | | | | | ○ | | | | | | | | | | | | | | | | | | | | |
| | | | | | | | | | | | | | | | | | ○ | | | | | | | | | | | | | | | | | | | | | | | | | | | | | | | | |
| | | | | | | | | | | | | | | | | | | | | | | | | | | | | | | | | | | | | | | | | | | | | | | | ○ | |
| | | | | | | | | | | | | | | | | | | | | | | | | | | | | | ○ | | △ | | | | | | | | | | | | | | | | | | |
| | | | | | | | | | | | | | | | | | | | | | | | | | | | | | | | △ | | | | | | | | | | | | | | | | | | |
| | | | | | | | | | | | | | | | | | | | | | | | | | | | | | | | △ | | | | | | | | | | | | | | | | | | |
| | | | | | | | | | | | | | | | | | | | | | | | | | | | | | | | ○ | | | | | | | | | | | | | | | | | | |
| | | ○ | | | | | | | | | | | | | | | | | | | | | | | | | | | | | | | | | | | | | | | | | | | | | | | |
| | | | | | | | | | ○ | | | | | | | | | ○ | | | | | | | ○ | | | | | | | | | | | ○ | | | | | | | | | | | | | |
| | ○ | | | | | | | | | ○ | | | | | | | | | ○ | | | | | | | | | | | | | | | | ○ | | △ | | | | ○ | | | | | | | | |
| | | | | | | | | | | | | | | | | | | | | | ○ | | | | | | | | | | | | | | | | | △ | | | | | | | | | | | |
| | | | | | ○ | | | | | | | | | | | | | | | | ○ | | | ○ | | | | | | | | | | | | | | ○ | | | | | | | | | | | |
| | | | | | | | | | | | ○ | | | | | | | | | | | | | | | | | | | | | | | | | | | | | | | | | | | | | | |
| ○ | | | | | | | | | | | | | | | | | | | | | | | | | | | | | | | | | | | | | | | | | | | | | | | | | |
| ○ | | | | | | | | | | | | | | | | | | | | | | | | | | | ○ | | | ○ | | | | | | | | | | | | | | | | | | | |
| | | | | | | | | ○ | | | | | | | | | | | | | | ○ | | | | | | | | | | | | | | | | | | | | | | | | | | | |
| | | | | | | | | | | | | | | | | | | | | | | | | | | | | | | | | | | | | | | | | | | | | | | | | | △ |
| | | | | | | | | | | | | | | | ○△oppo | | | | | | | | | | | | | | | | | | | | | | | | | | | | | | | | | | |
| | | | | | | | | | | | | | | | ○ | | | | | | | | | | | | | | | | | | | | | | | | | | | | | | | | | | |
| | | | ○ | | | | | | | | | | | | △ | ○ | | | | | | | | | | | | | | | | | | | | | | | ○ | | | | | | | | ○ | | |
| | | | | | | | | | | | | ○ | | | | | | | | | | | | | | | | | | | | | | | | | | | | | | | ○ | ○ | | | | | ○ |
| | | | | | | | ○ | | | | | | | ○ | | ○ | △ | | | | | | | △ | | | | | | | | | | | △ | | | △ | | | | | | | ○ | | | △ | ○ |
| | | | | | | | ○ | | | | | | | ○ | | ○ | | | | | | | | △ | | | | | | | | | | | | | | | | | | | | | | | | | ○ |
| | | | | | | | ○ | | | | | | | ○ | | ○ | | | | | | | | | | | | | | | | | | | | | | | | | | | | | | | | | ○ |

| #72_141213_03-1 | #72_141213_03-1 | #72_141213_03-1 | #72_141213_03-1 | #72_141213_03-1 | #72_141213_03-1 | #73_141102_04_w rite_merge_2 | #73_141102_04_w rite_merge_2 | #73_141102_04_w rite_merge_2 | #74_141210_06_sti tch_scale_m | #74_141210_06_sti tch_scale_m | #74_141210_06_sti tch_scale_m | #74_141210_06_sti tch_scale_m | #74_141210_06_sti tch_scale_m | #74_141210_06_sti tch_scale_m | #74_141210_06_sti tch_scale_m | #74_141210_06_sti tch_scale_m | #74_141210_06_sti tch_scale_m | #74_141210_06_sti tch_scale_m | #75_160617_01_Co mposite_cut copy | #75_160617_01_Co mposite_cut copy | #76_160617_05_Co mposite_cut copy | #76_160617_05_Co mposite_cut copy | #77_160617_07_Co mposite_cut copy | #77_160617_07_Co mposite_cut copy | #77_160617_07_Co mposite_cut copy | #77_160617_07_Co mposite_cut copy | #77_160617_07_Co mposite_cut copy | #77_160617_07_Co mposite_cut copy | #77_160617_07_Co mposite_cut copy | #77_160617_07_Co mposite_cut copy | #77_160617_07_Co mposite_cut copy | #77_160617_07_Co mposite_cut copy | #78_160704_10_01 _composite_crop | #78_160704_10_01 _composite_crop | #78_160704_10_01 _composite_crop | #78_160704_10_01 _composite_crop | #78_160704_10_01 _composite_crop | #79_160704_11_01 _composite_crop | #79_160704_11_01 _composite_crop | #79_160704_11_01 _composite_crop | #80_160704_15_01 _composite_crop | #80_160704_15_01 _composite_crop | #81_160704_17_02 _composite_crop_r | #81_160704_17_02 _composite_crop_r | #82_160712_18_l_ composite | #82_160712_18_l_ composite | #82_160712_18_l_ composite | #82_160712_18_l_ composite | #83_160712_20_l_ composite |
|---|---|---|---|---|---|---|---|---|---|---|---|---|---|---|---|---|---|---|---|---|---|---|---|---|---|---|---|---|---|---|---|---|---|---|---|---|---|---|---|---|---|---|---|---|---|---|---|---|---|
| | | | | | | 68 | | | 69 | | | | | | | | | | 70 | | 71 | | 72 | | | | | | | | | | 73 | | | | | 74 | | | 75 | | 76 | | 77 | | | | 78 |
| 494 | 495 | 496 | 497 | 498 | 499 | 500 | 501 | 502 | 503 | 504 | 505 | 506 | 507 | 508 | 509 | 510 | 511 | 512 | 513 | 514 | 515 | 516 | 517 | 518 | 519 | 520 | 521 | 522 | 523 | 524 | 525 | 526 | 527 | 528 | 529 | 530 | 531 | 532 | 533 | 534 | 535 | 536 | 537 | 538 | 539 | 540 | 541 | 542 | 543 |
| 4 | 5 | 6 | 7 | 8 | 9 | 1 | 2 | 3 | 1 | 2 | 3 | 4 | 5 | 6 | 7 | 8 | 9 | 10 | 1 | 2 | 1 | 2 | 1 | 2 | 3 | 4 | 5 | 6 | 7 | 8 | 9 | 10 | 1 | 2 | 3 | 4 | 5 | 1 | 2 | 3 | 1 | 2 | 1 | 2 | 1 | 2 | 3 | 4 | 1 |
| L | R | R | R | R | R | R | R | R | L | L | L | L | L | L | R | R | R | R | R | R | L | R | L | L | L | L | L | R | R | R | R | R | L | L | L | R | R | R | R | R | L | R | R | R | L | L | R | R | L |
| | | | | | | | | | | | | | | | | ○ | | | | | | | | | | | | | | | | | | | | | | | | | | | | | | | | ○ | |
| | | | | | | | | | ○ | | | | | | | | | | | | | | | ○ | | | | | | | | | | | | ○ | | | | | | | | | | | | | |
| | | | | | | | | | | ○ | | | | | | | | | | | | | | ○ | | | | | | | | | | | | ○ | | | ○ | | | | | | | | | | |
| | | | | | ○ | | | | | | | | | | ○ | | | | | | | | | | | | | | | | | | | | | | | | | | | | | | | | | | |
| | | | | | | | | | | | | ○ | | | | | | | | | | | | | ○ | | | | | | | | | | | | ○ | | | | | | | | | | | | |
| | | | | | | | | | | | | ○ | | | | | | | | ○ | ○ | | | | | | | | | | | | | | | | | | | | | | | | | | | | |
| | | | | | | | | | | | | | | | | | | | ○ | | | | | | | | | | | | | | | | | | | | | | | | | | | | | | |
| | | | | | | | | | | | | | | | | | | | ○ | | | | | | | | | | | | | | | | | | | | | | | | | | | | | | |
| △ | ○ | | | | | | | | | | | | | | | | | | | | | | | | | | | | | | | | | | | | | | | | △ | | | | | | | | |
| △ | ○ | | | | | | | | | | | | | | | | | | | | | | | | | | | | | | | | | | | | | | | | △ | | | | | | | | |
| | | | | | | | | | | | | | | | | | | | | | | | | | | | △ | | | | | | | | | | | | | | ○ | | | | | | | | |
| ○ | | | | | | | | | | | | | | | | | | | | | | | | | | | △ | | | | | | | | | | | | | | | △ | | | | | | | |
| | | | | | | | | | | | | | | | | | | | | | | | | | | | | | | | | | | | | | | | | | | | | | | | | | ○ |
| | | | | | | | | | | | | | | | | | | | | | | | | | | | | | | | | | | | | | | | | | | | | | | ○ | | | |
| | | | | | | | | ○ | | | | | | | | | | | | | | | | | | | | | | ○ | | | | | | | | | | ○ | | | | | | | | | |
| | | ○ | | | | | | | | | | | | | | | | | | | | | | | | | | | | | | | | | | | | | | | | | | | | | | | |
| | | | | | | △? | | | | | | | | | | | | | | | | | | | | | | | | | | | | | | | | | | | | ○ | | | | | | | |
| | | | | | | | | | | | | | | | | | | | | | | ○ | | | | | | | | | | | | | | | | | | | | | | | | | | | |
| | | | | ○ | | | | | | | | | ○ | | | | | | | | | | | | | | | | | | | | | | | | | | | | | | | | | | | | |
| | | | | | | | | | | | | | | | | | | | | | | | | | | | | | | | | ○ | | | | | | | | | | | | | | | | | |
| | | | | | | ○△oppo | | | | | | | | | | | | | | | | | | | | | | | | | | | | | ○ | | | | | | | | | | | | | | |
| | | | | | | ○△oppo | | | | | | | | | | | | | | | | | | | | | | | | | | | | | ○ | | | | | | | | | | | | | | |
| | | | | | | | | △ | | | | | | | | | | | | | | | | | | | | | | | | | | | | | | | | | | | | | | | | | |
| | | | | | | | | △ | | | | | | | | | | | | | | | | | | | | | | | | | | | | | | | | | | | | | | | | | |
| | | | | | | △ | | | | | | | | | | | | | | | | | ○ | | | | | | | | ○ | | | | △ | | | | | | | | | ○ | | ○ | | |
| | | | | | | | | | | | | | | ○ | | | | ○ | | | | | | | | | | ○ | | | | | | ○ | | | | | | | | | | ○ | | | | | |
| | | | | | | | | | | | | | | | | | | | | | | | | | | | | | | | | | | ○ | | | | | | | | | | | | | | | |
| | | | ○ | | | | ○ | | | | ○ | | | | | | ○ | | | | | | | | | | ○ | | | | | | ○ | | | | | ○ | | | | | ○ | | | | | | |
| | | | | | | | | | | | | | | | | | | | | | | | | | | | | | | | | | ○ | | | | | | | | | | | | | | | | |
| | | | | | | | ○ | | | | ○ | | | | | | | | | | | | | | | | ○ | | | | | | ○ | | | | | | | | | | ○ | | | | | | |
| | | | | | | | ○ | | | | ○ | | | | | | | | | | | | | | | | ○ | | | | | | ○ | | | | | | | | | | ○ | | | | | | |
| | | | | | | | | | | | | | | | | | | | | | | | | | | | | | | | | | ○ | | | | | | | | | | | | | | | | |

| #83_160712_20_l_composite | #83_160712_20_l_composite | #84_160712_22_l_composite | #84_160712_22_l_composite | #85_160712_25_l_composite | #85_160712_25_l_composite | #85_160712_25_l_composite | #85_160712_25_l_composite | #85_160712_25_l_composite | #85_160712_25_l_composite | #85_160712_25_l_composite | #85_160712_25_l_composite | #86_160712_26_l_Composite | #86_160712_26_l_Composite | #86_160712_26_l_Composite | #86_160712_26_l_Composite | #86_160712_26_l_Composite | #86_160712_26_l_Composite | total | |
|---|---|---|---|---|---|---|---|---|---|---|---|---|---|---|---|---|---|---|---|
| | | 79 | | 80 | | | | | | | | 81 | | | | | | | |
| 544 | 545 | 546 | 547 | 548 | 549 | 550 | 551 | 552 | 553 | 554 | 555 | 556 | 557 | 558 | 559 | 560 | 561 | 563 | |
| 2 | 3 | 1 | 2 | 1 | 2 | 3 | 4 | 5 | 6 | 7 | 8 | 1 | 2 | 3 | 4 | 5 | 6 | | |
| L | R | L | R | R | R | R | R | L | L | L | L | L | L | L | L | R | R | | |
| | | | | | | | | | | ○ | | | ○ | | | | | 29 | Dla dorsal (Dla_d) |
| | | | | | | | ○ | | | ○ | | | | ○ | | | ○ | 25 | Dla ventral (Dla_v) |
| | | | | | | | | | | | | | | | | | | 21 | Dld anterior (Dld_a) |
| | | | | | | | | | | | | | | | | | | 18 | Dld lateral to Dd2a (Dld_ma) |
| | | | | | | | | | | | | | | | | | | 21 | Dld leteral to Dd2p (Dld_m) |
| | | | | | | | | | | | | | | | | | | 32 | Dld posterior to Dd2 (Dld_md) |
| | | | | | | | | | | | | | | | | | | 5 | Dld posterir (Dld_d) |
| | | | | | | | | | | | | | | | | | | 4 | Dlp |
| | | | | | | | | | | | | | | | | | | 7 | Dlv anterior dorsal (Dlv_ad) |
| | | | | | | | | | | | | | | | | | | 7 | Dlv anterior ventral (Dlv_av) |
| | | | | | | | | | | | | | | | | ○ | | 18 | Dlv medial dorsal (Dlv_md) |
| | | | | | | | | | | | | | | | | | | 13 | Dlv medial ventral (Dlv_mv) |
| | | | | | | | | | | | | | | | | | | 7 | Dlv posterior dorsal (Dlv_pd) |
| | | | | | | ○ | | | | | | | | | | | | 12 | Dlv poesterior ventral (Dlv_pv) |
| | | | | | | | | | | | | | | | | | | 25 | Dd1 lateral |
| | | | | | | | | | | | | | | | | | | 11 | Dd1 medial |
| | | | | | | | | | | | | | | | | | | 49 | Dd2a lateral (Dd2a_l) |
| | | | | | | | | | | | ○ | | | | | | | 53 | Dd2a medial (Dd2a_m) |
| | | | | | | | | | | | | | | | | | | 29 | Dd2p lateral (Dd2p_l) |
| | | | | | | | | | | | | | | | | | | 41 | Dd2p medial (Dd2p_m) |
| | | | | | | | | | | | | | | | | | | 18 | Dm1 dorsal (Dm1_d) |
| | | | | | | | | | | | | | | | | | | 7 | Dm1 medial (Dm1_m) |
| | | | | | | | | | | | | | | | | | | 19 | Dm1 anterior ventral (Dm1_av1) |
| | | | | | | | | | | | | | | | | | | 19 | Dm2a anterior (Dm2a_a) |
| | | | | | | | | | | | | | | | | | | 17 | Dm2a medial (Dm2a_m) |
| | | | | | | | | ○ | | | | | | | | | | 16 | Dm2a posterior (Dm2a_p) |
| ○ | | | | | | | | | | | | | | | | | | 4 | Dm2b anterior (Dm2b_a) |
| | | | | | | | | | | | | | | | | | | 5 | Dm2b medial (Dm2b_m) |
| | ○ | | | | | | | | ○ | | | | | | | | | 4 | Dm2b posterior (Dm2b_p) |
| | | | | | | | | | | | | | | | | | | 3 | Dm3a antrior (Dm3a_a) |
| | | | | | | | | | | | | | | | | | | 3 | Dm3a medial (Dm3a_m) |
| | | | | | | | | | | | | | | | | | | 5 | Dm3a posterior (Dm3a_p) |
| | | | | | | | | | | | | | | | | | | 1 | Dm3b antrior (Dm3b_a) |
| | | | | | | | | | | | | | | | | | | 0 | Dm3b medial (Dm3b_m) |
| | | | | | ○ | | | | | | | | | | | | | 12 | Dm3b posterior (Dm3b_p) |
| | | | | | | | | | | | | | | | | | | 26 | Dm4 |
| | | | | ○△opp | | | | | | | | | | | | | | 21 | Dp anteriror (Dp_a) |
| | | | | ○ | | | | | | | | | | | | | | 14 | Dp posterior (Dp0_p ) |
| | | | | | | | | | | | | | | | | | | 18 | Dc3 |
| | | | | | | | | | | | | | | | | | | 11 | Dc4 |
| | | | | | | | | | | | | | | | | | | 9 | Dcpm |
| | | | | △ | | △ | | | | | | ○ | | | | | | 63 | GL |
| | | | | | | | | | | | | | | | | | | 13 | ICL |
| | | ○ | ○ | | | | | | | | | | | | | | | 16 | Vv |
| | | | | | | | | | | | | | | | | | | 2 | Vv inside (Vv_peri) |
| | | | | | | | | | | | | | | | | | | 1 | Vc |
| | | | | | | | | | | | | | | | ○ | | | 76 | Vd |
| | | | | | | | | | | | | | | | | | | 11 | Vd inside (Vd_peri) |
| | | | | | | | | | | | | | | | | | | 40 | Vp |
| | | | | | | | | | | | | | | | | | | 45 | Vs |
| | | | | | | | | | | | | | | | | | | 9 | Vs inside (Vs_peri) |
| | | | | | | | | | | | | | | | | | | 3 | Vl |
